# Evaluation of PRKACA as a Therapeutic Target for Fibrolamellar Carcinoma

**DOI:** 10.1101/2022.01.31.477690

**Authors:** Stefanie S. Schalm, Erin O’Hearn, Kevin Wilson, Timothy P. LaBranche, Grace Silva, Zhuo Zhang, Lucian DiPietro, Neil Bifulco, Richard Woessner, Nicolas Stransky, Darshan Sappal, Robert Campbell, Riadh Lobbardi, Michael Palmer, Joseph Kim, Chaoyang Ye, Marion Dorsch, Christoph Lengauer, Timothy Guzi, Vivek Kadambi, Andrew Garner, Klaus P. Hoeflich

**Affiliations:** Blueprint Medicines Corporation, Cambridge, Massachusetts

**Keywords:** fibrolamellar carcinoma, PRKACA, DNAJB1-PRKACA, kinase inhibitor

## Abstract

**Background & Aims:** Fibrolamellar carcinoma (FLC) is a rare, difficult-to-treat liver cancer primarily affecting pediatric and adolescent patients, and for which precision medicine approaches have historically not been possible. The *DNAJB1-PRKACA* gene fusion was identified as a driver of FLC pathogenesis. We aimed to assess whether FLC tumors maintain dependency on this gene fusion and determine if PRKACA is a viable therapeutic target.

**Methods:** FLC patient-derived xenograft (PDX) shRNA cell lines were implanted subcutaneously into female NOD-SCID mice and tumors were allowed to develop prior to randomization to doxycycline (to induce knockdown) or control groups. Tumor development was assessed every 2 days. To assess the effect of treatment with novel selective PRKACA small molecule kinase inhibitors, BLU0588 and BLU2864, FLC PDX tumor cells were implanted subcutaneously into NOD-SCID mice and tumors allowed to develop. Mice were randomized to treatment (BLU0588 and BLU2864, orally, once daily) or control groups and tumor size determined as above.

**Results:** Knockdown of *DNAJB1-PRKACA* reversed a FLC-specific gene signature and reduced PDX tumor growth in mice compared to the control group. Furthermore, FLC PDX tumor growth was significantly reduced with BLU0588 and BLU2864 treatment versus control (*P* = 0.003 and *P* = 0.0005, respectively).

**Conclusions:** We demonstrated, using an inducible knockdown and small molecule approaches, that FLC PDX tumors were dependent upon *DNAJB1*-*PRKACA* fusion activity. In addition, this study serves as a proof-of-concept that PRKACA is a viable therapeutic target for FLC and warrants further investigation.

## Introduction

Fibrolamellar carcinoma (FLC) represents <1% of all liver cancers in the United States (1,2). FLC primarily affects pediatric or adolescent patients, with the majority of patients being under 40 years old (2). Surgery is the mainstay treatment for FLC; there has been no clear benefit demonstrated with any chemotherapy or multi-kinase inhibitors, including sorafenib, which is the standard treatment for hepatocellular carcinoma (HCC) (3). FLC is often not diagnosed until the disease has metastasized, which complicates surgical resection (4,5). Most patients experience disease recurrence within 5 years (6,7), and the 5-year survival rate is approximately 75%– 85% for resected patients (7,8). New therapies are needed; however, until recently, precision medicine approaches have not been possible due to a lack of an identified driver of FLC pathogenesis.

FLC is clinically, histologically, and molecularly distinct from the most common forms of liver cancer, including HCC and cholangiocellular carcinoma (CCC). FLC is identified histologically by large tumor cells arranged in a cord-like network surrounded by lamellated collagen fibers, and is not typically associated with elevated α-fetoprotein (9). FLC can be characterized by positive immunohistochemical staining for markers of both hepatocyte and biliary differentiation including cytokeratin 7 (10) and CD68 (11), yet is often negative for the HCC marker glypican 3 (12). An FLC-specific gene expression signature has been identified that can further distinguish it from healthy liver, HCC, and CCC (13,14).

In 2014, an analysis of 15 FLC tumor samples yielded the discovery of an aberrant fusion between protein kinase A (*PKA*) alpha catalytic subunit (*PRKACA*) and heat shock protein 40 (*DNAJB1*) genes that was hypothesized to be a critical molecular event in FLC pathogenesis (15). The *DNAJB1-PRKACA* fusion is not expressed by normal surrounding liver tissue or other liver cancer types, but has been identified in 79%–100% of FLC tumor samples (15,16) and is functionally capable of driving FLC tumorigenesis when expressed in mice (17-20), making its presence a distinguishing hallmark of FLC and a likely driver of FLC tumorigenesis (15,21).

The 3′ 5′-cyclic adenosine monophosphate (cAMP)-regulated PKA tetramer is comprised of 2 catalytic subunits and 2 regulatory subunits. In the presence of cAMP, the catalytic subunits disassociate from regulatory subunits and become active. PKA regulates a broad array of cellular functions throughout the body (22). Through its regulation of transcriptional elements, the PKA signaling pathway is involved in liver metabolism (23). The chimeric kinase resulting from the *DNAJB1-PRKACA* fusion retains the kinase activity of PRKACA (15,24). The DNAJB1-PRKACA fusion protein is expressed at a higher level than wild-type (WT) PRKACA (15,25), and it has been postulated that these higher levels lead to increased cAMP-independent PKA signaling (26). The molecular mechanism of how the *DNAJB1-PRKACA* fusion contributes to FLC pathogenesis remains incompletely understood, and it is still not fully clear whether FLC tumors maintain dependency on DNAJB1-PRKACA expression for their continued growth, and whether PRKACA is a viable therapeutic target.

In this study, we demonstrate that 2 different modes of PRKACA inhibition result in FLC tumor growth inhibition and reversal of an FLC gene expression signature, using a patient-derived xenograft (PDX) FLC mouse model (27). This study serves as a proof-of-concept for PRKACA as a viable target for FLC.

## Results

### Characterization of FLC Patient-Derived Xenograft Mouse Model

The FLC PDX tumor model LI5132 expressed high levels of DNAJB1-PRKACA fusion protein and, to a lesser extent, the smaller WT PRKACA protein, distinguishable by their size difference via gel electrophoresis (**Figure 1a**). By contrast, Hep3B cell line-derived xenograft tumors expressed only WT PRKACA protein (**Figure 1a**). The high expression of PRKACA in FLC PDX tumor tissue is likely due to the replacement of exon 1 of *PRKACA* with exon 1 of *DNAJB1* containing the more active *DNAJB1* promoter (15) and upregulating PRKACA protein expression (**Supplementary Figure S1a, S1b**) (29). Phosphorylation of the PRKACA downstream targets, cAMP-responsive binding protein (CREB) and vasodilator-stimulated phosphoprotein (VASP), were measured to assess the cellular activity of DNAJB1-PRKACA fusion protein. Both CREB (Ser133) and VASP (Ser157) phosphorylation was increased in FLC PDX tumor tissue compared with Hep3B cell line derived xenograft (CDX) tumor tissue, supporting enhanced activation of PRKACA signaling from DNAJB1-PRKACA fusion expression (**Figure 1a**). Importantly, PDX tumors recapitulated the fibrolamellar histology that is observed in FLC, characterized by large cells with abundant cytoplasm, positive expression of cytokeratin 7 (10) and CD68 (11), and negative expression of glypican 3 (12) (**Figure 1b**).

**Figure 1.**
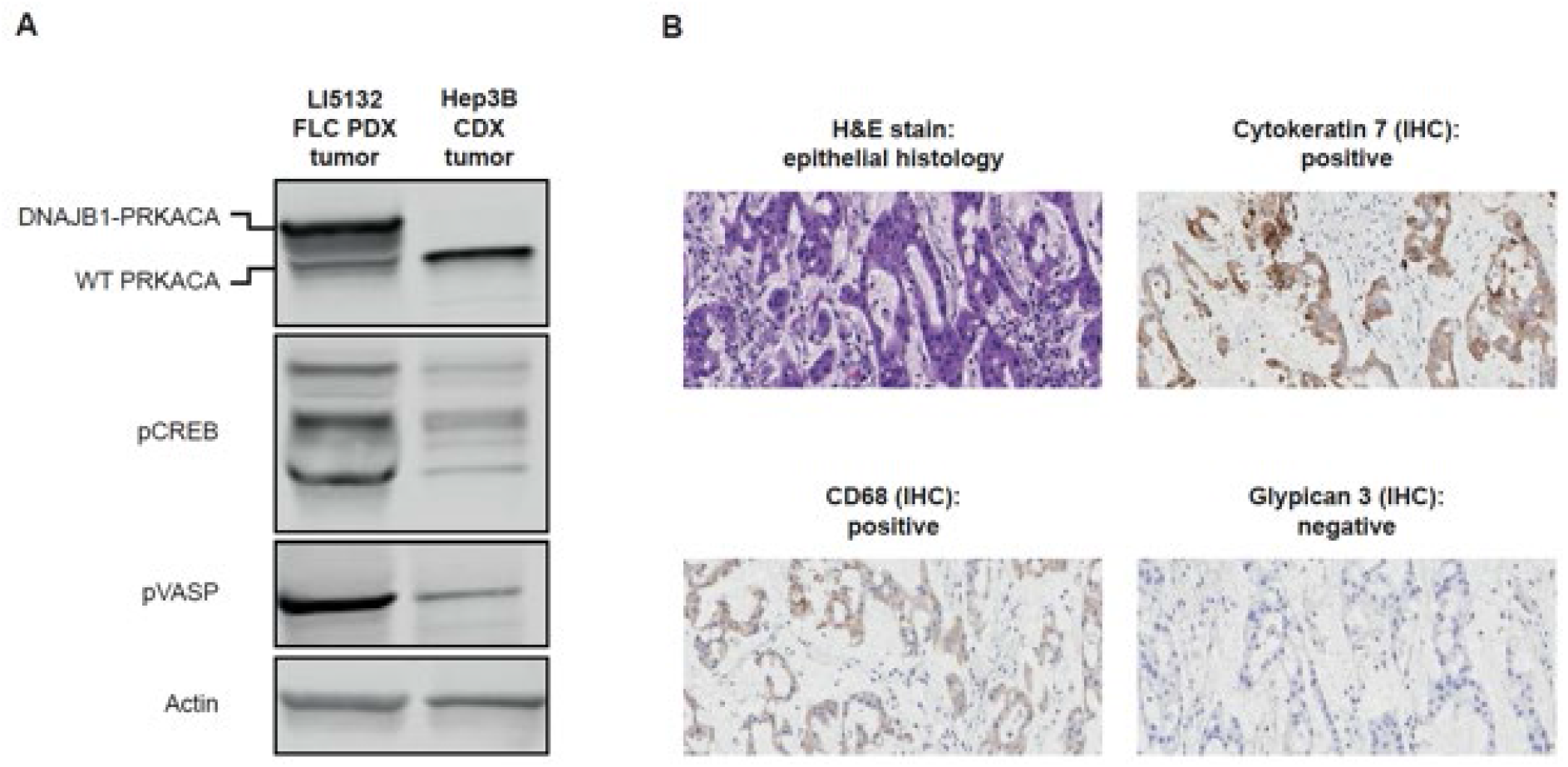
Characterization of an FLC PDX mouse model. (A) Expression of DNAJB1-PRKACA protein in extracts from FLC PDX tumors was associated with increased pCREB and pVASP. (B) Hematoxylin and eosin staining of sections from FLC PDX tumors shows a fibrolamellar morphology, and immunohistochemical staining shows positive expression of cytokeratin 7 and CD68, and negative expression of glypican 3 (20X magnification shown).

### Cellular Effects of the DNAJB1-PRKACA Fusion Protein in FLC PDX Spheroid Cultures

Western blot analysis of in vitro cultured engineered cell lines (**Figure 2a**) confirmed that doxycycline induction of the 3 PRKACA short hairpin ribonucleic acid (shRNA) constructs, but not the non-targeting (NT) shRNA construct, led to a reduction in both DNAJB1-PRKACA and WT PRKACA protein (**Figure 2b**).

**Figure 2.**
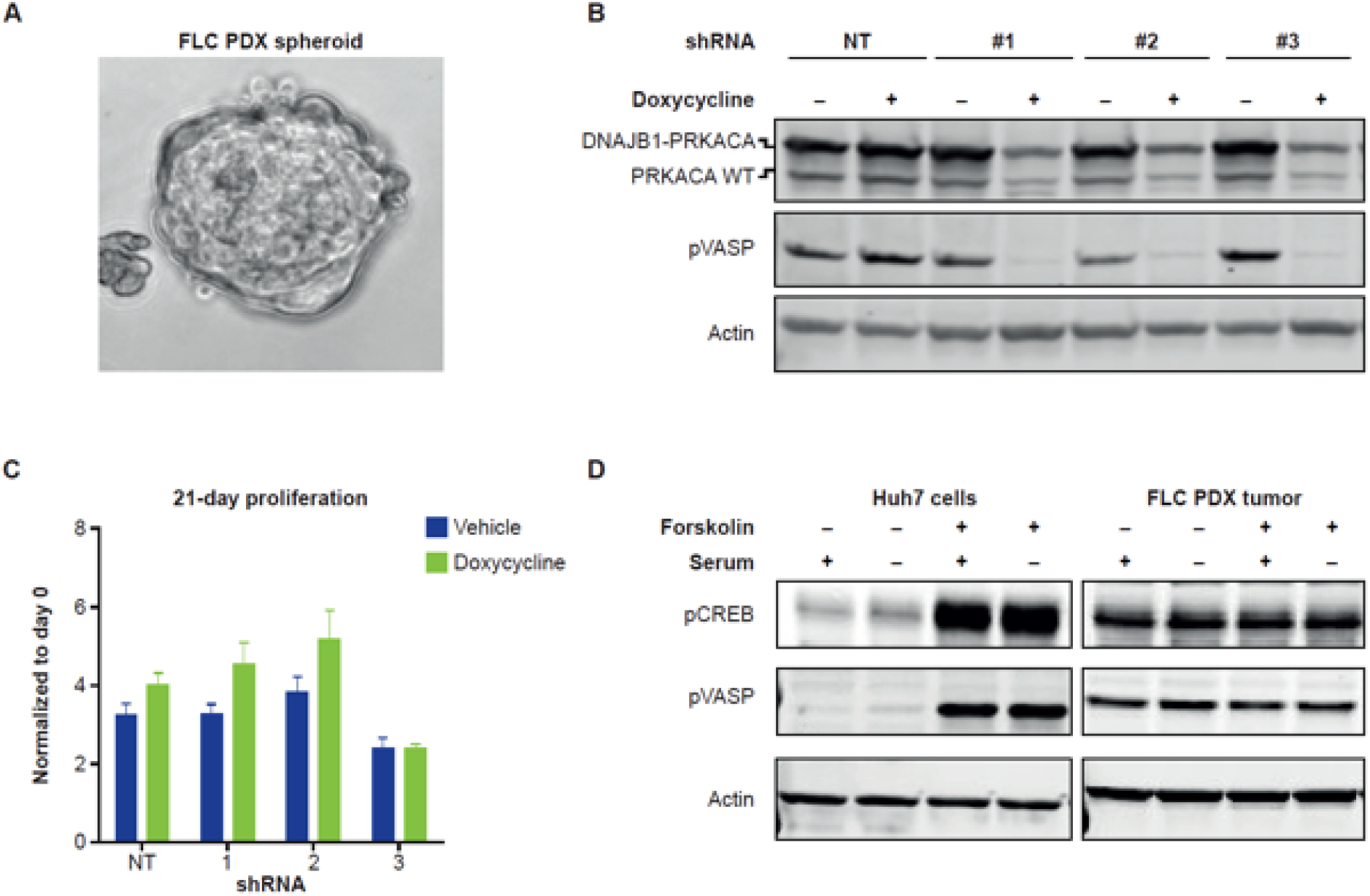
Effect of DNAJB1-PRKACA knockdown on FLC PDX spheroid cultures in vitro. (A) Spheroid cultures were generated from FLC PDX tumors (10X magnification shown). (B) Doxycycline treatment leads to DNAJB1-PRKACA and WT PRKACA knockdown in 3 clones expressing doxycycline-inducible shRNA but does not affect DNAJB1-PRKACA levels in cells expressing a non-targeting shRNA (3 independent biological experiments). (C) Doxycycline treatment does not affect proliferation of PRKACA shRNA-expressing clones or of cells expressing a non-targeting shRNA. Graph Bars and error bars represent the mean ± SD (7 technical replicates and 2 biological replicates). (D) Addition of forskolin (10 μM) increases pCREB and pVASP in extracts from Huh7 HCC cells and has no effect on pCREB or pVASP in extracts from FLC PDX tumors (2 independent biological replicates conducted on different days).

Doxycycline induction of the PRKACA shRNAs, but not the NT shRNA, also strongly reduced VASP phosphorylation, indicative of reduced PRKACA downstream signaling (**Figure 2b**). Cell proliferation was not appreciably different in PRKACA shRNA cell lines before and after induction (**Figure 2c**). Treatment with forskolin did not further enhance CREB and VASP phosphorylation in the FLC PDX tumor extracts (**Figure 2d**), suggesting PRKACA signaling is already fully activated by DNAJB1-PRKACA. By contrast, forskolin produced a strong effect on PRKACA signaling in Huh7 HCC cell extracts, which have a low basal level of unstimulated PRKACA activity (**Figure 2d**).

### Knockdown of DNAJB1-PRKACA Fusion Protein Inhibits Tumor Growth In Vivo

In mice harboring 1 of the 3 different PRKACA shRNA-expressing cell line xenografts (shRNAs #1, #2, and #4) treated with doxycycline, tumor growth was reduced over time compared with mice that were untreated (**Figure 3a-3c**).

**Figure 3.**
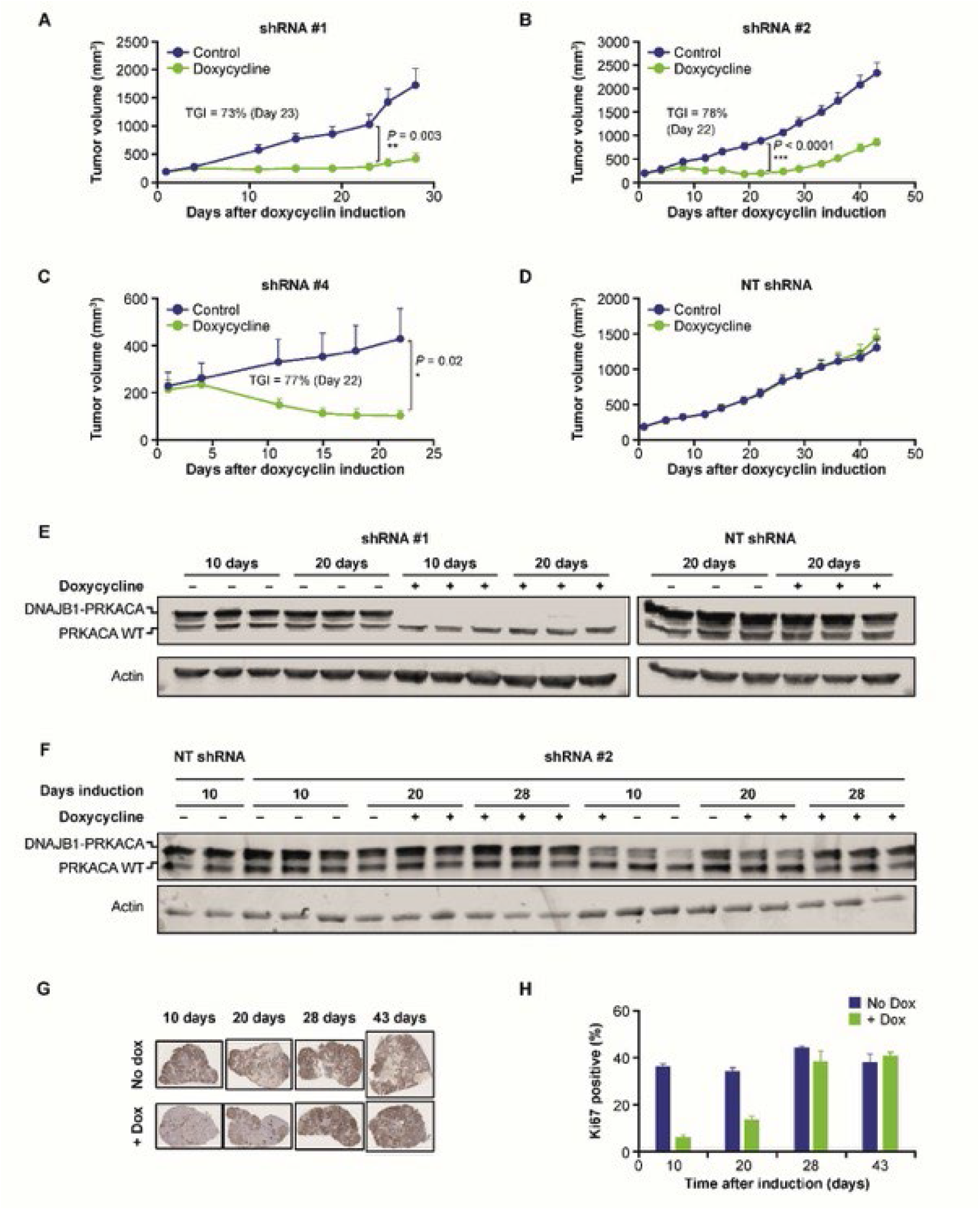
Effect of DNAJB1-PRKACA knockdown on PDX tumor growth in vivo. (A–C), Reduction in the growth rate of PDX tumors expressing doxycycline-inducible PRKACA shRNA was evident in mice treated with doxycycline compared with those that were not. Dots and error bars represent the mean ± SEM ([A] 6 animals per group; [B] 8 animals per group and the study was run twice with comparable results; [C] 6 animals in control group and 7 animals in doxycycline group). (D) No change in the growth rate of PDX tumors expressing doxycycline-inducible non-targeting shRNA was evident in mice treated with doxycycline compared with those that were not. Dots and error bars represent the mean ± SEM (6 animals per group and study was run twice with comparable results). (E) Expression of DNAJB1-PRKACA fusion protein, but not WT PRKACA, was reduced in vivo using shRNA #1 at 10 and 20 days. Non-targeting (NT) shRNA had no effect on fusion protein or WT protein expression (western blot of cell lysate n = 1; lysate from 3 animals per condition). (F) Measurement of DNAJB1-PRKACA and WT PRKACA expression in extracts from FLC PDX tumors expressing clone #2 indicatedDNAJB1-PRKACA expression recovers by 28 days after doxycycline treatment of mice (3 animals per condition for shRNA induction; 3 independent western blots of tumor lysate). (G) Immunohistochemical staining (4X magnification shown) shows the reduction in Ki67-positive nuclei in FLC PDX tumors within 10 days following doxycycline induction, and recovery to baseline levels approximately 28 days after doxycycline induction. Tumors from 3 mice per group were processed and stained for Ki67. (H) Ki67 percentage positive. Bar graphs and error represent mean ± SEM (3 xenograft tumors per group).

Importantly, tumor growth was unaffected by doxycycline treatment in mice harboring xenografts with NT shRNA (**Figure 3d**). Western blot analysis of tumor tissue from mice implanted with the shRNA #1 cell line indicated that expression of DNAJB1-PRKACA fusion protein, but not WT PRKACA, was reduced at 10 and 20 days of doxycycline treatment (**Figure 3e**), consistent with the observed reduction in tumor growth (**Figure 3a**). Induction of NT shRNA did not affect the expression levels of DNAJB1-PRKACA or WT PRKACA (**Figure 3e**). WT PRKACA expression appeared stable in the presence of doxycycline in the shRNA #1 cell line (**Figure 3e**), potentially due to endogenous expression of PRKACA from non-transfected mouse stromal cells.

Western blot analysis of shRNA #2 tumors indicated DNAJB1-PRKACA expression was also reduced 10 days after doxycycline treatment but recovered over time, with DNAJB1-PRKACA expression reaching baseline levels by approximately 28 days after doxycycline treatment (**Figure 3f**). Similar to the DNAJB1-PRKACA fusion expression levels, a slight recovery in the growth rate of xenograft tumors in mice treated with doxycycline is evident over time and could be due to the recovery of DNAJB1-PRKACA fusion protein levels via outgrowth of clonal populations with weaker DNAJB1-PRKACA knockdown (**Figure 3b**). In further support of the growth-suppressive role of DNAJB1-PRKACA knockdown, the percentage of cells positive for the cellular proliferation marker Ki67 was reduced 10 days after doxycycline treatment in PDX tumors expressing PRKACA shRNA #2 (**Figure 3g** and **3h**). In line with DNAJB1-PRKACA fusion protein expression recovering approximately 28 days following doxycycline treatment, Ki67 expression also recovered to near-baseline levels by 28 days post-induction (**Figure 3g, 3h**).

### The PRKACA-Selective Inhibitors BLU0588 and BLU2864 Inhibit PRKACA Signaling In Vitro and Reduce FLC Tumor Growth In Vivo

A library of over 10,000 chemically diverse kinase inhibitors annotated against the human kinome was interrogated to identify compounds with inhibitory activity against PRKACA. Iterative medicinal chemistry optimization was performed from these initial compounds to improve PRKACA potency, selectivity against related AGC-family kinases, and pharmaceutical properties, leading to the generation of 2 structurally distinct inhibitors, BLU0588 and BLU2864 (**Figure 4a**). Kinome-wide selectivity was assessed by testing BLU0588 and BLU2864 at 3 μM concentration across a panel of 400 human kinases using the KINOME*scan®* methodology (30). This profiling revealed that BLU0588 and BLU2864 have good to moderate selectivity against closely related AGC kinase family members, and excellent overall kinome selectivity profiles, with [S(10)] selectivity scores (30) of 0.047 and 0.057, respectively (**Supplementary Table S1**). As follow up, the dissociation constant (Kd) was determined for all non-mutant kinases that were bound by either BLU0588 or BLU2864 with >90% occupancy at the 3-μM screening concentration. Measurement of the Kd demonstrated that both molecules had the most potent binding affinity to PRKACA, 4 nM, and 3.3 nM, respectively. BLU0588 and BLU2864 exhibited a Kd value <100 nM for only 9 or 10 of the non-mutant kinases identified in the kinome screening assay (**Supplementary Table S2**).

**Figure 4.**
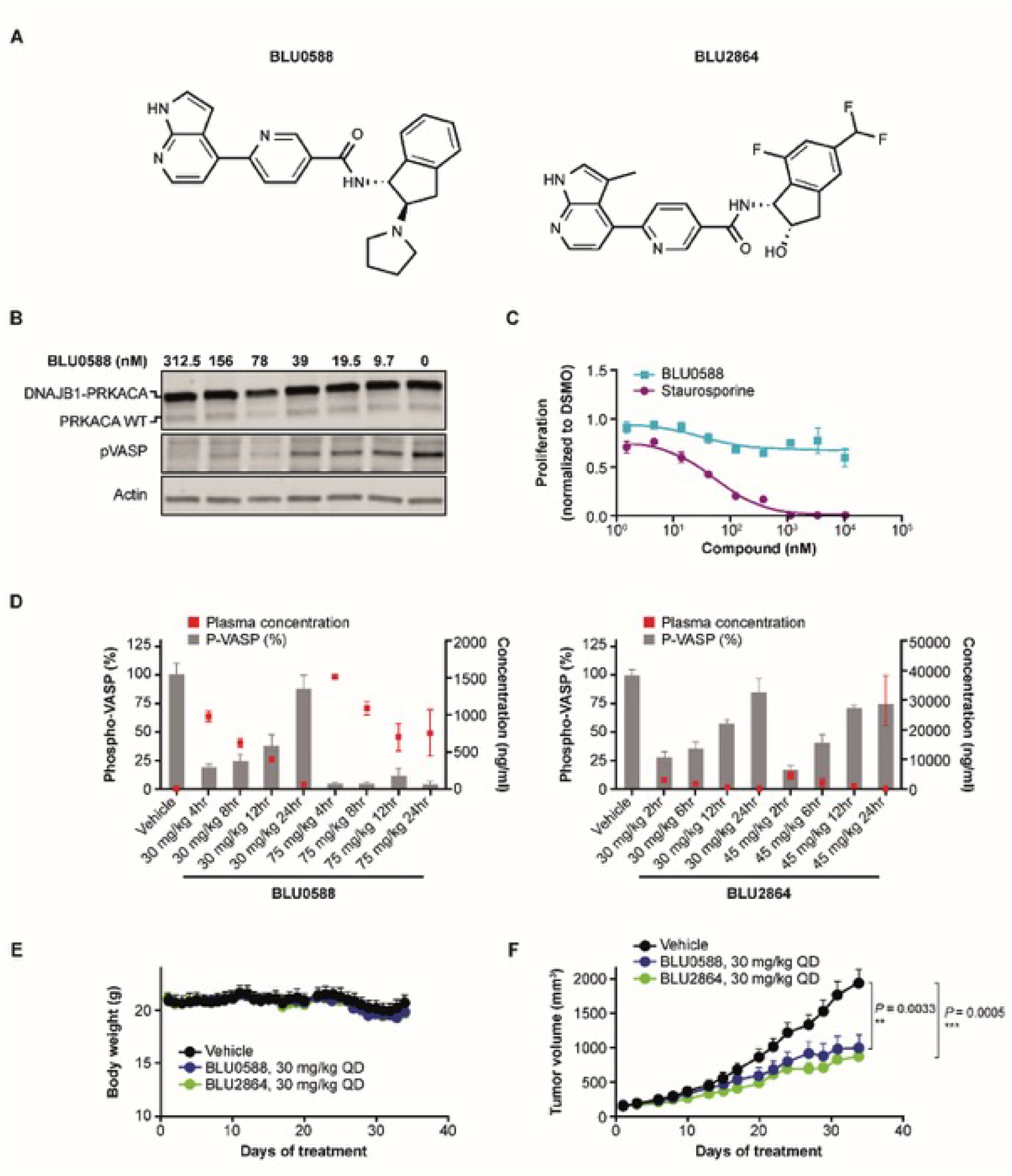
Effect of BLU0588 and BLU2864 on FLC PDX cells in vitro and on FLC PDX growth in vitro. (A) Structure of BLU0588 and BLU2864. (B) Treatment of FLC PDX cells with BLU0588 reduced pVASP in a dose-dependent manner. Experiment was independently repeated 3 times. (C) BLU0588 did not affect FLC PDX cell proliferation after 14 days compared with staurosporine treatment. Squares/dots and error bars represent mean ± SEM (1 biological replicate and 6 technical replicates). (D) The plasma concentration of BLU0588 and BLU2864 (right axis, dots) is shown with the level of pVASP (left axis, bars) after treatment with 30 mg/kg QD and 45 mg/kg QD (BLU2864) or 75 mg/kg QD (BLU0588) orally. Graph bars/dots and error bars represent mean ± SEM (PKPD analysis was conducted once with 3 animals per group [BLU0588] and twice with 3 animals per group [BLU864]). (E) Mouse body weight over time after treatment with 30 mg/kg QD BLU0588 and BLU2864. Dots and error bars represent mean ± SEM (study was conducted once with 9 animals per group [BLU0588] and once with 8 animals per group [BLU2864]). (F) Growth of FLC PDX tumors was reduced in mice treated with 30 mg/kg QD BLU0588 and BLU2864 compared with those treated with vehicle. Dots and error bars represent mean ± (study was conducted once with 9 animals per group [BLU0588] and once with 8 animals per group [BLU2864]).

BLU0588 and BLU2864 inhibited PRKACA catalytic activity with a half-maximal inhibitory concentration (IC50) of 1 nM and 0.3 nM, respectively, compared to an IC50 of 83.1 nM (83-fold) and 12.7 nM (42-fold) for the closely related AGC kinase ROCK2 (**Supplementary Table S1**). For BLU0588, AKT1, AKT2, and AKT3 (AGC kinases that are also closely related to PRKACA), IC50 values were 1540 nM, 3780 nM, and 397 nM, respectively; corresponding values for BLU2864 were 2120 nM, 4910 nM, and 475 nM (**Supplementary Table S1**). PRKACA cellular IC50 values were 25.0 nM and 36.6 nM with BLU0588 and BLU2864, respectively, which were determined from inhibition of VASP Ser157 phosphorylation in forskolin-stimulated Huh7 cells (**Supplementary Table S1; Figure 4b**). These results show BLU0588 and BLU2864 exhibited a strong degree of selectivity for PRKACA.

BLU0588 treatment led to a dose-dependent reduction in phosphorylated VASP in FLC PDX spheroid cultures, indicative of inhibited PRKACA signaling in the disease model (**Figure 4b** and **Supplementary Figure S2**). Similar to the effect of DNAJB1-PRKACA knockdown, proliferation was not substantially inhibited in by BLU0588 treatment in the FLC PDX spheroid model (**Figure 4c**).

For in vivo studies, mice were given either BLU0588 at 30 mg/kg and 75 mg/kg once daily (QD) or BLU2864 at 30 mg/kg and 45 mg/kg QD and monitored for 24 hours. Plasma concentrations peaked within 2–4 hours of QD dosing (**Figure 4d**). For BLU0588-treated mice, phosphorylated VASP was reduced to 19% and 4% of baseline phosphorylation levels with 30 mg/kg QD and 75 mg/kg QD, respectively, 4 hours after dosing; phosphorylated VASP levels fully recovered by 24 hours post-administration of 30 mg/kg QD. BLU2684 at 30 mg/kg QD reduced phosphorylated VASP levels to 27% of baseline 2 hours after dosing, which recovered by 24 hours post-administration of BLU2864 30 mg/kg and 45 mg/kg QD. These data indicate a single oral dose of BLU0588 and BLU2864 can inhibit PRKACA in vivo (**Figure 4d**). The highest tolerated dose of BLU0588 and of BLU2864 for more than 3 weeks of continuous dosing was established as 30 mg/kg QD in mice (**Figure 4e**). When mice harboring FLC PDX tumors were treated with BLU0588 or BLU2864 both given orally at 30 mg/kg QD, by Day 34 tumor growth was inhibited by 48.5% (*P* = 0.003) and by 45.3% (*P* = 0.0005), respectively (**Figure 4f**). These data suggest FLC PDX tumor growth is dependent on PRKACA catalytic activity.

### BLU0588 and PRKACA shRNA Define a Gene Signature of PRKACA Inhibition in FLC

To determine an FLC gene signature that depends on PRKACA expression, we performed RNA sequencing of 3 FLC PDX cell lines expressing PRKACA shRNA (shRNA #1, #2, and #3) and the NT shRNA control. Less than approximately 1% of the total reads per sample aligned to the mouse genome (GENCODE GRCm38/mm10), confirming that purification of human tumor cells was successful. The small number of aligned mouse reads were removed prior to expression quantification using Salmon (31). The majority (81%–97%) of PRKACA reads per sample were identified as the *DNAJB1-PRKACA* fusion and were not *PRKACA* WT (**Supplementary Table S3**), further supporting that the *DNAJB1-PRKACA* fusion is overexpressed and driving the FLC phenotype (**Supplementary Figure S1b**). A significant reduction in PRKACA transcript level (Wilcoxon Signed Rank test *P* = 0.022) was observed with shRNA knockdown (**Supplementary Figure S3**). We detected 572 protein-coding genes that are differentially expressed in all 3 PRKACA shRNA PDX cell lines vs the NT control line (242 negative fold change, **Supplementary Table S4a**; 330 positive fold change, **Supplementary Table S4b** and **Supplementary Figure S4**). Of those, 90 genes are shared between our experiment and FLC tissue vs normal adjacent tissue as described in Simon et al (14) (**Supplementary Figure S4** and **Supplementary Tables S4a and S4b**).

We detected 817 protein-coding genes that were differentially expressed in BLU0588 vs dimethyl sulfoxide (DMSO)-treated FLC PDX cells (323 negative fold change and 494 positive fold change; **Supplementary Tables S4a** and **S4b**, respectively). By matching the inverse log2 fold change across the list of BLU0588-modulated genes and the list of FLC-related genes previously reported by Simon et al (14), we identified a significant overlap between these 2 gene sets (n = 175; Fisher exact test *P* = 0.0005; **Supplementary Tables S5a** and **S5b**). Importantly, the list of BLU0588-modulated genes and the list of PRKACA shRNA-modulated genes showed a significant overlap of differentially expressed genes, defining a signature of PRKACA inhibition (N = 206; Fisher exact test *P* < 2.2×10-16), and suggests that the kinase activity of the DNAJB1-PRKACA fusion protein drives FLC-specific gene expression (**Figure 5a; Supplementary Figures S5a** and **S5b**). Next, we compared the list of genes affected by PRKACA inhibition identified above to the FLC gene signature identified by Simon et al (14) to determine those that can be reversed by PRKACA inhibition. By matching the inverse log2-fold change across both gene lists, we identified a core set of 39 FLC genes that are modulated by both BLU0588 treatment and shRNA knockdown and have an inverse modulation in FLC (**Figure 5b** and **Supplementary Figure S5b, Supplementary Tables S6a** and **S6b**). These genes include FLC markers such as carbamoyl phosphate synthetase 1 (CPS1), KRT86, and sonic hedgehog (SHH).

**Figure 5.**
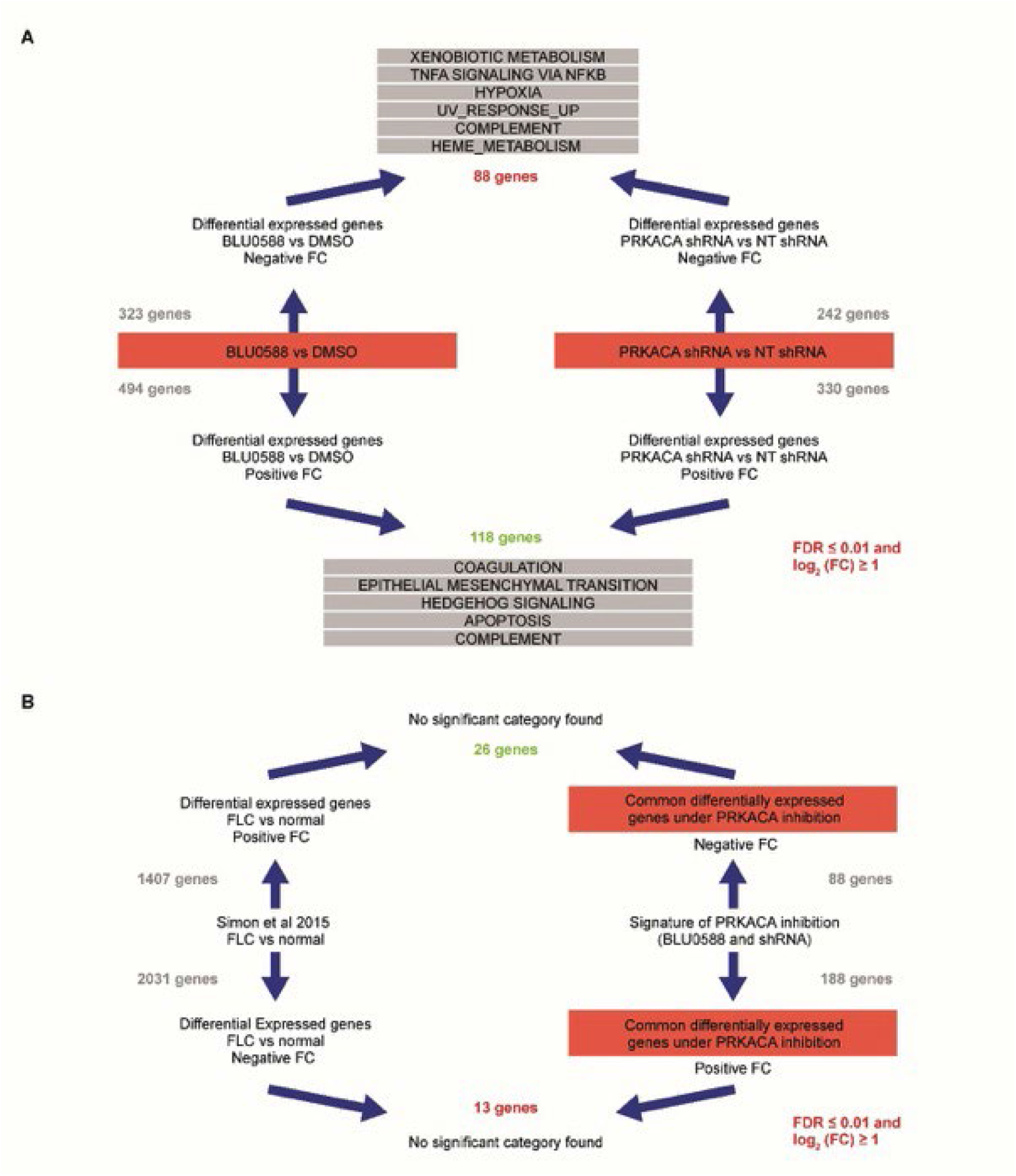
Identification of FLC-specific genes affected by PRKACA Inhibition. (A) Overlap of consensus shRNA with BLU0588 modulated gene expression. (B) Overlap of PRKACA inhibition signature with FLC gene signature.

To identify FLC pathways that are modulated in our signature of PRKACA inhibition and overlap with the Simon et al (14) gene set, we searched the canonical pathway collection available through the Molecular Signatures Database (32,33). We observed enrichment of pathways associated with liver and hepatocyte functions, including detoxification (drug metabolism of cytochrome P450 and biological oxidations), metabolism (retinol metabolism) and synthesis (steroid hormonal biosynthesis) with PRKACA inhibition, suggesting a reversal in FLC-affected pathways.

Since PRKACA is known to directly phosphorylate and modulate transcription factors (34), we searched for enrichment of transcription factor binding motifs from genes modulated by BLU0588, PRKACA shRNA knockdown, and the combined signature of PRKACA inhibition. Using the Hypergeometric Optimization of Motif Enrichment (33) motif finding tool, we identified previously described as well as novel transcription factor binding motifs (**Supplementary Figure S6a**). Hepatocyte nuclear factor 4 alpha (HNF4a) and JUN motifs are modulated in FLC (35), HNF4a is critical for liver-specific gene expression (36), and PKA directly phosphorylates and inhibits HNF4a function (37). When using the Clarivate Metacore analysis, we discovered that transcription factor estrogen related receptor alpha (ESRRA) appears to be the central direct target downstream of PRKACA, through which many other affected genes and transcription factors are connected. This interaction has previously been documented (38), and ESRRA is known to be central in hepatic metabolic dysfunction and diseases (39) (**Supplementary Figure S6b**).

To validate that PRKACA knockdown by shRNA or inhibition by BLU0588 treatment can reverse FLC-specific gene alterations, 4 genes were selected that are transcriptionally altered in FLC based on our RNA sequencing data: 2 upregulated genes, *CPS1* and glucose-6-phosphatase catalytic subunit (*G6PC)*; and 2 downregulated genes, glycogen phosphorylase, liver form (*PYGL)* and *SHH* (14). In 3 PRKACA shRNA-expressing cell lines, quantitative polymerase chain reaction assessment showed that upon doxycycline induction, *CPS1* and *G6PC* expression decreased, while *PYGL* and *SHH* increased in the 3 cell lines tested (**Figure 6a**). Expression of *G6PC* and *SHH* was unaffected by doxycycline in cells expressing non-targeting shRNA; however, expression of *CPS1* was moderately decreased and *PYGL* was moderately increased following doxycycline induction in cells expressing non-targeting shRNA. Similarly, in cells treated with BLU0588, expression of *CPS1* and *G6PC* were dose-dependently downregulated by BLU0588 treatment; this effect was observed within 24 hours of treatment, likely due to direct regulation. Expression of *PYGL* and *SHH* were dose-dependently upregulated by BLU0588 treatment, but this effect was observed after 14 days, suggesting this regulation is more indirect (**Figure 6b;** and **Supplementary Figs. S7a** and **S7b**). The IC50 values of these effects were between 55 nM and 180 nM, further suggesting gene expression can be modulated by directly inhibiting PRKACA kinase activity with BLU0588 (**Supplementary Figure S7**).

**Figure 6.**
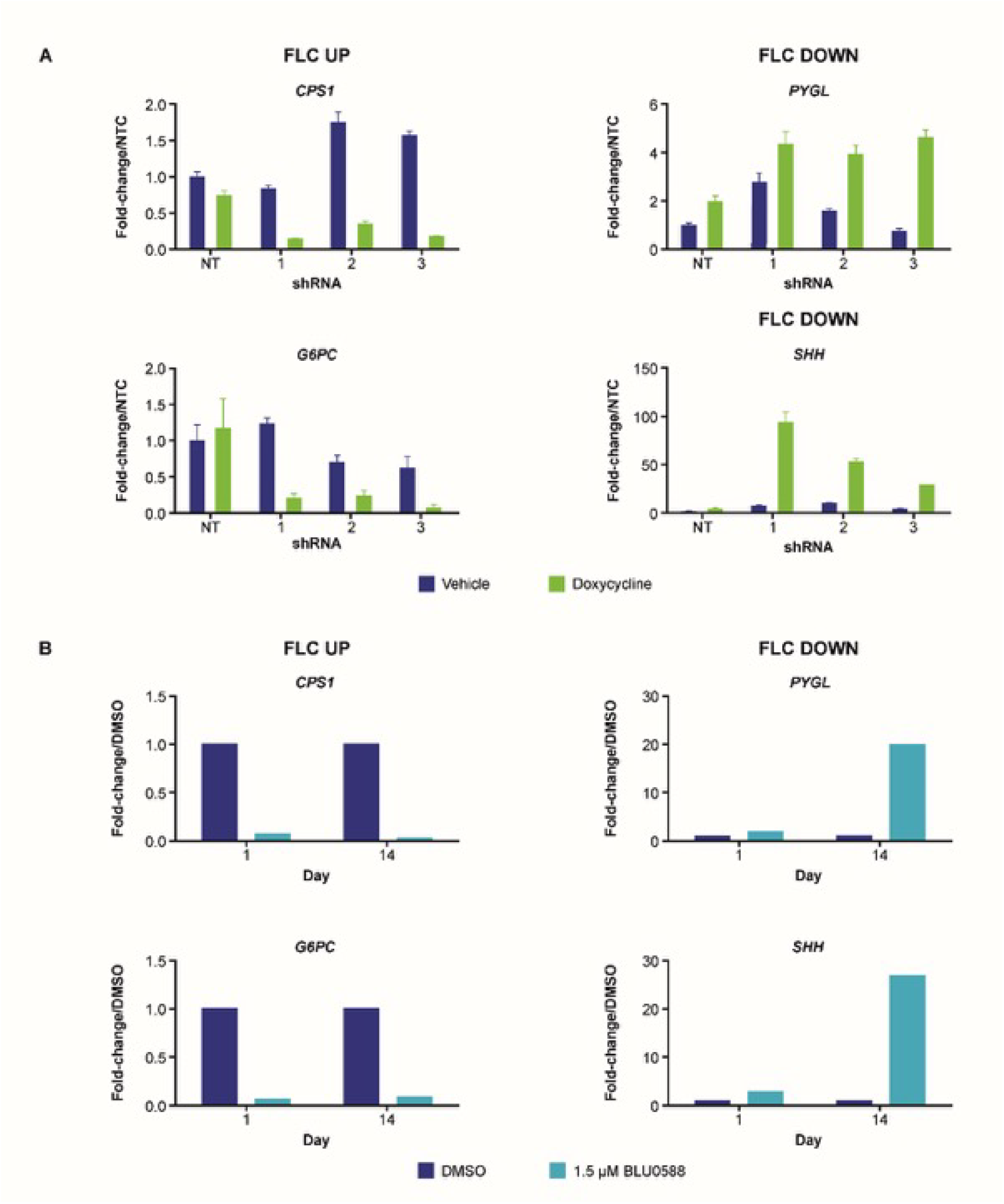
Effect of PRKACA knockdown or inhibition on FLC-specific gene expression. (A) Doxycycline reversed an FLC-specific gene signature, leading to downregulation of genes that are overexpressed in FLC (*CPS1* and *G6PC*, top) and upregulation of genes that are underexpressed in FLC (*PYGL* and *SHH*, bottom). Graph bars and error bars represent the mean ± SD (5 technical replicates and 3 independent biological replicates conducted on 3 different days). (B) BLU0588 treatment (1.5 μM for 1 day or 14 days) reversed a FLC-specific gene signature, leading to downregulation of genes that are overexpressed in FLC (*CPS1* and *G6PC*, top) and upregulation of genes that are underexpressed in FLC (*PYGL* and *SHH*, bottom; 5 technical replicates and 2 independent biological replicates).

## Discussion/Conclusion

This study is the first to evaluate the therapeutic potential for PRKACA inhibition in FLC. We characterized and validated an FLC PDX mouse model (27), and showed that PRKACA signaling and FLC-specific gene expression are dependent on DNAJB1-PRKACA fusion protein expression. Importantly, we demonstrated with 2 different modes of PRKACA inhibition that FLC PDX in vivo tumor growth was dependent on DNAJB1-PRKACA expression, supporting the notion that *DNAJB1-PRKACA* drives FLC tumor growth, and validating PRKACA as a therapeutic target for FLC. The selective PRKACA inhibitors developed in this study can also be used to validate PRKACA as a target for other diseases and to further study the molecular function of PRKACA.

We showed that DNAJB1-PRKACA fusion protein is overexpressed relative to WT PRKACA in an FLC PDX mouse model, consistent with reports of DNAJB1-PRKACA expression in FLC patient tumor samples (15,25). Moreover, downstream signaling of PRKACA was increased in FLC PDX tumors relative to Hep3B xenograft tumors that lack the DNAJB1-PRKACA fusion protein. Using this validated model of FLC PDX tumors, we showed that decreasing DNAJB1-PRKACA fusion protein levels both reduced downstream activity of the PRKACA pathway and reversed an FLC-specific gene signature. In PDX tumors, induction of PRKACA knockdown with doxycycline significantly reduced tumor growth, and the timing of this decrease in growth rate corresponded to the timing of DNAJB1-PRKACA knockdown and recovery after doxycycline treatment. No anti-proliferative effect was observed in vitro, potentially because of the slower in vitro proliferation rate, or because PRKACA might affect differentiation of more stem-like cells.

To our knowledge, BLU0588 and BLU2864 are the first selective small molecule PRKACA enzyme inhibitors. Using these molecules, we demonstrate the potential therapeutic value for FLC of inhibiting DNAJB1-PRKACA. In FLC PDX spheroid cultures, BLU0588 recapitulated the effects of DNAJB1-PRKACA knockdown. In vivo, treatment with BLU0588 and BLU2864 reduced tumor growth by approximately one-half relative to treatment with vehicle 34 days following the start of treatment. This finding was consistent with 3 different shRNA cell line xenografts. The pharmacokinetics of BLU0588 and BLU2864 show that, while 30 mg/kg QD did effectively inhibit downstream signaling of PRKACA, phosphorylated VASP levels returned to baseline levels by 24 hours. BLU0588 and BLU2864 are adenosine-5’-triphosphate (ATP)-competitive inhibitors that inhibit WT PRKACA in addition to the DNAJB1-PRKACA fusion protein. PRKACA is expressed in virtually all tissues and plays a critical role in cardiac function, making a strategy to selectively inhibit DNAJB1-PRKACA while sparing WT PRKACA desirable. The alternative conformation of the holoenzyme containing the DNAJB1-PRKACA fusion protein and the PKA regulatory subunit may constitute a strategy for selectively targeting the activity of the DNAJB1-PRKACA fusion protein (40). In addition, primary and metastatic FLC were shown to be sensitive to clinically available inhibitors of topoisomerase 1 and histone deacetylases, and to napabucasin (41), suggesting these agents as potential therapeutic strategies for FLC. The authors observed a variable response to PKA inhibitors, which may be due to the lack of sensitivity of the in vitro culture system (41).

Our findings have identified a gene signature dependent on PRKACA expression, and as such our gene set may contain potentially druggable targets for FLC. Some genes may be directly regulated by PRKACA through phosphorylation of transcription factors, such as HNFa (27), whereas other genes may be regulated in a more indirect manner. PRKACA regulation of these genes may potentially impact numerous cellular pathways which promote transformation and development of FLC. We observed only a partial overlap in gene expression changes between our findings and the analysis by Simon et al (14), likely due to using a single in vitro cultured PDX cell line, whereas the analysis by Simon et al was executed in a larger set of primary tumors and compared to adjacent normal tissue.

Although PRKACA is not a classical oncogene, constitutive activity of PRKACA and PKA signaling is known to play a pathogenic role in human disease and cancer. Activating mutations in PRKACA have been found to underly Cushing’s syndrome in patients with adrenal tumors (42-47), and constitutively active PKA signaling has also been shown to drive Carney Complex syndrome (48,49). In addition, PRKACA has been shown to mediate chemotherapy resistance (50). Recent data suggests *DNAJB1-PRKACA* fusions are not unique to FLC and may also occur in pancreatobiliary neoplasms (51).

In conclusion, this study supports the hypothesis that FLC tumor growth is dependent upon *DNAJB1-PRKACA* fusions and serves as an in vivo proof-of-concept that targeting hyperactive PRKACA signaling is a viable therapeutic strategy for patients with FLC. The development of selective PRKACA inhibitors also enables the validation of PRKACA as a therapeutic target for other diseases as well as for the study of the cellular function of PRKACA.

## Materials and Methods

### Generation of Inducible shRNA Cell Lines

FLC PDX tumors (Crown Biosciences) were dissociated with 2.5 mg/mL collagenase B (Sigma-Aldrich, #11088807001) in Roswell Park Memorial Institute 1640 medium (Thermo Fisher Scientific, #12633012). Mouse cells were depleted using a Mouse Cell Depletion Kit according to the manufacturer’s protocol (Miltenyi Biotech, #130-104-694). Dissociated FLC PDX cells were cultured as spheroids in Dulbecco’s Modified Eagle Medium (DMEM)/Nutrient Mixture F-12 (Thermo Fisher Scientific, #11330032) containing 20 ng/mL epidermal growth factor (StemCell Technologies, #02633), 10 ng/mL fibroblast growth factor (R&D Systems #233-FB-025), 2% B27 (Thermo Fisher Scientific, #12587010), and 1% N2 (Thermo Fisher Scientific, #17502048) on low-attachment plates. Cells were plated at 100,000 cells/well of 6-well plates (Corning) in culture medium and transduced with 106 viral particles containing PRKACA-targeting or NT shRNA (Horizon Discovery, SMARTvector™ Inducible Human PRKACA shRNA and SMARTvector™ Inducible Non-Targeting Control; PRKACA shRNA V3SH7670 – V3IHSHEG-8406259 [#1], V3ISHEG-6603667 [#2], V3ISHEG-5476684 [#3], V3IHSHEG_8059495 [#4], and NT snRNA VSC6586) in culture medium containing 8 μg/mL polybrene. Clones expressing shRNA constructs were selected for and maintained in 2 μg/mL puromycin.

### Western Blot Analysis

Frozen tumor sections of FLC PDX tumor tissue were homogenized in PhosphoSafe Extraction Reagent with added Halt™ protease inhibitor cocktail. Analysis of FLC PDX spheroids expressing inducible PRKACA shRNA, spheroids were treated with 0.5 μg/ml doxycycline or DMSO vehicle for 10 days, or with BLU0588, BLU2864, or DMSO vehicle for 4 hours. Cells were lysed in PhosphoSafe Extraction Reagent (EMD Millipore, #71296) with added Halt™ protease inhibitor cocktail (Thermo Fisher Scientific, #78430). Protein concentration was quantified using the Pierce BCA Protein Assay (Thermo Fisher Scientific, #23227). Proteins were resolved using SDS-PAGE on 20% gradient gels (Bio-Rad, #5671095) and electrotransferred onto nitrocellulose microporous membranes (Bio-Rad, #1704159). Immunodetection was performed using standard procedure and the following primary antibodies (all from Cell Signaling Technology): PRKACA #4782, pVASP #3114, and pCREB (#9198, ROCK #4563, and β-actin loading control #3700). Goat anti-rabbit secondary antibodies were purchased from Invitrogen (#A-21057) and Licor Biosciences (#926-32211), and chemiluminescent signals were detected using the LI-COR Biosciences Odyssey^®^ Imaging System.

### Quantitative Polymerase Chain Reaction

Lysates were prepared as described for western blotting. Total RNA was extracted using an RNeasy Plus Mini Kit (Qiagen, #74136) reagent/kit and reverse transcribed to complementary DNA (cDNA) using a High Capacity cDNA Reverse Transcription Kit (ThermoFisher Scientific, #4368813). The expression of *PYGL* (assay ID, Hs00958087_m1), *SHH* (assay ID, Hs01123832_m1), *CPS1* (assay ID, Hs00919483_m1), and *G6PC* (assay ID, Hs02560787_s1) was detected using TaqMan™ (FAM) Gene Expression Assays (ThermoFisher Scientific, #4331182) and the TaqMan Gene Expression Master Mix (ThermoFisher Scientific, #4369016), using the ViiA™ 7 Real-Time PCR System (ThermoFisher Scientific, #4453536). The relative quantification of gene expression was calculated using the 2-ΔΔCt method, and gene expression levels were normalized to a housekeeping control (*HMBS*; assay ID, Hs00609296_g1) using ViiA™ 7 software.

### Transcriptome Sequencing

To evaluate whether FLC gene expression signature depends on pathway activation by *DNAJB1-PRKACA* expression, we assessed gene expression by transcriptome sequencing (RNA sequencing) in the engineered shRNA cell lines. With DESeq2 (28) we assessed differential gene expression in PRKACA shRNA cell lines before and after doxycycline induction. Using a false discovery rate threshold of 0.01 and absolute log2 fold change of 1, we detected differentially expressed genes associated with each shPRKACA. A consensus of the differentially expressed genes was defined by overlapping (matching by the log2-fold change direction) significant DESeq2 results for the 3 PRKACA shRNA constructs used.

### Cell Proliferation Analyses

Dissociated FLC PDX cells were cultured as spheroids, transduced, and selected for expression of shRNA constructs as described above. Cells were plated in 96-well collagen-coated plates. PRKACA shRNA-expressing clones and cells expressing an NT shRNA were treated with 0.5 μg/ml doxycycline (for induction of shRNA) or DMSO vehicle and allowed to grow for 21 days. Proliferation was measured using a CellTiter-Glo® Luminescent Cell Viability Assay (Promega, #G7570) per the manufacturer’s instructions.

For determination of the effect of BLU0588 on cell proliferation in vitro, FLC PDX spheroids were cultured as above and were treated with BLU0588 at a range of concentrations, staurosporine, or DMSO vehicle. Proliferation was measured after 14 days as described above.

### Characterization of FLC PDX Tumors in Vivo

The PDX mouse model of FLC, LI5132 (27) (Crown Bioscience) was used, and a control CDX model generated in-house using a Hep3B HCC cell line as a control (Cat #HB-8064, American Type Culture Collection). An FLC PDX model was developed using inducible knockdown shRNA cell lines, which were generated as previously described above.

FLC PDX shRNA cell lines or Hep3B cells were implanted (500,000 cells) subcutaneously into the rear flank of 6-to 8-week-old female NOD-SCID mice (Jackson Laboratory, Bar Harbor, ME). Tumors were allowed to grow to an average size of 150–250 mm^3^. For shRNA knockdown experiments, mice were randomized into doxycycline treatment or control groups using the multi-task method in StudyLog software (South San Francisco, CA). ShRNA expression was induced using 2 mg/ml doxycycline and 5% sucrose added to drinking water; control mice received 5% sucrose in drinking water. Tumors were measured with calipers and mice were weighed twice weekly. Tumor volume was calculated using the formula volume = (length × width^2^)/2.

For immunohistochemical staining in formalin-fixed, paraffin-embedded FLC PDX tumors, staining was conducted on a Leica Bond RXm (Leica Biosystems) using standard chromogenic methods. For heat-induced epitope retrieval, slides were heated in either a pH6 citrate-based buffer for 20 minutes at 96°C (glypican 3, cytokeratin 7, Ki67), or a pH9 ethylenediaminetetraacetic acid (EDTA)-based buffer for 20 minutes at 96°C (CD68), followed by a 15-minute antibody incubation. Antibody binding for Ki67 (Abcam, #ab16667), CD68 (Abcam, #ab955), cytokeratin 7 (Abcam, #ab181598), and glypican 3 (Cell Marque, #261M) was detected using BOND Polymer Refine Detection (Leica Biosystems, #DS9800), including a hematoxylin counterstain to visualize nuclei.

### Characterization of BLU0588 and BLU2864

BLU0588 and BLU2864 were developed from screening hits identified from a proprietary library of compounds. PRKACA inhibiting activity was screened using the EZ Reader 2 electrophoretic mobility shift platform (PerkinElmer). PRKACA enzyme (0.007 ng/mL; Millipore, #539482) or ROCK2 enzyme (0.006 ng/mL; SignalChem, #R11-11H-10) was added to each well of a 384-well plate containing 1 μM Kemptide peptide substrate (5-FAM-LRRASLG; AnaSpec 2933), Km concentrations of ATP (5 μM ATP), and a concentration series of test compounds (1% final DMSO) in (100 mM HEPES buffer pH 7.5, 0.015% Brij-35, 10 mM MgCl2, 1 mM DTT) and incubated for 90 minutes at 25°C. The reaction was stopped with addition of Stop buffer (100 mM HEPES pH 7.5, 0.015% Brij-35, 35 mM EDTA, and 0.2% Coating Reagent 3 [PerkinElmer]). Results were read with the EZ Reader 2 and the IC50 was calculated using a 4-parameter fit. For the AKT1-3 Nanosyn enzyme assay, test compounds were diluted in 100% DMSO using 3-fold dilution steps. The final compound concentration in the assay ranged from 10 μM to 0.056 nM, and compounds were tested in a single well for each dilution, with the final concentration of DMSO in all assays kept at 1%. The reference compound, staurosporine, was tested in an identical manner.

Phosphorylation of VASP on Serine 157 was used as a readout of PRKACA cellular activity. Phosphorylation was detected with a homogeneous time-resolved fluorescence assay, following the manufacturer’s protocol (Cisbio, #63ADK066PEH). PKA was activated by the addition of forskolin (Sigma-Aldrich, #F3917), and the dose response to PRKACA inhibitors was measured in forskolin-stimulated human Huh7 cells as follows. Briefly, Huh7 cells were plated at a density of 2 x 104 cells per well in a 384-well opti-well cell culture plate in 15 μl of serum- and phenol-free DMEM (Gibco, #21063-029), and incubated overnight at 37°C, 5% CO2. The next day, 3 μl of a dosed concentration series of test compound (0.24% DMSO final concentration) was added to the wells, and the cells were incubated for an additional 4 hours at 37°C, 5% CO2. Two μl of forskolin was added at a final concentration of 5 μM and the plates were incubated for 30 minutes at 37°C, 5% CO2. Five μl of lysis buffer containing 1% Halt™ protease cocktail inhibitors (ThermoFisher, #78430), was added to the cells and the mixture was incubated under gentle shaking for 30 min at room temperature. Ten μl of this lysate was transferred to a 384-well proxi plate and 2.5 μL of the premixed antibody solution was added. An antibody solution was prepared by combining phospho-VASP cryptate antibody and phospho-VASP d2 antibody, at 20-fold dilution into a buffer solution following the manufacturer’s protocol. The lysate and antibody mixture was incubated for either 3 h at room temperature or overnight at 4°C. The fluorescence emission at 2 different wavelengths (665 nm and 620 nm) was read on an EnVision instrument.

For in vivo studies of BLU0588 or BLU2864, FLC PDX tumor cells were implanted subcutaneously into the rear flank of 6-to 8-week-old female NOD-SCID mice (Jackson Laboratory, Bar Harbor, ME) and allowed to grow to 150 mm3 for efficacy studies or 500 mm3 for pharmacokinetic/pharmacodynamic studies. Mice were randomized into control or treatment groups using the multi-task method in Study Log software. BLU0588 was dissolved in 20% Solutol in 0.5% methylcellulose and BLU2864 was dissolved in 10% DMSO, 10% Solutol HS15, 20% hydroxypropyl-cyclodextrin, and administered orally, once daily. Tumor size and mouse body weight was determined as above. For pharmacokinetic/pharmacodynamic studies, plasma and tumors were collected at pre-specified time points, and BLU0588 and BLU2864 were detected using Liquid Chromatography Triple Quadrupole Mass Spectrometry. Phospho-VASP was measured using western blotting as described above.

## Abbreviations

ATP: adenosine-5’-triphosphate
cAMP: 3′ 5′-cyclic adenosine monophosphate
CCC: cholangiocellular carcinoma
cDNA: complementary DNA
CDX: cell line derived xenograft
CPS1: carbamoyl phosphate synthetase 1
CREB: cAMP-responsive binding protein
DMEM: Dulbecco’s Modified Eagle Medium
DMSO: dimethyl sulfoxide
DNAJB1: heat shock protein 40
EDTA: ethylenediaminetetraacetic acid
ESRRA: estrogen related receptor alpha
FLC: fibrolamellar carcinoma
G6PC: glucose-6-phosphatase catalytic subunit
HNF4a: hepatocyte nuclear factor 4 alpha
HCC: hepatocellular carcinoma
IC_50_: half-maximal inhibitory concentration
K_d_: dissociation constant
NT: non-targeting
PDX: patient-derived xenograft
PKA: protein kinase A
PRKACA: PKA alpha catalytic subunit
PYGL: glycogen phosphorylase liver form
QD: once daily
SHH: sonic hedgehog
shRNA: short hairpin ribonucleic acid
VASP: vasodilator-stimulated phosphoprotein
WT: wild type

## Acknowledgements

Medical writing support was provided by Allison Cherry, PhD, and Elizabeth G. Wheatley, PhD, of Nexus Global Group Science, LLC, and editorial support, including formatting, proofreading, and submission was provided by Travis Taylor, BA, Paragon, Knutsford, supported by Blueprint Medicines Corporation according to Good Publication Practice guidelines (www.ismpp.org/gpp3). Lola Reid, PhD, of University of North Carolina at Chapel Hill, was involved in the development of the initial manuscript draft. The Sponsor was involved in the study design, collection, analysis and interpretation of data, as well as data checking of information provided in the manuscript. However, ultimate responsibility for opinions, conclusions, and data interpretation lies with the authors.

## References

1. Riggle KM, Turnham R, Scott JD, Yeung RS, Riehle KJ. Fibrolamellar Hepatocellular Carcinoma: Mechanistic Distinction From Adult Hepatocellular Carcinoma. Pediatr Blood Cancer 2016;63:1163-7

2. Eggert T, McGlynn KA, Duffy A, Manns MP, Greten TF, Altekruse SF. Fibrolamellar hepatocellular carcinoma in the USA, 2000-2010: A detailed report on frequency, treatment and outcome based on the Surveillance, Epidemiology, and End Results database. United European Gastroenterol J 2013;1:351-7

3. Bayer HealthCare Pharmaceuticals Inc. 2018 NEXAVAR (sorafenib). Prescribing Information.

4. Ang CS, Kelley RK, Choti MA, Cosgrove DP, Chou JF, Klimstra D, et al. Clinicopathologic characteristics and survival outcomes of patients with fibrolamellar carcinoma: data from the fibrolamellar carcinoma consortium. Gastrointest Cancer Res 2013;6:3-9

5. Kaseb AO, Shama M, Sahin IH, Nooka A, Hassabo HM, Vauthey JN, et al. Prognostic indicators and treatment outcome in 94 cases of fibrolamellar hepatocellular carcinoma. Oncology 2013;85:197-203

6. Groeschl RT, Miura JT, Wong RK, Bloomston M, Lidsky ML, Clary BM, et al. Multi-institutional analysis of recurrence and survival after hepatectomy for fibrolamellar carcinoma. J Surg Oncol 2014;110:412-5

7. Stipa F, Yoon SS, Liau KH, Fong Y, Jarnagin WR, D’Angelica M, et al. Outcome of patients with fibrolamellar hepatocellular carcinoma. Cancer 2006;106:1331-8

8. Chakrabarti S, Tella SH, Kommalapati A, Huffman BM, Yadav S, Riaz IB, et al. Clinicopathological features and outcomes of fibrolamellar hepatocellular carcinoma. J Gastrointest Oncol 2019;10:554-61

9. Ward SC, Waxman S. Fibrolamellar carcinoma: a review with focus on genetics and comparison to other malignant primary liver tumors. Semin Liver Dis 2011;31:61-70

10. Ward SC, Huang J, Tickoo SK, Thung SN, Ladanyi M, Klimstra DS. Fibrolamellar carcinoma of the liver exhibits immunohistochemical evidence of both hepatocyte and bile duct differentiation. Mod Pathol 2010;23:1180-90

11. Ross HM, Daniel HD, Vivekanandan P, Kannangai R, Yeh MM, Wu TT, et al. Fibrolamellar carcinomas are positive for CD68. Mod Pathol 2011;24:390-5

12. Capurro M, Wanless IR, Sherman M, Deboer G, Shi W, Miyoshi E, et al. Glypican-3: a novel serum and histochemical marker for hepatocellular carcinoma. Gastroenterology 2003;125:89-97

13. Dinh TA, Vitucci EC, Wauthier E, Graham RP, Pitman WA, Oikawa T, et al. Comprehensive analysis of The Cancer Genome Atlas reveals a unique gene and non-coding RNA signature of fibrolamellar carcinoma. Sci Rep 2017;7:44653

14. Simon EP, Freije CA, Farber BA, Lalazar G, Darcy DG, Honeyman JN, et al. Transcriptomic characterization of fibrolamellar hepatocellular carcinoma. Proc Natl Acad Sci U S A 2015;112:E5916-25

15. Honeyman JN, Simon EP, Robine N, Chiaroni-Clarke R, Darcy DG, Lim, II, et al. Detection of a recurrent DNAJB1-PRKACA chimeric transcript in fibrolamellar hepatocellular carcinoma. Science 2014;343:1010-4

16. Cornella H, Alsinet C, Sayols S, Zhang Z, Hao K, Cabellos L, et al. Unique genomic profile of fibrolamellar hepatocellular carcinoma. Gastroenterology 2015;148:806-18 e10

17. Engelholm LH, Riaz A, Serra D, Dagnaes-Hansen F, Johansen JV, Santoni-Rugiu E, et al. CRISPR/Cas9 Engineering of Adult Mouse Liver Demonstrates That the Dnajb1-Prkaca Gene Fusion Is Sufficient to Induce Tumors Resembling Fibrolamellar Hepatocellular Carcinoma. Gastroenterology 2017;153:1662-73 e10

18. Kastenhuber ER, Lalazar G, Houlihan SL, Tschaharganeh DF, Baslan T, Chen CC, et al. DNAJB1-PRKACA fusion kinase interacts with beta-catenin and the liver regenerative response to drive fibrolamellar hepatocellular carcinoma. Proc Natl Acad Sci U S A 2017;114:13076-84

19. Turnham RE, Smith FD, Kenerson HL, Omar MH, Golkowski M, Garcia I, et al. An acquired scaffolding function of the DNAJ-PKAc fusion contributes to oncogenic signaling in fibrolamellar carcinoma. Elife 2019;8

20. Dinh TA, Jewell ML, Kanke M, Francisco A, Sritharan R, Turnham RE, et al. MicroRNA-375 Suppresses the Growth and Invasion of Fibrolamellar Carcinoma. Cell Mol Gastroenterol Hepatol 2019;7:803-17

21. Graham RP, Jin L, Knutson DL, Kloft-Nelson SM, Greipp PT, Waldburger N, et al. DNAJB1-PRKACA is specific for fibrolamellar carcinoma. Mod Pathol 2015;28:822-9

22. Turnham RE, Scott JD. Protein kinase A catalytic subunit isoform PRKACA; History, function and physiology. Gene 2016;577:101-8

23. Wahlang B, McClain C, Barve S, Gobejishvili L. Role of cAMP and phosphodiesterase signaling in liver health and disease. Cell Signal 2018;49:105-15

24. Olivieri C, Walker C, Karamafrooz A, Wang Y, Manu VS, Porcelli F, et al. Defective internal allosteric network imparts dysfunctional ATP/substrate-binding cooperativity in oncogenic chimera of protein kinase A. Commun Biol 2021;4:321

25. Riggle KM, Riehle KJ, Kenerson HL, Turnham R, Homma MK, Kazami M, et al. Enhanced cAMP-stimulated protein kinase A activity in human fibrolamellar hepatocellular carcinoma. Pediatr Res 2016;80:110-8

26. Cheung J, Ginter C, Cassidy M, Franklin MC, Rudolph MJ, Robine N, et al. Structural insights into mis-regulation of protein kinase A in human tumors. Proc Natl Acad Sci U S A 2015;112:1374-9

27. Thatte J, Talaoc EC, Scott C, Ren K, Broudy T. Kinase Inhibitor Demonstrates Efficacy in a Patient-Derived Xenograft Model of Fibrolamellar Hepatocellular Carcinoma Featuring DNAJB1-PRKACA Fusion. 2017. p 1074.

28. Love MI, Huber W, Anders S. Moderated estimation of fold change and dispersion for RNA-seq data with DESeq2. Genome Biol 2014;15:550

29. Leinonen MK, Miettinen J, Heikkinen S, Pitkaniemi J, Malila N. Quality measures of the population-based Finnish Cancer Registry indicate sound data quality for solid malignant tumours. Eur J Cancer 2017;77:31-9

30. Davis MI, Hunt JP, Herrgard S, Ciceri P, Wodicka LM, Pallares G, et al. Comprehensive analysis of kinase inhibitor selectivity. Nat Biotechnol 2011;29:1046-51

31. Patro R, Duggal G, Love MI, Irizarry RA, Kingsford C. Salmon provides fast and bias-aware quantification of transcript expression. Nat Methods 2017;14:417-9

32. Liberzon A, Birger C, Thorvaldsdottir H, Ghandi M, Mesirov JP, Tamayo P. The Molecular Signatures Database (MSigDB) hallmark gene set collection. Cell Syst 2015;1:417-25

33. Subramanian A, Tamayo P, Mootha VK, Mukherjee S, Ebert BL, Gillette MA, et al. Gene set enrichment analysis: a knowledge-based approach for interpreting genome-wide expression profiles. Proc Natl Acad Sci U S A 2005;102:15545-50

34. Isobe K, Jung HJ, Yang CR, Claxton J, Sandoval P, Burg MB, et al. Systems-level identification of PKA-dependent signaling in epithelial cells. Proc Natl Acad Sci U S A 2017;114:E8875-E84

35. Dinh TA, Sritharan R, Smith FD, Francisco AB, Ma RK, Bunaciu RP, et al. Hotspots of Aberrant Enhancer Activity in Fibrolamellar Carcinoma Reveal Candidate Oncogenic Pathways and Therapeutic Vulnerabilities. Cell Rep 2020;31:107509

36. Spath GF, Weiss MC. Hepatocyte nuclear factor 4 expression overcomes repression of the hepatic phenotype in dedifferentiated hepatoma cells. Mol Cell Biol 1997;17:1913-22

37. Viollet B, Kahn A, Raymondjean M. Protein kinase A-dependent phosphorylation modulates DNA-binding activity of hepatocyte nuclear factor 4. Mol Cell Biol 1997;17:4208-19

38. Liu D, Benlhabib H, Mendelson CR. cAMP enhances estrogen-related receptor alpha (ERRalpha) transcriptional activity at the SP-A promoter by increasing its interaction with protein kinase A and steroid receptor coactivator 2 (SRC-2). Mol Endocrinol 2009;23:772-83

39. Xia H, Dufour CR, Giguere V. ERRalpha as a Bridge Between Transcription and Function: Role in Liver Metabolism and Disease. Front Endocrinol (Lausanne) 2019;10:206

40. Cao B, Lu TW, Martinez Fiesco JA, Tomasini M, Fan L, Simon SM, et al. Structures of the PKA RIalpha Holoenzyme with the FLHCC Driver J-PKAcalpha or Wild-Type PKAcalpha. Structure 2019;27:816-28 e4

41. Lalazar G, Requena D, Ramos-Espiritu L, Ng D, Bhola PD, de Jong YP, et al. Identification of Novel Therapeutic Targets for Fibrolamellar Carcinoma Using Patient Derived Xenografts and Direct from Patient Screening. Cancer Discov 2021

42. Beuschlein F, Fassnacht M, Assie G, Calebiro D, Stratakis CA, Osswald A, et al. Constitutive activation of PKA catalytic subunit in adrenal Cushing’s syndrome. N Engl J Med 2014;370:1019-28

43. Cao Y, He M, Gao Z, Peng Y, Li Y, Li L, et al. Activating hotspot L205R mutation in PRKACA and adrenal Cushing’s syndrome. Science 2014;344:913-7

44. Di Dalmazi G, Kisker C, Calebiro D, Mannelli M, Canu L, Arnaldi G, et al. Novel somatic mutations in the catalytic subunit of the protein kinase A as a cause of adrenal Cushing’s syndrome: a European multicentric study. J Clin Endocrinol Metab 2014;99:E2093-100

45. Goh G, Scholl UI, Healy JM, Choi M, Prasad ML, Nelson-Williams C, et al. Recurrent activating mutation in PRKACA in cortisol-producing adrenal tumors. Nat Genet 2014;46:613-7

46. Nakajima Y, Okamura T, Gohko T, Satoh T, Hashimoto K, Shibusawa N, et al. Somatic mutations of the catalytic subunit of cyclic AMP-dependent protein kinase (PRKACA) gene in Japanese patients with several adrenal adenomas secreting cortisol [Rapid Communication]. Endocr J 2014;61:825-32

47. Sato Y, Maekawa S, Ishii R, Sanada M, Morikawa T, Shiraishi Y, et al. Recurrent somatic mutations underlie corticotropin-independent Cushing’s syndrome. Science 2014;344:917-20

48. Forlino A, Vetro A, Garavelli L, Ciccone R, London E, Stratakis CA, et al. PRKACB and Carney complex. N Engl J Med 2014;370:1065-7

49. Yin Z, Pringle DR, Jones GN, Kelly KM, Kirschner LS. Differential role of PKA catalytic subunits in mediating phenotypes caused by knockout of the Carney complex gene Prkar1a. Mol Endocrinol 2011;25:1786-93

50. Moody SE, Schinzel AC, Singh S, Izzo F, Strickland MR, Luo L, et al. PRKACA mediates resistance to HER2-targeted therapy in breast cancer cells and restores anti-apoptotic signaling. Oncogene 2015;34:2061-71

51. Vyas M, Hechtman JF, Zhang Y, Benayed R, Yavas A, Askan G, et al. DNAJB1-PRKACA fusions occur in oncocytic pancreatic and biliary neoplasms and are not specific for fibrolamellar hepatocellular carcinoma. Mod Pathol 2020;33:648-56

