## Supplementary Material for "Evaluation of PRKACA as a Therapeutic Target for Fibrolamellar Carcinoma"

### Supplementary information

**Supplementary Table S1.** Characterization of BLU0588 and BLU2864

| | **BLU0588** | **BLU2864** |
|---|---|---|
| **PRKACA Biochemical** IC50 (@ ATP KM) | 1 nM | 0.3 nM |
| **PRKACB Biochemical** IC50 (@ ATP KM) | 0.8 nM | 0.7 nM |
| **ROCK2 Biochemical** IC50 (@ATP KM) | 83.1 nM | 12.7 nM |
| **AKT1 Biochemical IC50 (@ATP KM)** | 1540 nM | 2120.0 nM |
| **AKT2 Biochemical IC50 (@ATP KM)** | 3780 nM | 4910.0 nM |
| **AKT3 Biochemical IC50 (@ATP KM)** | 397 nM | 475.0 nM |
| **PRKX Biochemical IC50 (@ATP KM)** | 25.2 nM | 6.8 nM |
| Cell Huh7 **p-VASP** IC50 | 25 nM | 36.6 nM |
| FLC PDX **p-VASP** IC50 | 31 nM | |
| Cell ROCK1/2 **p-MYPT1** WB IC50 | 303 nM | 240 nM |
| S(10@3uM) | 0.047 | 0.057 |

**Supplementary Table S2.** Binding profile of BLU0588 and BLU2864

| | BLU0588 | |
|---|---|---|
| **Target** | **PoC** | **Kd (nM)** |
| PKAC-alpha | 0.4 | 4 |
| PKAC-beta | 2.4 | 4.8 |
| ROCK2 | 0 | 5 |
| ROCK1 | 0.1 | 5 |
| HASPIN | 0.3 | 21 |
| JAK3(JH1domain-catalytic) | 0 | 51.5 |
| MYLK4 | 0.3 | 55 |
| JAK2(JH1domain-catalytic) | 2.1 | 58 |
| PRKX | 4.6 | 61 |
| SNARK | 2.1 | 78 |
| SgK110 | 7.1 | 105 |
| PRKG2 | 9.5 | 105 |
| PRKG1 | 3.1 | 160 |
| PAK4 | 6.5 | 170 |
| PAK7 | 1.6 | 180 |
| YSK4 | 1.2 | 185 |
| PRKCE | 0 | 200 |
| PRKCD | 5.6 | 250 |
| GRK7 | 21 | 310 |
| RET | 9 | 315 |
| LATS2 | 0 | 445 |
| MEK3 | 12 | 470 |
| PKN1 | 8.6 | 520 |
| RPS6KA4(Kin.Dom.2-C-terminal) | 8.7 | 630 |
| TYK2(JH1domain-catalytic) | 12 | 690 |
| GRK1 | 25 | 710 |
| JAK1(JH1domain-catalytic) | 0 | 790 |
| RSK4(Kin.Dom.1-N-terminal) | 19 | 1005 |
| ARK5 | 0.8 | 1500 |
| RSK2(Kin.Dom.1-N-terminal) | 26 | 1650 |
| MEK1 | 49 | 1900 |
| GCN2(Kin.Dom.2,S808G) | 0 | 2050 |
| MEK4 | 56 | 2700 |
| STK33 | 37 | 3000 |
| MEK2 | 55 | 3000 |
| CDC2L5 | 3.1 | 10000 |

| | BLU2864 | |
|---|---|---|
| **Target** | **PoC** | **Kd (nM)** |
| PKAC-alpha | 4.5 | 3.3 |
| ROCK2 | 0.2 | 4.8 |
| ROCK1 | 0.4 | 5.4 |
| PKAC-beta | 0.7 | 6.3 |
| PRKX | 0 | 16.5 |
| SNARK | 1.1 | 26 |
| JAK2(JH1domain-catalytic) | 0.7 | 27.5 |
| MEK1 | 0.3 | 29 |
| JAK3(JH1domain-catalytic) | 7 | 50 |
| MEK4 | 2.6 | 68.5 |
| TYK2(JH1domain-catalytic) | 5.8 | 99.5 |
| MEK3 | 4.5 | 110 |
| LATS2 | 0 | 140 |
| PRKG2 | 18 | 165 |
| YSK4 | 4 | 180 |
| PRKG1 | 8.5 | 195 |
| RSK4(Kin.Dom.1-N-terminal) | 7.4 | 245 |
| MEK2 | 1.1 | 255 |
| MYLK4 | 3.9 | 320 |
| BIKE | 12 | 370 |
| HASPIN | 11 | 382.5 |
| GRK7 | 3.9 | 385 |
| IKK-beta | 14 | 400 |
| STK33 | 8 | 420 |
| PAK4 | 17 | 420 |
| RSK2(Kin.Dom.1-N-terminal) | 5.1 | 470 |
| PRKCD | 14 | 580 |
| GRK1 | 9.3 | 650 |
| PAK7 | 14 | 675 |
| JAK1(JH1domain-catalytic) | 32 | 675 |
| RET | 29 | 850 |
| ARK5 | 42 | 920 |
| PRKCE | 18 | 1010 |
| PKN1 | 59 | 1600 |
| SgK110 | 23 | 1650 |
| JAK1(JH2domain-pseudokinase) | 3.7 | 2300 |
| RPS6KA4(Kin.Dom.2-C-terminal) | 61 | 3000 |

**Supplementary Table S3.** Percent PRKACA fusion reads.

| Sample | Percentage |
|---|---|
| DMSO-1 | 94 |
| DMSO-2 | 95 |
| BLU0588-1 | 95 |
| BLU0588-2 | 93 |
| shPRKACA #1 no Dox-1 | 89 |
| shPRKACA #1 no Dox-2 | 82 |
| shPRKACA #1 + Dox-1 | 85 |
| shPRKACA #1 + Dox-2 | 84 |
| shPRKACA #2 no Dox-1 | 91 |
| shPRKACA #2 no Dox-2 | 92 |
| shPRKACA #2 + Dox-1 | 91 |
| shPRKACA #2 + Dox-2 | 97 |
| shPRKACA #3 no Dox-1 | 92 |
| shPRKACA #3 no Dox-2 | 89 |
| shPRKACA #3 + Dox-1 | 81 |
| shPRKACA #3 + Dox-2 | 89 |

PRAGUE fusion vs WT: [Fusion reads/ (Fusion + WT reads)] * 100

**Supplementary Table 4A.** Consensus PRKACA shRNA modulated genes

| log2FC_Pos_Simon | Overlap-All3_shRNA-NEG | Overlap |
|---|---|---|
| MARCH4 | PPARGC1A | AKR1B15 |
| MARCH10 | FKBP11 | AKR1C3 |
| AACS | PDE4B | ATP1B1 |
| AAGAB | ENTPD1 | C8G |
| AAK1 | CPS1 | CA9 |
| ABCA12 | DDIT4 | CABYR |
| ABCA2 | SLC16A11 | CALCA |
| ABCA3 | RHOBTB1 | CGREF1 |
| ABCA7 | C8G | COL5A3 |
| ABCB5 | COL5A3 | CORIN |
| ABCB8 | NEIL3 | CPLX2 |
| ABCC5 | AKR1C3 | CPS1 |
| ABCF2 | CGREF1 | CTH |
| ABHD17C | TMOD1 | DDIT4 |
| ABHD4 | GLA | E2F8 |
| ABLIM2 | HPD | ENTPD1 |
| AC006132.1 | UBXN10 | EPS8L3 |
| AC008443.1 | SLC5A6 | EVA1C |
| AC010547.9 | TLL2 | FKBP11 |
| AC096677.1 | KYNU | FSTL4 |
| AC145676.2 | ATP1B1 | GK |
| ACAN | E2F8 | GLA |
| ACKR3 | TM4SF5 | GPRIN3 |
| ACLY | CTH | HKDC1 |
| ACSL4 | PRKACA | HPD |
| ACYP1 | KCNE4 | KCNE4 |
| ADAM12 | GK | KCNU1 |
| ADAM22 | EVA1C | KRT86 |
| ADAM32 | CA9 | KYNU |
| ADAMDEC1 | SLC7A1 | MUC13 |
| ADAMTS14 | CORIN | MUC5B |
| ADAMTS15 | GPRIN3 | NEB |
| ADAMTS18 | SLC22A11 | NEIL3 |
| ADAMTS6 | PBK | PBK |
| ADAMTS9 | PDE3A | PDE3A |
| ADAMTSL5 | UTS2B | PDE4B |
| ADAT1 | SDCBP2 | PPARGC1A |
| ADAT2 | HKDC1 | PRKACA |
| ADCY2 | CABYR | RHOBTB1 |
| ADD2 | NEB | S100P |

| | | |
|---|---|---|
| ADRA2A | KRT86 | SDCBP2 |
| ADSSL1 | EPS8L3 | SLC16A11 |
| AEN | TESC | SLC22A11 |
| AF131215.5 | FSTL4 | SLC5A6 |
| AFAP1 | AKR1B15 | SLC7A1 |
| AGA | S100P | TESC |
| AGR2 | MUC5B | TLL2 |
| AGRN | KCNU1 | TM4SF5 |
| AHI1 | CALCA | TMOD1 |
| AHNAK2 | CPLX2 | UBXN10 |
| AHRR | MUC13 | UTS2B |
| AIFM2 | CPB2 | |
| AIM1L | CLDN2 | |
| AK8 | PTGS2 | |
| AKAP12 | SMIM9 | |
| AKR1B15 | SCARA5 | |
| AKR1C3 | CYP24A1 | |
| AKTIP | UGT2B10 | |
| AL133373.1 | ROPN1L | |
| AL358813.2 | APOB | |
| AL359878.1 | C11orf86 | |
| ALDH18A1 | PDK4 | |
| ALDH1A2 | SLC39A5 | |
| ALDH1L2 | ADH4 | |
| ALDOA | MPP1 | |
| ALG1 | HEPACAM | |
| ALPK3 | CA2 | |
| AMFR | PCK1 | |
| ANKRD18A | C8A | |
| ANKRD22 | ADH6 | |
| ANKRD29 | ASGR1 | |
| ANKRD52 | ABCG2 | |
| ANKS6 | C3 | |
| ANLN | ALDOB | |
| ANO2 | KHK | |
| ANO4 | PRODH2 | |
| ANTXR1 | LYPD3 | |
| ANXA2 | S100A14 | |
| ANXA5 | PLA2G4A | |
| AP1S3 | PKLR | |
| APCDD1L | ALDH1L1 | |
| APLN | PDE7B | |
| APLP1 | SIK1 | |

| | |
|---|---|
| APOO | CU639417.2 |
| AQPEP | MTTP |
| AREG | TNFSF11 |
| ARG2 | G6PC |
| ARHGAP11A | SLC6A4 |
| ARHGAP11B | SULT1B1 |
| ARHGAP18 | ANPEP |
| ARHGAP22 | DEGS2 |
| ARHGAP36 | SLC23A1 |
| ARHGAP39 | PAGE4 |
| ARHGEF35 | UGT2B17 |
| ARHGEF39 | ANKS4B |
| ARHGEF5 | GPT |
| ARL14 | PGC |
| ARL2BP | MAT1A |
| ARMC9 | HPN |
| ARNTL2 | PKDCC |
| AS3MT | ARHGEF16 |
| ASB15 | TCF7L1 |
| ASIC1 | CDHR5 |
| ASNS | RBFOX3 |
| ASPHD1 | HMOX1 |
| ASRGL1 | UGT2A3 |
| ATIC | F2 |
| ATP1B1 | AKR1B10 |
| ATP2A1 | PREB |
| ATP2B4 | SLC16A13 |
| ATP6V1B1 | UGT1A4 |
| ATP8A2 | SSTR1 |
| ATP8B3 | FMO5 |
| ATR | XPNPEP2 |
| AURKA | PAH |
| B3GNT3 | NOSTRIN |
| B3GNT5 | SLCO2B1 |
| B3GNTL1 | GOLGA6A |
| B4GALNT1 | AL132639.3 |
| B9D1 | DDC |
| BAG2 | PCCA |
| BAG3 | HRCT1 |
| BAI2 | MYO1A |
| BAIAP2L2 | AGMAT |
| BBC3 | ANG |
| BBS2 | APOC4-APOC2 |

| | |
|---|---|
| BCAR1 | CYP4F3 |
| BCAT1 | HYAL1 |
| BCAT2 | AL163636.2 |
| BCL11A | SRGN |
| BCL2L2-PABPN1 | SLC13A3 |
| BDKRB1 | CD55 |
| BDKRB2 | GAS2 |
| BDNF | RND1 |
| BEND3 | FGL1 |
| BEND6 | CDHR2 |
| BEST3 | CBR1 |
| BFSP1 | HNF4A |
| BICD1 | CLDN7 |
| BIRC5 | FERMT1 |
| BLM | DNAJC12 |
| BLVRA | PEX11A |
| BMP6 | CD68 |
| BMP8A | GSTA2 |
| BMP8B | SLC19A3 |
| BNC2 | CMBL |
| BRE | F7 |
| BRSK2 | GGH |
| BSG | DACT2 |
| BUB1 | IQCH |
| BUB1B | NR4A1 |
| C10orf128 | CREB3L3 |
| C10orf131 | THRB |
| C10orf2 | ENTPD2 |
| C10orf90 | SLC43A1 |
| C12orf39 | CYP4F11 |
| C12orf5 | SERPINA10 |
| C14orf183 | PC |
| C15orf65 | RNASE4 |
| C16orf46 | TTPA |
| C16orf59 | WIF1 |
| C16orf93 | TUBAL3 |
| C18orf56 | SOWAHA |
| C19orf54 | LENG9 |
| C1orf105 | PTGR1 |
| C1orf198 | PDE8B |
| C1orf229 | NGEF |
| C1QL1 | ESRP2 |
| C20orf96 | CCL15 |

| | |
|---|---|
| C2CD4A | PER2 |
| C2orf27A | GCLM |
| C2orf66 | PIPOX |
| C2orf81 | DUSP1 |
| C3orf36 | MLKL |
| C3orf52 | EDNRA |
| C3orf67 | TTC22 |
| C4orf47 | CYP2C18 |
| C5AR2 | AVPI1 |
| C5orf46 | CIDEB |
| C6orf163 | ID4 |
| C6orf164 | BASP1 |
| C6orf195 | EIF4EBP3 |
| C8G | APOH |
| C8orf87 | TFR2 |
| C9orf116 | FRRS1 |
| C9orf163 | TGFBR3L |
| C9orf57 | CES1 |
| C9orf66 | AKR1C4 |
| CA12 | MGST1 |
| CA5A | GRTP1 |
| CA5B | GJB1 |
| CA8 | SRXN1 |
| CA9 | MST1 |
| CABYR | APOC1 |
| CACNA1C | MYO7B |
| CACNA1D | UGDH |
| CACNB2 | MTRR |
| CACNB4 | AMBP |
| CALCA | LDHD |
| CALCB | NR1I2 |
| CAMK2N2 | GLYCTK |
| CAP2 | AKR1C1 |
| CAPN11 | ESPN |
| CATSPERB | CHST13 |
| CBX2 | METTL7A |
| CBX8 | A1CF |
| CC2D2B | KCNE3 |
| CCDC102B | KLF15 |
| CCDC113 | AMT |
| CCDC13 | SGK2 |
| CCDC136 | SLC1A2 |
| CCDC169 | HSD11B2 |

| | |
|---|---|
| CCDC170 | UBA7 |
| CCDC177 | EPAS1 |
| CCDC64 | GPAM |
| CCDC78 | TRIM15 |
| CCDC80 | NAT6 |
| CCDC85A | CDH16 |
| CCDC88C | ELMO3 |
| CCNA2 | BRD4 |
| CCNB1 | KCNQ1 |
| CCNB2 | LRRC8C |
| CCNE1 | NDRG2 |
| CCNF | GDF7 |
| CCNO | C2orf72 |
| CCR8 | HAAO |
| CCRN4L | ZFP36 |
| CCT3 | AGFG2 |
| CCT5 | NR1H3 |
| CCT6A | EPB41L4B |
| CD109 | LRIG3 |
| CD200 | AMN |
| CD248 | BHMT2 |
| CD34 | ARHGAP4 |
| CD46 | ZNF331 |
| CDC20 | COQ8A |
| CDC20B | NR0B1 |
| CDC25C | SAT1 |
| CDC45 | SHROOM1 |
| CDC6 | UGT1A3 |
| CDCA2 | PECR |
| CDCA5 | CGN |
| CDCA7 | |
| CDH11 | |
| CDH13 | |
| CDH17 | |
| CDH24 | |
| CDH6 | |
| CDK1 | |
| CDK6 | |
| CDKN2A | |
| CDKN2B | |
| CDKN3 | |
| CDX1 | |
| CDYL2 | |

| |
|---|
| CELSR3 |
| CEND1 |
| CENPF |
| CENPI |
| CENPK |
| CENPL |
| CENPO |
| CEP128 |
| CEP152 |
| CEP55 |
| CERCAM |
| CERKL |
| CGA |
| CGREF1 |
| CHAC1 |
| CHCHD3 |
| CHEK1 |
| CHI3L1 |
| CHML |
| CHN1 |
| CHRDL2 |
| CHRNA5 |
| CHST10 |
| CHST6 |
| CHST8 |
| CIAPIN1 |
| CIDEC |
| CKB |
| CKS2 |
| CLCF1 |
| CLDN5 |
| CLEC18B |
| CLGN |
| CLIC5 |
| CLIP4 |
| CLSPN |
| CMTM4 |
| CNNM2 |
| COL10A1 |
| COL11A1 |
| COL15A1 |
| COL17A1 |
| COL1A1 |

| |
|---|
| COL1A2 |
| COL22A1 |
| COL4A1 |
| COL4A2 |
| COL5A1 |
| COL5A2 |
| COL5A3 |
| COL6A3 |
| COL6A5 |
| COL8A1 |
| COL9A1 |
| COMP |
| COQ9 |
| CORIN |
| CORO2A |
| COX4I2 |
| CPA6 |
| CPD |
| CPE |
| CPLX1 |
| CPLX2 |
| CPNE5 |
| CPNE7 |
| CPS1 |
| CPXM1 |
| CRCP |
| CREB3L1 |
| CREG2 |
| CRLF2 |
| CRMP1 |
| CRP |
| CRYM |
| CSGALNACT1 |
| CSMD2 |
| CSNK2A2 |
| CSPG4 |
| CTD-2600O9.1 |
| CTH |
| CTHRC1 |
| CXCL16 |
| CXorf36 |
| CYBA |
| CYCS |

| |
|---|
| CYP17A1 |
| CYP19A1 |
| CYP1B1 |
| CYP21A2 |
| CYP27B1 |
| CYSTM1 |
| DAB2 |
| DAGLA |
| DCAF4L1 |
| DCDC2 |
| DCLK2 |
| DCX |
| DDIT4 |
| DDIT4L |
| DEFB1 |
| DEFB132 |
| DEPDC1 |
| DFNA5 |
| DGKI |
| DHRS7 |
| DIAPH3 |
| DIO2 |
| DKK1 |
| DKK2 |
| DKK4 |
| DLGAP4 |
| DLGAP5 |
| DNAAF1 |
| DNAAF3 |
| DNAH17 |
| DNAH5 |
| DNAH7 |
| DNAJC6 |
| DNER |
| DNM3 |
| DOCK5 |
| DOK4 |
| DOK5 |
| DPCD |
| DPEP1 |
| DPP10 |
| DPY19L1 |
| DPYSL4 |

| |
|---|
| DSEL |
| DTL |
| DTNA |
| DUSP4 |
| DUSP8 |
| DYDC2 |
| E2F1 |
| E2F3 |
| E2F5 |
| E2F7 |
| E2F8 |
| EARS2 |
| EBF1 |
| EBF2 |
| EBF3 |
| ECEL1 |
| ECT2 |
| EDA2R |
| EDIL3 |
| EEF1A2 |
| EFCAB12 |
| EFEMP1 |
| EFNA3 |
| EFNA5 |
| EGF |
| EGFL6 |
| EGLN3 |
| EHD2 |
| EIF4G3 |
| ELOVL3 |
| ELOVL7 |
| EMC7 |
| EME1 |
| ENDOD1 |
| ENO1 |
| ENOX1 |
| ENTPD1 |
| EPB41L1 |
| EPDR1 |
| EPHA10 |
| EPHX4 |
| EPS8L3 |
| ERBB2 |

| |
|---|
| ERC2 |
| ERCC6L |
| EREG |
| ERMP1 |
| ESCO2 |
| ESM1 |
| ESRRG |
| ETV1 |
| ETV4 |
| EVA1C |
| EVC |
| EXO1 |
| EYS |
| EZH2 |
| EZR |
| F13A1 |
| F2RL1 |
| F2RL3 |
| FA2H |
| FAHD2B |
| FAM101A |
| FAM111B |
| FAM115C |
| FAM127A |
| FAM127B |
| FAM127C |
| FAM132B |
| FAM135A |
| FAM155A |
| FAM155B |
| FAM167A |
| FAM169B |
| FAM177B |
| FAM179A |
| FAM188B |
| FAM189B |
| FAM196A |
| FAM19A5 |
| FAM227A |
| FAM64A |
| FAM71F2 |
| FAM78B |
| FAM81A |

| |
|---|
| FAM83A |
| FAM86A |
| FAM86B1 |
| FANCI |
| FAP |
| FASTKD1 |
| FAT1 |
| FBN1 |
| FBXL18 |
| FBXL8 |
| FBXO25 |
| FBXO36 |
| FBXO39 |
| FGF7 |
| FHDC1 |
| FKBP11 |
| FKBP14 |
| FKBP1C |
| FLAD1 |
| FLVCR1 |
| FMNL2 |
| FMNL3 |
| FNDC1 |
| FNDC3A |
| FOXC1 |
| FOXF2 |
| FOXL1 |
| FOXM1 |
| FOXQ1 |
| FOXS1 |
| FRRS1L |
| FRZB |
| FSD1L |
| FST |
| FSTL4 |
| FTO |
| FURIN |
| FZD10 |
| FZD2 |
| FZD6 |
| FZD8 |
| G6PD |
| GABRB3 |

| |
|---|
| GABRD |
| GABRE |
| GABRQ |
| GAD1 |
| GAL3ST4 |
| GALNT10 |
| GALNT12 |
| GALNT5 |
| GALNT7 |
| GALNTL6 |
| GALR2 |
| GAPDH |
| GAREML |
| GARS |
| GAS8 |
| GATSL2 |
| GCNT3 |
| GDA |
| GDF15 |
| GGN |
| GHRHR |
| GINS1 |
| GJA5 |
| GJC1 |
| GK |
| GLA |
| GLCE |
| GLIS3 |
| GLP1R |
| GLRB |
| GLRX |
| GLS |
| GMDS |
| GNA12 |
| GNAL |
| GNAZ |
| GNB5 |
| GNG4 |
| GOLGA6L9 |
| GOLM1 |
| GOLT1B |
| GOT1 |
| GPATCH4 |

| |
|---|
| GPC2 |
| GPD1L |
| GPI |
| GPR1 |
| GPR107 |
| GPR150 |
| GPR37 |
| GPR4 |
| GPR56 |
| GPR63 |
| GPR64 |
| GPR68 |
| GPR97 |
| GPRIN1 |
| GPRIN3 |
| GPSM2 |
| GPX8 |
| GRAMD1A |
| GRAMD1B |
| GRAMD4 |
| GREM2 |
| GRIA3 |
| GRK7 |
| GRM5 |
| GRM7 |
| GSDMC |
| GSN |
| GTF2E1 |
| GTF2IRD1 |
| GTPBP4 |
| GUCY2C |
| GULP1 |
| GXYLT2 |
| H2AFY2 |
| HAGHL |
| HAPLN1 |
| HAS3 |
| HAVCR1 |
| HCN1 |
| HEATR2 |
| HEG1 |
| HELLS |
| HES2 |

| |
|---|
| HES4 |
| HEYL |
| HHATL |
| HHIPL2 |
| HHLA3 |
| HIGD1B |
| HIST1H1A |
| HIST1H1C |
| HIST1H1T |
| HIST1H2AA |
| HIST1H2AB |
| HIST1H2AC |
| HIST1H2AG |
| HIST1H2AH |
| HIST1H2AI |
| HIST1H2AK |
| HIST1H2BB |
| HIST1H2BG |
| HIST1H2BJ |
| HIST1H2BK |
| HIST1H2BN |
| HIST1H3A |
| HIST1H3D |
| HIST1H4I |
| HIST2H3D |
| HJURP |
| HK2 |
| HKDC1 |
| HMGB3 |
| HNF1B |
| HOMER1 |
| HOPX |
| HOXD1 |
| HOXD3 |
| HOXD9 |
| HPD |
| HSF4 |
| HSP90AA1 |
| HSP90AB1 |
| HSPA12A |
| HSPA12B |
| HSPA4L |
| HSPA5 |

| |
|---|
| HSPB6 |
| HSPH1 |
| HTATIP2 |
| HTR2A |
| HTRA1 |
| HYOU1 |
| IARS |
| ICAM5 |
| IER3 |
| IER5L |
| IGDCC4 |
| IGF2BP3 |
| IGFBPL1 |
| IGSF9B |
| IL17D |
| IL2RA |
| IL31RA |
| IL32 |
| INPP4B |
| INSL4 |
| INTS9 |
| INTU |
| IQCA1 |
| IQCD |
| IQCE |
| IQCK |
| IQGAP3 |
| IRAK1 |
| IRF4 |
| IRX3 |
| IRX5 |
| ISG15 |
| ISG20 |
| ITGA11 |
| ITGA2 |
| ITGA6 |
| ITGA7 |
| ITGAV |
| ITGB4 |
| ITPKA |
| JAG1 |
| JAG2 |
| JPH1 |

| |
|---|
| KAAG1 |
| KAL1 |
| KCNE1L |
| KCNE4 |
| KCNF1 |
| KCNJ5 |
| KCNJ6 |
| KCNK9 |
| KCNN3 |
| KCNQ3 |
| KCNU1 |
| KDELR3 |
| KIAA0100 |
| KIAA0319 |
| KIAA0556 |
| KIAA0895L |
| KIAA1024 |
| KIAA1199 |
| KIAA1211L |
| KIAA1244 |
| KIAA1324 |
| KIAA1462 |
| KIAA1522 |
| KIAA1549 |
| KIAA1549L |
| KIF14 |
| KIF18B |
| KIF20A |
| KIF21B |
| KIF23 |
| KIF24 |
| KIF26B |
| KIF2C |
| KIF3A |
| KIF4A |
| KIF5A |
| KLC2 |
| KLF5 |
| KLHDC7B |
| KLHL21 |
| KLHL29 |
| KNDC1 |
| KPNA2 |

| |
|---|
| KRT222 |
| KRT7 |
| KRT81 |
| KRT86 |
| KSR1 |
| KYNU |
| LAMA1 |
| LAMA3 |
| LAMA4 |
| LAMB4 |
| LAMC1 |
| LANCL1 |
| LANCL3 |
| LAPTM4B |
| LBP |
| LCN2 |
| LDHB |
| LEF1 |
| LEP |
| LETM2 |
| LGALS3 |
| LGI2 |
| LHCGR |
| LHFPL5 |
| LHX6 |
| LIF |
| LIMCH1 |
| LIMK1 |
| LIMK2 |
| LIPH |
| LITAF |
| LOH12CR1 |
| LONRF2 |
| LOX |
| LOXL2 |
| LPAR3 |
| LPAR4 |
| LPCAT1 |
| LPL |
| LPPR4 |
| LRP8 |
| LRRC1 |
| LRRC16A |

| |
|---|
| LRRC37A3 |
| LRRC69 |
| LRRC73 |
| LRRC8E |
| LSAMP |
| LTBP2 |
| LUZP2 |
| LYPD1 |
| LZTS1 |
| MAB21L3 |
| MAFK |
| MALL |
| MANF |
| MAP1A |
| MAP1B |
| MAP1LC3B |
| MAP1LC3B2 |
| MAP3K9 |
| MAP9 |
| MAPK12 |
| MAPT |
| MARS2 |
| MATN3 |
| MBOAT4 |
| MCAM |
| MCF2L2 |
| MCM10 |
| MCM4 |
| MCTP2 |
| MECOM |
| MED9 |
| MELK |
| MEX3B |
| MFSD6 |
| MGAT3 |
| MGAT5B |
| MGC4294 |
| MID1 |
| MKI67 |
| MLEC |
| MMP1 |
| MMP10 |
| MMP11 |

| |
|---|
| MMP14 |
| MMP2 |
| MN1 |
| MND1 |
| MPHOSPH6 |
| MPP3 |
| MPP4 |
| MPV17 |
| MPV17L2 |
| MPZ |
| MRAP |
| MRAP2 |
| MRAS |
| MRPS12 |
| MRVI1 |
| MST1R |
| MT-ATP8 |
| MT-ND5 |
| MT-ND6 |
| MT3 |
| MTFR2 |
| MTHFD1L |
| MTRNR2L2 |
| MUC13 |
| MUC3A |
| MUC5B |
| MURC |
| MVP |
| MYBPHL |
| MYCN |
| MYEF2 |
| MYH4 |
| MYO1E |
| MYO5C |
| MYOF |
| MYOM3 |
| MYRF |
| N4BP3 |
| NAV1 |
| NAV2 |
| NBEA |
| NCAPG |
| NCR3LG1 |

| |
|---|
| NDUFA4L2 |
| NEB |
| NEIL3 |
| NEK2 |
| NETO2 |
| NHS |
| NIPAL2 |
| NKX1-2 |
| NKX2-3 |
| NME1 |
| NMNAT2 |
| NOL3 |
| NOMO1 |
| NOMO2 |
| NOMO3 |
| NOTCH3 |
| NOV |
| NOVA1 |
| NOX1 |
| NOX4 |
| NPAS2 |
| NPCDR1 |
| NPFFR2 |
| NPNT |
| NPTX2 |
| NPTXR |
| NPY5R |
| NQO1 |
| NR4A2 |
| NRG2 |
| NRIP2 |
| NRXN3 |
| NT5C3A |
| NT5DC2 |
| NTM |
| NTS |
| NUF2 |
| NXPH4 |
| OAT |
| OGDH |
| OGFOD1 |
| OLAH |
| OLFML2A |

| |
|---|
| OLFML2B |
| OPHN1 |
| OPN3 |
| OR1F1 |
| OR2A7 |
| OR2AG2 |
| OR2B6 |
| OR2D2 |
| OR51E1 |
| OR51E2 |
| OR6A2 |
| ORC1 |
| ORC6 |
| OSBPL3 |
| OSBPL9 |
| OSMR |
| OSR1 |
| OSR2 |
| OTOG |
| OTUB2 |
| OTUD7A |
| OTX1 |
| OXCT1 |
| P4HA2 |
| PAEP |
| PAK3 |
| PALLD |
| PAPPA |
| PAQR4 |
| PAQR5 |
| PARM1 |
| PARPBP |
| PBK |
| PBX1 |
| PCDH17 |
| PCDHB10 |
| PCDHB11 |
| PCDHB13 |
| PCDHB8 |
| PCDHGA1 |
| PCDHGA12 |
| PCDHGA4 |
| PCDHGA5 |

| |
|---|
| PCDHGA7 |
| PCDHGA8 |
| PCDHGB1 |
| PCDHGB2 |
| PCDHGC5 |
| PCDP1 |
| PCED1B |
| PCM1 |
| PCNT |
| PCNXL2 |
| PCSK1 |
| PDE10A |
| PDE1C |
| PDE3A |
| PDE3B |
| PDE4A |
| PDE4B |
| PDE4D |
| PDF |
| PDGFA |
| PDGFRB |
| PDGFRL |
| PDK1 |
| PDLIM7 |
| PDX1 |
| PDZD2 |
| PFKFB2 |
| PFKM |
| PFKP |
| PHEX |
| PHF17 |
| PHLDA2 |
| PHLDA3 |
| PHPT1 |
| PIAS3 |
| PIGA |
| PIP4K2C |
| PITPNM1 |
| PITX1 |
| PKP1 |
| PLA2G2A |
| PLA2G2C |
| PLA2G4E |

| |
|---|
| PLCB4 |
| PLCD3 |
| PLCE1 |
| PLCH1 |
| PLEKHA8 |
| PLEKHG2 |
| PLEKHH1 |
| PLEKHN1 |
| PLK1 |
| PLOD3 |
| PLP2 |
| PLVAP |
| PLXDC1 |
| PM20D2 |
| PMEPA1 |
| PMFBP1 |
| PNMA1 |
| PODNL1 |
| PODXL |
| POLN |
| POLR1A |
| POLR2C |
| POLR3G |
| PON2 |
| POSTN |
| POU5F1 |
| PPA1 |
| PPARGC1A |
| PPAT |
| PPIA |
| PPP1R13L |
| PPP1R36 |
| PPP1R3D |
| PPP1R3G |
| PPP2R2C |
| PRAMEF10 |
| PRAMEF2 |
| PRAMEF4 |
| PRC1 |
| PRDM6 |
| PRDM7 |
| PRKACA |
| PRND |

| |
|---|
| PRODH |
| PRR15 |
| PRR16 |
| PRR26 |
| PRRG3 |
| PRRX1 |
| PRSS27 |
| PSMD14 |
| PSME3 |
| PSORS1C1 |
| PTCHD4 |
| PTGFR |
| PTGFRN |
| PTP4A3 |
| PTPDC1 |
| PTPLA |
| PTPN14 |
| PTPN5 |
| PTPRM |
| PTPRR |
| PTPRU |
| PVRL1 |
| PYCR1 |
| PYGB |
| QRFPR |
| RAB3A |
| RAB3B |
| RAB6B |
| RACGAP1 |
| RAET1E |
| RANBP17 |
| RAP1GAP |
| RARRES1 |
| RASD1 |
| RASD2 |
| RASEF |
| RASGEF1A |
| RASGRF2 |
| RASL11B |
| RASL12 |
| RASSF6 |
| RASSF9 |
| RBM20 |

| |
|---|
| RBM24 |
| RBM44 |
| RBPMS |
| RECQL4 |
| RELL2 |
| RFX8 |
| RGAG4 |
| RGCC |
| RGS17 |
| RGS5 |
| RGS6 |
| RGS9 |
| RHBDD2 |
| RHOBTB1 |
| RHOBTB2 |
| RHOF |
| RHOQ |
| RHPN1 |
| RMI2 |
| RNF157 |
| RNFT2 |
| ROBO1 |
| ROR2 |
| RP1-102H19.8 |
| RP11-17M16.1 |
| RP11-212D19.4 |
| RP11-248J23.6 |
| RP11-366L20.2 |
| RP11-796G6.2 |
| RPGR |
| RPGRIP1L |
| RPP40 |
| RPRML |
| RPS6KA2 |
| RPS6KL1 |
| RRM2 |
| RRS1 |
| RUNX1 |
| S100A3 |
| S100P |
| SAPCD2 |
| SATB2 |
| SCG2 |

| |
|---|
| SCIN |
| SCML4 |
| SCN4A |
| SDCBP2 |
| SDIM1 |
| SDSL |
| SEMA3F |
| SEMA3G |
| SEMA5B |
| SERHL2 |
| SERPINB3 |
| SERTAD4 |
| SEZ6L2 |
| SFMBT2 |
| SFN |
| SFRP2 |
| SFRP4 |
| SGIP1 |
| SGPP2 |
| SH3PXD2B |
| SH3RF3 |
| SHC1 |
| SHCBP1 |
| SHISA2 |
| SHROOM4 |
| SIAE |
| SIX1 |
| SIX4 |
| SKA1 |
| SLC16A11 |
| SLC16A14 |
| SLC22A11 |
| SLC22A12 |
| SLC22A15 |
| SLC22A23 |
| SLC22A5 |
| SLC25A12 |
| SLC25A15 |
| SLC25A24 |
| SLC26A2 |
| SLC26A7 |
| SLC26A9 |
| SLC27A4 |

| |
|---|
| SLC2A1 |
| SLC2A5 |
| SLC35B4 |
| SLC35C1 |
| SLC35E4 |
| SLC35G2 |
| SLC36A1 |
| SLC38A1 |
| SLC44A3 |
| SLC45A1 |
| SLC4A11 |
| SLC52A3 |
| SLC5A6 |
| SLC6A11 |
| SLC6A17 |
| SLC6A3 |
| SLC6A6 |
| SLC6A8 |
| SLC6A9 |
| SLC7A1 |
| SLC7A11 |
| SLC7A2 |
| SLC7A5 |
| SLC7A6 |
| SLCO1C1 |
| SLCO2A1 |
| SLCO5A1 |
| SLIT2 |
| SLITRK4 |
| SMAD2 |
| SMCO2 |
| SMKR1 |
| SMOC2 |
| SMOX |
| SNAP25 |
| SNPH |
| SOGA1 |
| SORT1 |
| SOX11 |
| SOX12 |
| SOX21 |
| SOX4 |
| SP140 |

| |
|---|
| SP6 |
| SPA17 |
| SPATA17 |
| SPATC1L |
| SPATS2 |
| SPECC1 |
| SPINK1 |
| SPIRE1 |
| SPOCD1 |
| SPOCK1 |
| SPRED3 |
| SPTBN5 |
| SPTSSA |
| SRGAP1 |
| SRM |
| SRPX2 |
| SRRM3 |
| SRSF12 |
| ST6GALNAC4 |
| ST8SIA5 |
| STAMBPL1 |
| STC1 |
| STC2 |
| STEAP1 |
| STEAP1B |
| STK32B |
| STRA6 |
| STRIP2 |
| STS |
| STXBP1 |
| STXBP4 |
| STXBP5 |
| SULF1 |
| SULF2 |
| SULT1C2 |
| SULT2B1 |
| SULT4A1 |
| SYNDIG1 |
| SYT13 |
| TAF4B |
| TAF6 |
| TAF7L |
| TAGLN2 |

| |
|---|
| TANC2 |
| TAS2R3 |
| TAS2R4 |
| TAS2R5 |
| TAX1BP3 |
| TBC1D16 |
| TBC1D30 |
| TBC1D31 |
| TC2N |
| TCF23 |
| TCN1 |
| TCTN3 |
| TEAD4 |
| TENM4 |
| TESC |
| TEX9 |
| TG |
| TGFB2 |
| TGFBR1 |
| TGIF1 |
| TGM2 |
| TGM3 |
| THBS2 |
| THBS4 |
| THSD7A |
| THY1 |
| TICAM2 |
| TICRR |
| TIMP1 |
| TK1 |
| TLCD1 |
| TLCD2 |
| TLDC1 |
| TLE6 |
| TLL2 |
| TM4SF5 |
| TMC1 |
| TMC5 |
| TMC7 |
| TMED3 |
| TMEM104 |
| TMEM108 |
| TMEM119 |

| |
|---|
| TMEM132A |
| TMEM136 |
| TMEM144 |
| TMEM145 |
| TMEM156 |
| TMEM163 |
| TMEM165 |
| TMEM178B |
| TMEM2 |
| TMEM200A |
| TMEM217 |
| TMEM245 |
| TMEM246 |
| TMEM45A |
| TMEM55A |
| TMEM59L |
| TMOD1 |
| TMTC2 |
| TNFRSF10A |
| TNFRSF12A |
| TNFRSF21 |
| TNFRSF4 |
| TNFSF4 |
| TNRC6C |
| TOMM40L |
| TOP2A |
| TOX |
| TPBG |
| TPBGL |
| TPM4 |
| TRAF2 |
| TRIM31 |
| TRIM40 |
| TRIM46 |
| TRIM59 |
| TRIM7 |
| TRIM8 |
| TRIP13 |
| TRPC4 |
| TRPC6 |
| TRPM2 |
| TRPS1 |
| TSHZ2 |

| |
|---|
| TSPAN13 |
| TTC26 |
| TTC39A |
| TTK |
| TTL |
| TTYH2 |
| TUBA4A |
| TXNDC17 |
| TXNRD1 |
| TYMS |
| UBE2C |
| UBE2T |
| UBFD1 |
| UBXN10 |
| UCHL1 |
| UCN |
| UGT2B11 |
| ULBP1 |
| ULBP2 |
| ULBP3 |
| UNC119B |
| UNC13A |
| UNC13B |
| UNC5B |
| UNC5D |
| URB1 |
| USB1 |
| USH1C |
| USP49 |
| USP54 |
| UTS2B |
| VARS |
| VASH2 |
| VAX2 |
| VCAN |
| VLDLR |
| VSIG1 |
| VSX1 |
| VWA7 |
| VWF |
| WDR13 |
| WIPF3 |
| WNT16 |

| |
|---|
| WWC1 |
| XKR3 |
| XKR6 |
| XKRX |
| XPR1 |
| XRCC2 |
| YIPF6 |
| YWHAG |
| ZBTB38 |
| ZC3H12B |
| ZFHX3 |
| ZFP69B |
| ZHX1-C8ORF76 |
| ZIC2 |
| ZMAT3 |
| ZMYND15 |
| ZNF154 |
| ZNF215 |
| ZNF23 |
| ZNF233 |
| ZNF365 |
| ZNF382 |
| ZNF385D |
| ZNF57 |
| ZNF683 |
| ZNF695 |
| ZNF703 |
| ZNF724P |
| ZNF827 |
| ZP3 |
| ZSCAN5A |
| ZSCAN9 |
| ZWINT |
| ZXDB |

**Supplementary Table 4B.** Consensus PRKACA shRNA modulated genes

| log2FC_Neg_Simon | Overlap-All3_shRNA-POS | Overlap | Hallmark |
|---|---|---|---|
| MARCH2 | IGLON5 | ADAMTSL2 | EPITHELIAL_MESENCHYMAL_TRANSITION |
| SEPT4 | ETNK2 | AMOTL2 | ESTROGEN_RESPONSE_LATE |
| A1BG | GJB3 | ANXA3 | COAGULATION |
| A1CF | CASC10 | ARVCF | TNFA_SIGNALING_VIA_NFKB |
| AADAC | IGSF9 | CAPN5 | CHOLESTEROL_HOMEOSTASIS |
| AADAT | IFIT1 | CASC10 | IL6_JAK_STAT3_SIGNALING |
| ABAT | KRT19 | CD9 | |
| ABCA13 | ECM1 | CDH2 | |
| ABCA6 | TMEM63C | COL18A1 | |
| ABCA9 | SKIDA1 | CXADR | |
| ABCB11 | WNT11 | CYP2S1 | |
| ABCG2 | MT1G | EBF4 | |
| ABCG5 | ADAMTSL2 | ECM1 | |
| ABCG8 | TRIB1 | ETNK2 | |
| ABHD1 | HMGCS2 | GJB3 | |
| ABHD15 | ANXA3 | GPR162 | |
| ABHD5 | NTN4 | HMGCS2 | |
| AC007405.2 | COL18A1 | IFIT1 | |
| AC010368.2 | NGFR | IGLON5 | |
| AC011484.1 | CDH2 | IGSF9 | |
| AC011841.1 | CYP2S1 | JUN | |
| AC018755.1 | JUN | KRT19 | |
| AC021860.1 | RHOB | LIMD2 | |
| AC023590.1 | CAPN5 | MT1G | |
| AC103801.2 | ARVCF | NGFR | |
| AC104667.3 | SDK2 | NTN4 | |
| AC104809.3 | GPR162 | NUDT4 | |
| AC110781.3 | VPS37D | RGL1 | |
| ACAA1 | CXADR | RHOB | |
| ACAA2 | TINAGL1 | SDK2 | |
| ACACB | AMOTL2 | SHH | |
| ACADL | RGL1 | SKIDA1 | |
| ACADM | SHH | TINAGL1 | |
| ACADS | LIMD2 | TMEM63C | |
| ACADSB | TP53INP1 | TP53INP1 | |
| ACAT1 | NUDT4 | TRIB1 | |
| ACAT2 | YPEL1 | VPS37D | |
| ACBD4 | CD9 | WNT11 | |
| ACE2 | EBF4 | YPEL1 | |
| ACKR4 | SRPX2 | | |
| ACOT1 | FOXC2 | | |
| ACOT12 | SGPP1 | | |
| ACOT13 | PAPLN | | |
| ACOT2 | CTXN1 | | |
| ACOT4 | RIMKLA | | |
| ACOX2 | ATXN1 | | |
| ACP5 | SORT1 | | |
| ACSF2 | SLC4A11 | | |
| ACSL1 | SEMA4F | | |
| ACSL5 | MMP24 | | |
| ACSL6 | DBNDD1 | | |
| ACSM2A | FAM84A | | |
| ACSM2B | BAMBI | | |
| ACSM3 | DTNA | | |
| ACSM5 | CITED2 | | |

| | |
|---|---|
| ACSS2 | MGAT5 |
| ACY3 | TGFA |
| ADAM11 | DNAJB2 |
| ADAMTS13 | THBS1 |
| ADAMTSL2 | TRIM29 |
| ADAMTSL3 | OSBP2 |
| ADAMTSL4 | IMPDH1 |
| ADAP1 | GATA6 |
| ADCY1 | EMP3 |
| ADCY10 | FLVCR2 |
| ADCYAP1R1 | FAM171A2 |
| ADH1A | PTPN21 |
| ADH1B | RHOQ |
| ADH4 | RTN2 |
| ADH6 | CCDC148 |
| ADH7 | ZCCHC12 |
| ADI1 | ABHD2 |
| ADIRF | PCDHGA9 |
| ADM | PIK3R3 |
| ADRA1A | PCDHGC3 |
| ADRA1B | EEPD1 |
| ADRA2B | PCDHGB2 |
| ADRB1 | GOLM1 |
| ADRB2 | SLC35E4 |
| ADTRP | FLG |
| AFF3 | ACTN1 |
| AFM | ABTB2 |
| AGBL2 | MFSD6 |
| AGL | MPP2 |
| AGMAT | PLCB1 |
| AGMO | PDLIM7 |
| AGPAT2 | CAPS |
| AGTR1 | RCAN3 |
| AGXT | HSPG2 |
| AGXT2 | KRT80 |
| AHSG | PCDHB8 |
| AIG1 | LRRN2 |
| AJAP1 | DNAJC18 |
| AKAP3 | ADAMTS14 |
| AKR1A1 | C15orf52 |
| AKR1C4 | OSR2 |
| AKR1D1 | PPL |
| AL078585.1 | NPTXR |
| AL590714.1 | IQSEC2 |
| ALAD | KLF6 |
| ALAS1 | CHRNA3 |
| ALAS2 | FLNA |
| ALB | PCDHA4 |
| ALDH1L1 | DPYSL2 |
| ALDH2 | SLC29A4 |
| ALDH5A1 | SHC2 |
| ALDH6A1 | SLFN12 |
| ALDH7A1 | SH3RF1 |
| ALDH9A1 | LTBP4 |
| ALDOC | DNER |
| ALLC | TLE1 |
| ALPL | MAP1B |
| AMACR | TMEM132A |

| | |
|---|---|
| AMBP | EMID1 |
| AMHR2 | RASSF4 |
| AMN | REEP1 |
| AMOTL2 | DUSP6 |
| AMT | SUGCT |
| ANG | BCAM |
| ANGPTL6 | CERCAM |
| ANGPTL7 | FZD6 |
| ANK3 | MAPK8IP2 |
| ANKRD20A3 | ZNF608 |
| ANKRD55 | TUBB3 |
| ANO1 | ABHD6 |
| ANPEP | PCDHB11 |
| ANXA10 | VCL |
| ANXA3 | VWA5B2 |
| ANXA9 | SMURF2 |
| AOC1 | SULF2 |
| AP1M2 | STMN3 |
| APBA1 | NAP1L3 |
| APH1A | DUSP8 |
| APOA1 | CACNA1S |
| APOA2 | FAM13B |
| APOA5 | FLRT3 |
| APOBEC3A | CCDC74A |
| APOC1 | PCDHB14 |
| APOC3 | RAB6B |
| APOF | INHBB |
| APOH | SHROOM3 |
| APOL1 | FOXO6 |
| APOL6 | CPNE2 |
| APOM | ABCC4 |
| AQP11 | C12orf75 |
| AQP7 | FBXL2 |
| AQP9 | PCDHB15 |
| AR | SERINC2 |
| ARC | WNT10A |
| ARHGAP20 | SOX4 |
| ARHGEF10L | TNFRSF12A |
| ARHGEF26 | KIRREL |
| ARHGEF38 | SPON2 |
| ARHGEF40 | ORAI2 |
| ARID3C | CMTM3 |
| ARID5A | TMEM158 |
| ARL4D | UNC5B |
| ARMC3 | ZNF185 |
| ARMC5 | PCDHB2 |
| ARRDC3 | PCDHB7 |
| ARSF | GLI2 |
| ART3 | MBOAT2 |
| ART4 | SHANK1 |
| ART5 | CRISPLD2 |
| ARVCF | TMEM173 |
| ASB4 | PCDHB6 |
| ASB9 | ZNF423 |
| ASCL2 | TANC2 |
| ASGR1 | CDK5R1 |
| ASMTL | SERPINE2 |
| ASPA | TNFRSF21 |

| | |
|---|---|
| ASPDH | SALL2 |
| ASTL | PDGFA |
| ASTN1 | RGCC |
| ASXL3 | SCRN1 |
| ATAD3C | RPS6KA2 |
| ATF3 | MEIS3 |
| ATF5 | TMSB4X |
| ATF7IP2 | ASAP3 |
| ATHL1 | FAM110B |
| ATOH8 | TPM1 |
| ATP11C | APLP1 |
| ATP13A4 | MAP2 |
| ATP7B | DACT1 |
| ATRNL1 | REEP2 |
| AUTS2 | RIMS3 |
| AVPR1A | PRRT4 |
| AXL | PIK3IP1 |
| AZGP1 | PROCR |
| AZU1 | PLAT |
| B3GAT1 | MYO5A |
| B3GAT2 | MFAP2 |
| B3GNT8 | PHLDB1 |
| B4GALNT3 | ADAMTS15 |
| BAALC | VIM |
| BACH2 | LRRC75A |
| BAI3 | IGSF9B |
| BAIAP2 | PTPN14 |
| BAIAP3 | HEG1 |
| BBOX1 | MAPK8IP1 |
| BCHE | MEX3B |
| BCL2A1 | AHNAK |
| BCL6 | LRRC49 |
| BCO2 | RCOR2 |
| BDH1 | IL17RD |
| BEND4 | TUBB6 |
| BEND7 | COL6A1 |
| BEX1 | MICAL1 |
| BEX4 | LRP12 |
| BHMT | SEPT5 |
| BHMT2 | NKD2 |
| BLK | KLF9 |
| BLNK | TUBB2A |
| BMP10 | CACNB3 |
| BMP3 | PHLDA3 |
| BMP5 | ADGRB2 |
| BMPER | MT1X |
| BNC1 | GAS6 |
| BOK | NKAIN1 |
| BPHL | TPBG |
| BPI | GLCCI1 |
| BRINP2 | TNNT1 |
| BRIP1 | PIFO |
| BTD | ADORA1 |
| BTNL8 | CRIP2 |
| BZRAP1 | CHST3 |
| C10orf11 | TBC1D9 |
| C10orf25 | GALNT5 |
| C10orf67 | MRAS |

| | |
|---|---|
| C11orf31 | EFR3B |
| C11orf35 | UGT8 |
| C11orf54 | SEMA7A |
| C11orf65 | EGR1 |
| C11orf71 | DLG3 |
| C11orf85 | MEX3A |
| C12orf42 | ATP2A3 |
| C12orf68 | PDLIM4 |
| C12orf74 | CDKN1C |
| C14orf164 | NDRG4 |
| C14orf180 | PLK2 |
| C14orf80 | FN1 |
| C15orf26 | EPPK1 |
| C15orf43 | TMEM255A |
| C16orf96 | MT2A |
| C19orf12 | CAMK1D |
| C19orf38 | PCDHB10 |
| C19orf66 | STK32B |
| C19orf71 | BMP7 |
| C19orf80 | TPPP3 |
| C1orf115 | ANXA6 |
| C1orf116 | GLB1L2 |
| C1orf162 | TIMP2 |
| C1orf168 | RUNX2 |
| C1orf172 | PDLIM1 |
| C1orf173 | TIMP3 |
| C1orf21 | PMAIP1 |
| C1orf210 | GJB4 |
| C1orf226 | MYL9 |
| C1orf53 | GJA3 |
| C1QB | MPZL2 |
| C1QC | TGFB2 |
| C1R | KRT20 |
| C1RL | PAK1 |
| C1S | KCNH3 |
| C2 | KCNN1 |
| C21orf2 | TAGLN3 |
| C21orf33 | ST8SIA2 |
| C21orf37 | MFGE8 |
| C21orf62 | GLIPR1 |
| C21orf67 | IGFL2 |
| C21orf88 | AMOT |
| C21orf90 | PAM |
| C21orf91 | LFNG |
| C2CD4B | KLK5 |
| C2orf40 | ANK1 |
| C2orf54 | NACAD |
| C2orf72 | ESM1 |
| C4BPA | WNT9A |
| C5AR1 | AHNAK2 |
| C5orf27 | GPR173 |
| C5orf49 | FILIP1L |
| C6 | B3GNT7 |
| C7orf10 | SYNGR3 |
| C8A | ANTXR2 |
| C8B | SLC34A2 |
| C8orf82 | SYT11 |
| C9 | LTBP1 |

| | |
|---|---|
| C9orf43 | GSN |
| C9orf72 | ANGPTL2 |
| C9orf96 | GPR161 |
| CA1 | ACTA2 |
| CA14 | FGF20 |
| CABLES1 | AHRR |
| CABP4 | PCSK5 |
| CACNA2D2 | HDAC5 |
| CADM2 | ZDHHC2 |
| CALN1 | CREB5 |
| CAMK2B | HAPLN3 |
| CAMK2N1 | CLIP3 |
| CAMK4 | CLSTN2 |
| CAND2 | SBK1 |
| CAPN12 | PNMA2 |
| CAPN5 | LOXL2 |
| CAPZA3 | EFNB3 |
| CARHSP1 | TMEM59L |
| CASC10 | GLIS3 |
| CASR | TMEFF1 |
| CAT | NPC2 |
| CBFA2T3 | GPR1 |
| CBLC | CALHM1 |
| CBLN4 | TMCC2 |
| CBR4 | SPARC |
| CBS | AMIGO2 |
| CBX7 | OLFML2A |
| CCBE1 | AQP1 |
| CCDC150 | CD8B |
| CCDC151 | ZDHHC22 |
| CCDC152 | SCARA3 |
| CCDC158 | ISM1 |
| CCDC178 | PDGFB |
| CCDC64B | ADAMTS7 |
| CCDC68 | CLIC5 |
| CCDC73 | LTBP2 |
| CCL14 | TUBA1A |
| CCL16 | MT1F |
| CCL2 | WNT6 |
| CCL23 | BMF |
| CCL24 | TMEM265 |
| CCL28 | KRTAP2-3 |
| CCL3 | KRTAP3-1 |
| CCL4L1 | |
| CCNB1IP1 | |
| CCR1 | |
| CCR9 | |
| CCT6B | |
| CD14 | |
| CD160 | |
| CD163 | |
| CD180 | |
| CD1C | |
| CD1D | |
| CD1E | |
| CD244 | |
| CD274 | |
| CD300A | |

| |
|---|
| CD300C |
| CD300E |
| CD300LB |
| CD300LG |
| CD302 |
| CD36 |
| CD4 |
| CD5L |
| CD81 |
| CD82 |
| CD83 |
| CD9 |
| CDH19 |
| CDH2 |
| CDH23 |
| CDH4 |
| CDHR5 |
| CDK18 |
| CDK3 |
| CEACAM3 |
| CEACAM4 |
| CEACAM6 |
| CEBPA |
| CEBPD |
| CECR2 |
| CENPV |
| CERS4 |
| CES1 |
| CES2 |
| CES3 |
| CES4A |
| CES5A |
| CETP |
| CFB |
| CFD |
| CFHR1 |
| CFHR2 |
| CFHR3 |
| CFHR4 |
| CFHR5 |
| CFL2 |
| CFP |
| CFTR |
| CGN |
| CGNL1 |
| CHAD |
| CHADL |
| CHDC2 |
| CHDH |
| CHN2 |
| CHRDL1 |
| CHRM2 |
| CHRNA4 |
| CHST13 |
| CHST4 |
| CHST7 |
| CIDEB |
| CILP |

| |
|---|
| CISH |
| CLDN1 |
| CLDN10 |
| CLDN14 |
| CLDN19 |
| CLDN3 |
| CLDN4 |
| CLEC1B |
| CLEC3B |
| CLEC4C |
| CLEC4D |
| CLEC4E |
| CLEC4G |
| CLEC4M |
| CLVS2 |
| CMA1 |
| CMBL |
| CMPK2 |
| CMTM2 |
| CMTM8 |
| CMYA5 |
| CNBD1 |
| CNDP1 |
| CNGA1 |
| CNKSR2 |
| CNPY3 |
| CNST |
| CNTLN |
| CNTN5 |
| COBLL1 |
| COL11A2 |
| COL18A1 |
| COL25A1 |
| COL27A1 |
| COL28A1 |
| COL6A6 |
| COLEC10 |
| COLEC11 |
| COMT |
| COQ10A |
| COX6A2 |
| CP |
| CPA3 |
| CPAMD8 |
| CPB2 |
| CPEB3 |
| CPED1 |
| CPN1 |
| CPN2 |
| CPNE6 |
| CR1 |
| CR1L |
| CRHBP |
| CRLS1 |
| CRYAA |
| CRYL1 |
| CSAD |
| CSF3 |

| |
|---|
| CSF3R |
| CSRNP1 |
| CTD-2014B16.3 |
| CTF1 |
| CTNNA3 |
| CTSG |
| CTSL |
| CTTNBP2 |
| CUX2 |
| CX3CL1 |
| CXADR |
| CXCL1 |
| CXCL12 |
| CXCL14 |
| CXCL2 |
| CXCL3 |
| CXCR1 |
| CXCR2 |
| CXorf22 |
| CXorf66 |
| CYB5A |
| CYP11A1 |
| CYP1A1 |
| CYP1A2 |
| CYP26A1 |
| CYP26B1 |
| CYP27A1 |
| CYP27C1 |
| CYP2A13 |
| CYP2A6 |
| CYP2A7 |
| CYP2B6 |
| CYP2C19 |
| CYP2C8 |
| CYP2C9 |
| CYP2D6 |
| CYP2J2 |
| CYP2S1 |
| CYP39A1 |
| CYP3A4 |
| CYP3A43 |
| CYP3A5 |
| CYP3A7 |
| CYP4A11 |
| CYP4A22 |
| CYP4F12 |
| CYP4F2 |
| CYP4F3 |
| CYP4V2 |
| CYP4X1 |
| CYP4Z1 |
| CYP7A1 |
| CYP7B1 |
| CYP8B1 |
| DAB1 |
| DACH1 |
| DAK |
| DAO |

| |
|---|
| DBH |
| DBI |
| DCAF11 |
| DCHS2 |
| DCT |
| DCXR |
| DDC |
| DDT |
| DDTL |
| DDX25 |
| DEFA3 |
| DEPDC7 |
| DGAT2 |
| DGCR6L |
| DHCR24 |
| DHCR7 |
| DHFR |
| DHODH |
| DHRS1 |
| DHRS12 |
| DHRS2 |
| DHRS4 |
| DHRS4L2 |
| DHTKD1 |
| DIO1 |
| DIRAS3 |
| DLEU7 |
| DLGAP2 |
| DMD |
| DMGDH |
| DMRTA1 |
| DNAH6 |
| DNAJC12 |
| DNAJC19 |
| DNALI1 |
| DNASE1L3 |
| DNMT3L |
| DOK2 |
| DOK6 |
| DPEP2 |
| DPEP3 |
| DPF3 |
| DPP4 |
| DPPA4 |
| DPT |
| DPYD |
| DPYS |
| DRD1 |
| DSC2 |
| DSCAM |
| DSG1 |
| DSG4 |
| DTX1 |
| DUSP10 |
| DUSP14 |
| DYNLRB2 |
| EBF4 |
| EBI3 |

| |
|---|
| EBP |
| EBPL |
| ECH1 |
| ECHDC2 |
| ECHDC3 |
| ECI2 |
| ECM1 |
| EDAR |
| EDN1 |
| EDNRB |
| EFCAB1 |
| EFCC1 |
| EFHD1 |
| EFNA2 |
| EHD3 |
| EHHADH |
| EIF1AY |
| ELAC1 |
| ELAVL4 |
| ELF5 |
| ELFN1 |
| ELOVL2 |
| ELOVL6 |
| EMILIN3 |
| EMP2 |
| EMR1 |
| EMR3 |
| ENDOU |
| ENHO |
| ENO3 |
| ENPEP |
| ENPP3 |
| ENPP6 |
| ENPP7 |
| ENTPD5 |
| ENTPD8 |
| EPB41L4B |
| EPHA1 |
| EPHA7 |
| EPHX1 |
| EPHX2 |
| EPO |
| EPOR |
| EPS8L2 |
| ERF |
| ERVFRD-1 |
| ESPN |
| ESPNL |
| ESR1 |
| ESRP1 |
| ETFDH |
| ETNK2 |
| EVA1A |
| EVPLL |
| EXOC3L4 |
| EXPH5 |
| EYA4 |
| F10 |

| |
|---|
| F11 |
| F12 |
| F13B |
| F2 |
| F7 |
| F8 |
| F9 |
| FAAH |
| FABP1 |
| FABP3 |
| FADS2 |
| FADS6 |
| FAH |
| FAM107A |
| FAM110C |
| FAM124A |
| FAM129C |
| FAM134B |
| FAM135B |
| FAM13A |
| FAM149A |
| FAM150B |
| FAM151A |
| FAM163B |
| FAM169A |
| FAM180A |
| FAM198A |
| FAM211B |
| FAM213A |
| FAM227B |
| FAM229B |
| FAM26F |
| FAM3B |
| FAM46C |
| FAM65C |
| FAM69B |
| FAM71D |
| FAM83B |
| FAM83E |
| FAM9B |
| FANCC |
| FAT3 |
| FAXDC2 |
| FBN3 |
| FBP1 |
| FBXO15 |
| FBXO2 |
| FBXO40 |
| FBXO6 |
| FBXO7 |
| FCAMR |
| FCAR |
| FCER1A |
| FCER2 |
| FCGR2B |
| FCGR3A |
| FCGR3B |
| FCGRT |

| |
|---|
| FCN1 |
| FCN2 |
| FCN3 |
| FCRL1 |
| FCRLB |
| FDPS |
| FERMT2 |
| FETUB |
| FEZ1 |
| FFAR2 |
| FGD4 |
| FGF10 |
| FGF21 |
| FGF23 |
| FGFBP2 |
| FGFR2 |
| FGFR3 |
| FGFR4 |
| FGL2 |
| FGR |
| FHL1 |
| FHOD3 |
| FITM1 |
| FLRT1 |
| FMO3 |
| FMO4 |
| FMO5 |
| FNDC5 |
| FNIP2 |
| FOLH1 |
| FOS |
| FOSB |
| FOXA2 |
| FOXA3 |
| FOXD3 |
| FOXN2 |
| FOXN3 |
| FOXO1 |
| FPR1 |
| FPR2 |
| FRAS1 |
| FRAT1 |
| FREM1 |
| FREM2 |
| FRMD1 |
| FRMD4B |
| FRMD7 |
| FRMPD4 |
| FRRS1 |
| FTCD |
| FTCDNL1 |
| FUOM |
| FUT3 |
| FUT6 |
| FUZ |
| FXN |
| FXYD1 |
| FXYD3 |

| |
|---|
| FXYD7 |
| FZD9 |
| G0S2 |
| GABRA2 |
| GABRG1 |
| GABRP |
| GADD45A |
| GADD45G |
| GAL3ST2 |
| GALK1 |
| GALM |
| GALNT14 |
| GALNT3 |
| GALT |
| GAMT |
| GAS2 |
| GATA4 |
| GATA5 |
| GATM |
| GBP1 |
| GBP7 |
| GCAT |
| GCDH |
| GCGR |
| GCHFR |
| GCK |
| GCKR |
| GCOM1 |
| GCSAML |
| GDF2 |
| GDPD4 |
| GFI1B |
| GFRA1 |
| GFRA2 |
| GFRA3 |
| GGACT |
| GGT6 |
| GHR |
| GHRL |
| GIPC2 |
| GJB2 |
| GJB3 |
| GJC3 |
| GLIPR1L2 |
| GLOD5 |
| GLP2R |
| GLUD1 |
| GLUL |
| GLYAT |
| GLYATL1 |
| GLYATL3 |
| GLYCTK |
| GMPR |
| GNA14 |
| GNAO1 |
| GNAT1 |
| GNE |
| GNG5P2 |

| |
|---|
| GNMT |
| GNPNAT1 |
| GNRH2 |
| GOLGA6A |
| GOLGA6B |
| GOLGA8M |
| GP1BA |
| GPAM |
| GPC6 |
| GPD1 |
| GPER1 |
| GPHN |
| GPLD1 |
| GPM6A |
| GPM6B |
| GPR123 |
| GPR125 |
| GPR126 |
| GPR128 |
| GPR142 |
| GPR143 |
| GPR158 |
| GPR162 |
| GPR182 |
| GPR75 |
| GPR82 |
| GPR83 |
| GPR88 |
| GPR98 |
| GPRIN2 |
| GRAMD1C |
| GRAP |
| GREB1 |
| GREB1L |
| GRHL1 |
| GRHL2 |
| GRHL3 |
| GRHPR |
| GRIK3 |
| GRM8 |
| GRTP1 |
| GSDMB |
| GSTA1 |
| GSTA2 |
| GSTM5 |
| GSTT2 |
| GSTT2B |
| GSTZ1 |
| GYLTL1B |
| GYS2 |
| GZMM |
| H1FX |
| HAAO |
| HABP2 |
| HAGH |
| HAMP |
| HAO1 |
| HAO2 |

| |
|---|
| HAP1 |
| HAS2 |
| HAUS4 |
| HBA1 |
| HBA2 |
| HBB |
| HBCBP |
| HBD |
| HBG2 |
| HCAR2 |
| HCAR3 |
| HDAC6 |
| HDC |
| HEMGN |
| HEPACAM |
| HERC5 |
| HEY2 |
| HFE2 |
| HGF |
| HGFAC |
| HHIP |
| HK3 |
| HLF |
| HLX |
| HMGCL |
| HMGCLL1 |
| HMGCR |
| HMGCS1 |
| HMGCS2 |
| HMOX1 |
| HNMT |
| HOGA1 |
| HOMER2 |
| HORMAD2 |
| HP |
| HPGDS |
| HPN |
| HPR |
| HPRT1 |
| HPX |
| HRASLS2 |
| HRG |
| HS3ST3A1 |
| HS3ST3B1 |
| HS3ST4 |
| HSBP1L1 |
| HSD11B1 |
| HSD17B11 |
| HSD17B13 |
| HSD17B14 |
| HSD17B6 |
| HSD17B8 |
| HSD3B1 |
| HSD3B2 |
| HSDL2 |
| HSPB9 |
| HYDIN |
| ICAM4 |

| |
|---|
| ID2 |
| IDNK |
| IDO2 |
| IFI27 |
| IFI44L |
| IFIT1 |
| IFIT1B |
| IFIT2 |
| IFIT3 |
| IFITM10 |
| IFNLR1 |
| IFT46 |
| IGF2 |
| IGFALS |
| IGFBP1 |
| IGFBP2 |
| IGFBP3 |
| IGJ |
| IGLON5 |
| IGSF10 |
| IGSF23 |
| IGSF9 |
| IHH |
| IL10 |
| IL11RA |
| IL13RA2 |
| IL17RB |
| IL17RC |
| IL17RE |
| IL1B |
| IL1RAP |
| IL1RN |
| IL20RA |
| IL22RA1 |
| IL23A |
| IL27 |
| IL33 |
| IMPA2 |
| INHBC |
| INMT |
| INS-IGF2 |
| INSIG2 |
| IQGAP2 |
| IQSEC3 |
| IRF6 |
| IRF7 |
| ISPD |
| ITCH |
| ITGA2B |
| ITGA9 |
| ITGAD |
| ITIH1 |
| ITIH2 |
| ITIH3 |
| ITLN1 |
| ITPR2 |
| IYD |
| JAKMIP2 |

| |
|---|
| JDP2 |
| JUN |
| JUNB |
| JUND |
| KANK4 |
| KAZN |
| KBTBD11 |
| KCNAB1 |
| KCND3 |
| KCNE1 |
| KCNH7 |
| KCNH8 |
| KCNJ10 |
| KCNJ13 |
| KCNJ15 |
| KCNJ3 |
| KCNK1 |
| KCNK17 |
| KCNK3 |
| KCNK5 |
| KCNMB2 |
| KCNN2 |
| KCTD16 |
| KDM8 |
| KHK |
| KIAA1161 |
| KIF12 |
| KIF17 |
| KIF19 |
| KIF1A |
| KIF25 |
| KIF26A |
| KIF6 |
| KIRREL3 |
| KLB |
| KLC4 |
| KLF10 |
| KLF11 |
| KLF12 |
| KLF4 |
| KLHL13 |
| KLHL33 |
| KLHL4 |
| KLKB1 |
| KLLN |
| KLRF1 |
| KNG1 |
| KRBOX1 |
| KRT1 |
| KRT19 |
| KRT73 |
| KRTAP5-9 |
| KRTCAP3 |
| L1CAM |
| LAG3 |
| LARGE |
| LBX2 |
| LCAT |

| |
|---|
| LDLR |
| LDLRAD4 |
| LEAP2 |
| LEPREL1 |
| LGALS12 |
| LGALS4 |
| LGI1 |
| LGI4 |
| LGSN |
| LHX2 |
| LIFR |
| LILRA1 |
| LILRA2 |
| LILRA5 |
| LILRA6 |
| LILRB1 |
| LILRB2 |
| LILRB3 |
| LILRB5 |
| LIMD2 |
| LIME1 |
| LINC00923 |
| LINGO4 |
| LIPA |
| LIPC |
| LIPG |
| LIPJ |
| LIPN |
| LNP1 |
| LONRF3 |
| LPA |
| LPIN2 |
| LRAT |
| LRFN5 |
| LRIG3 |
| LRP1B |
| LRP3 |
| LRRC16B |
| LRRC19 |
| LRRC25 |
| LRRC3 |
| LRRC31 |
| LRRC3DN |
| LRRC4 |
| LRRC55 |
| LRRFIP2 |
| LRRIQ1 |
| LRRK2 |
| LRRN1 |
| LRRN3 |
| LRRN4 |
| LRRTM1 |
| LRRTM2 |
| LRRTM4 |
| LSR |
| LSS |
| LST1 |
| LST3 |

| |
|---|
| LTK |
| LY6E |
| LYNX1 |
| LYVE1 |
| LZTFL1 |
| MAATS1 |
| MACROD1 |
| MAFF |
| MAG |
| MAGEH1 |
| MAK |
| MAN1C1 |
| MAOA |
| MAOB |
| MAP1LC3A |
| MAP3K13 |
| MAPK4 |
| MARCO |
| MARVELD3 |
| MASP1 |
| MASP2 |
| MAT1A |
| MATN2 |
| MBL2 |
| MBNL3 |
| MBOAT1 |
| MCHR1 |
| MDGA2 |
| MEFV |
| MEGF10 |
| MEI4 |
| MEIOB |
| MEP1B |
| MEST |
| METTL20 |
| METTL7A |
| MFAP3L |
| MFI2 |
| MFSD2A |
| MFSD4 |
| MGAM |
| MGMT |
| MGST2 |
| MLANA |
| MLF1 |
| MLIP |
| MLK4 |
| MMAB |
| MME |
| MMP19 |
| MMP25 |
| MMP7 |
| MMP8 |
| MMRN1 |
| MOGAT2 |
| MOGAT3 |
| MORC1 |
| MORC3 |

| |
|---|
| MPDZ |
| MPPED1 |
| MPST |
| MRC1L1 |
| MREG |
| MRGPRF |
| MRO |
| MROH2A |
| MROH2B |
| MROH7 |
| MROH8 |
| MRPL23 |
| MRPS6 |
| MS4A2 |
| MS4A6A |
| MSMO1 |
| MST1 |
| MT1G |
| MT1H |
| MT1M |
| MTHFD1 |
| MTHFS |
| MTTP |
| MTUS2 |
| MUSK |
| MUT |
| MVK |
| MYCL |
| MYCT1 |
| MYH7B |
| MYL3 |
| MYO15A |
| MYO16 |
| MYO1B |
| MYO1F |
| MYO3A |
| MYRIP |
| MYT1L |
| N4BP2L1 |
| NAAA |
| NAALAD2 |
| NAB2 |
| NAGS |
| NANOS1 |
| NAT1 |
| NAT2 |
| NAT8 |
| NCAM1 |
| NCF1 |
| NCKAP5 |
| NCMAP |
| NCR1 |
| NDNF |
| NDRG2 |
| NDST3 |
| NECAB2 |
| NEIL1 |
| NEU4 |

| |
|---|
| NFAM1 |
| NFATC2 |
| NFE2 |
| NFIA |
| NFKBIA |
| NFKBIZ |
| NGF |
| NGFR |
| NGFRAP1 |
| NHSL1 |
| NIPAL1 |
| NIPSNAP1 |
| NLRC4 |
| NLRP11 |
| NLRP12 |
| NLRP14 |
| NLRP3 |
| NLRP6 |
| NMRK1 |
| NOL4 |
| NOS1 |
| NOS1AP |
| NOTUM |
| NOXO1 |
| NPAS3 |
| NPC1L1 |
| NPR1 |
| NPR2 |
| NR0B2 |
| NR1H3 |
| NR1I2 |
| NR1I3 |
| NR2F6 |
| NR5A2 |
| NRAP |
| NREP |
| NRG1 |
| NRG3 |
| NRG4 |
| NRN1 |
| NRXN1 |
| NSUN6 |
| NSUN7 |
| NT5DC3 |
| NT5E |
| NTF3 |
| NTHL1 |
| NTN1 |
| NTN3 |
| NTN4 |
| NTRK1 |
| NUDT10 |
| NUDT4 |
| NUDT6 |
| NUGGC |
| NUP62CL |
| NXF3 |
| NYAP1 |

| |
|---|
| OAF |
| OAS1 |
| OCLN |
| ODF3L1 |
| OGDHL |
| OIT3 |
| OLFM1 |
| OLFM4 |
| OLFML3 |
| OPRK1 |
| OR10J5 |
| OR13C3 |
| OR13C4 |
| OR13C5 |
| OR13C9 |
| OR2W3 |
| ORMDL3 |
| OSBPL6 |
| OSGIN1 |
| OTC |
| OTOA |
| OXER1 |
| OXT |
| P2RX3 |
| P2RX4 |
| P2RX6 |
| P2RY12 |
| P2RY13 |
| P2RY2 |
| P4HA1 |
| PACRG |
| PACSIN1 |
| PACSIN3 |
| PADI4 |
| PAGE5 |
| PAK7 |
| PALM3 |
| PAMR1 |
| PANK1 |
| PANX2 |
| PAQR7 |
| PAQR9 |
| PATZ1 |
| PBLD |
| PC |
| PCDH15 |
| PCDH20 |
| PCDH9 |
| PCDHAC1 |
| PCDHAC2 |
| PCK2 |
| PCLO |
| PCOLCE2 |
| PCP2 |
| PCP4L1 |
| PCSK2 |
| PCSK9 |
| PCYT2 |

| |
|---|
| PDCD1LG2 |
| PDE11A |
| PDE2A |
| PDE6G |
| PDE8B |
| PDIA5 |
| PDK2 |
| PDLIM2 |
| PDXP |
| PDZRN4 |
| PEBP1 |
| PEBP4 |
| PECR |
| PEG10 |
| PEG3 |
| PEMT |
| PER2 |
| PER3 |
| PEX11G |
| PFKFB1 |
| PGAM2 |
| PGAP3 |
| PGLYRP2 |
| PGM1 |
| PGM2 |
| PGM5 |
| PGRMC1 |
| PHACTR3 |
| PHGDH |
| PHLDA1 |
| PHLPP1 |
| PHOSPHO1 |
| PHYH |
| PHYHD1 |
| PHYHIPL |
| PI16 |
| PID1 |
| PIGR |
| PIK3AP1 |
| PIK3C2G |
| PIK3R1 |
| PIM1 |
| PIPOX |
| PITPNM3 |
| PKD2L1 |
| PKHD1L1 |
| PKLR |
| PKP2 |
| PLA1A |
| PLA2G12B |
| PLA2G16 |
| PLAC8 |
| PLCL2 |
| PLCXD2 |
| PLD5 |
| PLEK2 |
| PLEKHA4 |
| PLEKHA6 |

| |
|---|
| PLEKHB1 |
| PLEKHF1 |
| PLEKHG6 |
| PLEKHG7 |
| PLG |
| PLGLB1 |
| PLGLB2 |
| PLIN1 |
| PLIN2 |
| PLK3 |
| PLP1 |
| PLSCR4 |
| PM20D1 |
| PMEL |
| PMM1 |
| PMP2 |
| PNMA3 |
| PNMA6C |
| PNMAL2 |
| PNPLA3 |
| PNPLA7 |
| POFUT1 |
| POLR3GL |
| POMC |
| PON3 |
| POU6F2 |
| PPAP2B |
| PPARA |
| PPBP |
| PPFIBP2 |
| PPP1R15A |
| PPP1R1A |
| PPP1R3B |
| PPP1R3C |
| PPP4R4 |
| PPP6R2 |
| PQLC1 |
| PRAM1 |
| PRAP1 |
| PRELID2 |
| PRG4 |
| PRH2 |
| PRIMA1 |
| PRKAR2B |
| PRKCB |
| PRLR |
| PROC |
| PRODH2 |
| PROK2 |
| PROM1 |
| PROSER2 |
| PROZ |
| PRPSAP1 |
| PRR18 |
| PRR22 |
| PRR5 |
| PRRG4 |
| PRSS12 |

| |
|---|
| PRSS22 |
| PRSS36 |
| PRSS42 |
| PRSS45 |
| PRSS50 |
| PRSS53 |
| PRSS8 |
| PSAT1 |
| PTCHD3 |
| PTCRA |
| PTGDR2 |
| PTGR1 |
| PTGS2 |
| PTH1R |
| PTK6 |
| PTMS |
| PTN |
| PTPRB |
| PTPRD |
| PTPRN2 |
| PTPRS |
| PTPRT |
| PVALB |
| PVRL3 |
| PXMP2 |
| PYGL |
| PYROXD2 |
| PZP |
| QPRT |
| QRICH2 |
| QSOX1 |
| RAB17 |
| RAB25 |
| RAB26 |
| RAB27B |
| RAB39A |
| RAB3IL1 |
| RAD51AP2 |
| RAD54L2 |
| RAD9B |
| RAG1 |
| RALGPS2 |
| RALYL |
| RANBP3L |
| RARRES2 |
| RARRES3 |
| RASGEF1B |
| RASGRP2 |
| RASGRP4 |
| RASL10A |
| RASL10B |
| RASL11A |
| RASSF5 |
| RBP5 |
| RCAN1 |
| RCL1 |
| RD3L |
| RDH16 |

| |
|---|
| RENBP |
| RET |
| RFNG |
| RFPL1 |
| RGL1 |
| RGN |
| RGPD2 |
| RGS18 |
| RGS3 |
| RGSL1 |
| RHBG |
| RHCE |
| RHOB |
| RIC3 |
| RIPK4 |
| RIPPLY1 |
| RIPPLY3 |
| RMDN2 |
| RMND5A |
| RND2 |
| RNF125 |
| RNF144B |
| RNF152 |
| RNF165 |
| ROBO2 |
| ROPN1L |
| RORC |
| ROS1 |
| RP11-10A14.4 |
| RP11-146D12.2 |
| RP11-181C3.1 |
| RP11-242G20.1 |
| RP11-321F6.1 |
| RP11-422N16.3 |
| RP11-595B24.2 |
| RP11-650K20.3 |
| RP11-676J12.7 |
| RP11-766F14.2 |
| RP11-817J15.3 |
| RP11-867G23.8 |
| RP11-986E7.7 |
| RP11-998D10.1 |
| RPGRIP1 |
| RPS29 |
| RPS4Y1 |
| RPS6KA6 |
| RSAD2 |
| RSPH10B |
| RSPH4A |
| RSPO2 |
| RTN4RL1 |
| RTN4RL2 |
| RTP3 |
| RTP4 |
| RTTN |
| RUNDC3B |
| RXFP1 |
| RXRG |

| |
|---|
| S100A1 |
| S100A12 |
| S100A14 |
| S100A8 |
| S100A9 |
| S1PR5 |
| SAA4 |
| SALL4 |
| SAMD4A |
| SAMD5 |
| SARDH |
| SAT2 |
| SATB1 |
| SC5D |
| SCG5 |
| SCGB3A1 |
| SCIMP |
| SCN11A |
| SCN2A |
| SCN3A |
| SCN7A |
| SCN9A |
| SCNN1B |
| SCNN1D |
| SCP2 |
| SCRN2 |
| SDC3 |
| SDK2 |
| SDPR |
| SEC14L2 |
| SEC14L3 |
| SEC14L4 |
| SELE |
| SELENBP1 |
| SELO |
| SEMA3D |
| SEMA3E |
| SEMA4A |
| SEMA4G |
| SEMA6A |
| SEMA6C |
| SEMA6D |
| SEPP1 |
| SERP2 |
| SERPINA10 |
| SERPINA11 |
| SERPINA4 |
| SERPINA5 |
| SERPINA6 |
| SERPINA7 |
| SERPINC1 |
| SERPIND1 |
| SERPINE1 |
| SERPINF1 |
| SERPINF2 |
| SEZ6L |
| SFRP1 |
| SFRP5 |

| |
|---|
| SFTPD |
| SFXN5 |
| SGCE |
| SGCZ |
| SH2D1B |
| SH2D4A |
| SH2D6 |
| SH3BP2 |
| SH3BP5 |
| SH3GL2 |
| SH3RF2 |
| SHB |
| SHBG |
| SHD |
| SHF |
| SHH |
| SHMT1 |
| SHROOM2 |
| SIGIRR |
| SIGLEC1 |
| SIGLEC11 |
| SIGLEC14 |
| SIGLEC15 |
| SIGLEC7 |
| SIGLEC9 |
| SIM1 |
| SIRPB1 |
| SIRT5 |
| SIVA1 |
| SKAP1 |
| SKIDA1 |
| SKOR1 |
| SLAIN1 |
| SLC10A1 |
| SLC13A5 |
| SLC15A1 |
| SLC16A1 |
| SLC16A10 |
| SLC16A12 |
| SLC16A2 |
| SLC16A4 |
| SLC16A9 |
| SLC17A2 |
| SLC17A8 |
| SLC18A2 |
| SLC19A1 |
| SLC19A2 |
| SLC19A3 |
| SLC22A1 |
| SLC22A10 |
| SLC22A24 |
| SLC22A25 |
| SLC22A7 |
| SLC22A9 |
| SLC23A1 |
| SLC24A2 |
| SLC25A1 |
| SLC25A13 |

| |
|---|
| SLC25A18 |
| SLC25A20 |
| SLC25A21 |
| SLC25A21-AS1 |
| SLC25A27 |
| SLC25A34 |
| SLC25A47 |
| SLC26A1 |
| SLC26A5 |
| SLC27A2 |
| SLC27A3 |
| SLC27A5 |
| SLC28A1 |
| SLC28A2 |
| SLC2A12 |
| SLC2A2 |
| SLC2A4RG |
| SLC2A9 |
| SLC30A1 |
| SLC30A10 |
| SLC31A2 |
| SLC34A1 |
| SLC35D1 |
| SLC37A4 |
| SLC38A11 |
| SLC38A4 |
| SLC39A4 |
| SLC39A5 |
| SLC3A1 |
| SLC43A3 |
| SLC45A3 |
| SLC47A1 |
| SLC4A1 |
| SLC51A |
| SLC52A1 |
| SLC5A1 |
| SLC5A7 |
| SLC5A9 |
| SLC6A1 |
| SLC6A12 |
| SLC6A13 |
| SLC6A19 |
| SLC6A20 |
| SLC6A4 |
| SLC7A8 |
| SLC7A9 |
| SLC8A1 |
| SLC9A3R2 |
| SLC9B2 |
| SLCO1A2 |
| SLCO1B3 |
| SLCO1B7 |
| SLCO2B1 |
| SLCO4C1 |
| SLITRK2 |
| SLITRK3 |
| SLITRK6 |
| SMAD6 |

| |
|---|
| SMCO3 |
| SMIM1 |
| SMIM14 |
| SMIM19 |
| SMIM9 |
| SMLR1 |
| SMO |
| SMPD3 |
| SNCA |
| SNTB1 |
| SNTG1 |
| SOAT2 |
| SOBP |
| SOCS2 |
| SOCS6 |
| SOD1 |
| SORCS1 |
| SORD |
| SORL1 |
| SOX10 |
| SOX5 |
| SPDYC |
| SPECC1L-ADORA2A |
| SPI1 |
| SPIB |
| SPIC |
| SPINT2 |
| SPOCK3 |
| SPP2 |
| SPSB3 |
| SPSB4 |
| SPTBN2 |
| SQLE |
| SRCIN1 |
| SRD5A1 |
| SRPX |
| SSTR1 |
| SSTR2 |
| ST14 |
| ST3GAL1 |
| ST3GAL6 |
| ST6GAL1 |
| ST6GALNAC2 |
| ST6GALNAC3 |
| ST8SIA3 |
| STAB1 |
| STAB2 |
| STAG3 |
| STARD10 |
| STARD4 |
| STEAP3 |
| STEAP4 |
| STMND1 |
| STPG2 |
| SUCNR1 |
| SULT1A1 |
| SULT1A2 |
| SULT1E1 |

| |
|---|
| SULT2A1 |
| SUN2 |
| SUSD4 |
| SYBU |
| SYCE1 |
| SYDE2 |
| SYNE4 |
| SYNGR1 |
| SYT1 |
| SYT10 |
| SYT12 |
| SYT15 |
| SYT17 |
| SYT7 |
| SYTL4 |
| TACSTD2 |
| TADA1 |
| TAS1R3 |
| TBL1Y |
| TBX20 |
| TBXA2R |
| TBXAS1 |
| TCEA3 |
| TCEAL2 |
| TCF21 |
| TCHH |
| TCL1A |
| TCP10L |
| TCP10L2 |
| TCTEX1D1 |
| TCTEX1D4 |
| TDRD10 |
| TDRD6 |
| TEF |
| TEK |
| TEKT2 |
| TEKT5 |
| TENM1 |
| TENM2 |
| TESK2 |
| TEX30 |
| TF |
| TFPI2 |
| TFR2 |
| TGFBR3 |
| THEMIS2 |
| THNSL1 |
| THOP1 |
| THRSP |
| TIAM1 |
| TIMD4 |
| TINAGL1 |
| TIPARP |
| TJP2 |
| TKTL1 |
| TLR4 |
| TM6SF2 |
| TMCO6 |

| |
|---|
| TMEFF2 |
| TMEM105 |
| TMEM121 |
| TMEM125 |
| TMEM132C |
| TMEM132D |
| TMEM139 |
| TMEM150C |
| TMEM170B |
| TMEM176B |
| TMEM200B |
| TMEM200C |
| TMEM220 |
| TMEM232 |
| TMEM25 |
| TMEM252 |
| TMEM26 |
| TMEM27 |
| TMEM30B |
| TMEM37 |
| TMEM45B |
| TMEM47 |
| TMEM52 |
| TMEM56 |
| TMEM63C |
| TMEM71 |
| TMEM82 |
| TMEM86B |
| TMEM97 |
| TMIE |
| TMPO |
| TMPRSS2 |
| TMPRSS4 |
| TMPRSS6 |
| TMPRSS9 |
| TMSB4Y |
| TNF |
| TNFAIP8L1 |
| TNFRSF11B |
| TNFSF10 |
| TNFSF11 |
| TNN |
| TNNC1 |
| TNR |
| TOM1L1 |
| TOX2 |
| TP53I13 |
| TP53INP1 |
| TPH2 |
| TPPP2 |
| TPRG1 |
| TPSAB1 |
| TPST2 |
| TRABD2B |
| TRAPPC3L |
| TRDN |
| TREH |
| TREM1 |

| |
|---|
| TREML2 |
| TRHDE |
| TRIB1 |
| TRIM58 |
| TRIM63 |
| TRPC5 |
| TRPM6 |
| TRPM8 |
| TRPV4 |
| TRPV6 |
| TSHR |
| TSLP |
| TSPAN11 |
| TSPAN12 |
| TSPAN7 |
| TSPAN9 |
| TST |
| TSTD1 |
| TTBK1 |
| TTC36 |
| TTC38 |
| TTC40 |
| TTC7B |
| TTPA |
| TTR |
| TUBB1 |
| TUBE1 |
| TUSC1 |
| TXNDC16 |
| TXNIP |
| TXNRD2 |
| UAP1 |
| UGP2 |
| UGT1A1 |
| UGT2B10 |
| UGT2B15 |
| UGT2B17 |
| UGT2B7 |
| UGT3A1 |
| UNC13D |
| UNC79 |
| UNC93A |
| UPB1 |
| UROC1 |
| USH2A |
| USP18 |
| USP44 |
| USP51 |
| UTY |
| VCAM1 |
| VEPH1 |
| VIL1 |
| VIPR1 |
| VIPR2 |
| VMO1 |
| VNN1 |
| VNN2 |
| VNN3 |

| |
|---|
| VPS37B |
| VPS37D |
| VSIG4 |
| VSNL1 |
| VTCN1 |
| VWA3B |
| VWCE |
| VWDE |
| WBSCR27 |
| WDR17 |
| WDR72 |
| WNK2 |
| WNK3 |
| WNT11 |
| WNT5A |
| WNT5B |
| WNT7A |
| XAGE3 |
| XDH |
| XKR4 |
| XPNPEP2 |
| XRCC6BP1 |
| XYLB |
| YBX2 |
| YPEL1 |
| YPEL2 |
| ZBTB18 |
| ZC3H12C |
| ZCCHC6 |
| ZCWPW1 |
| ZDHHC19 |
| ZFP1 |
| ZFY |
| ZG16 |
| ZIC1 |
| ZMYND12 |
| ZNF175 |
| ZNF268 |
| ZNF311 |
| ZNF334 |
| ZNF354C |
| ZNF358 |
| ZNF367 |
| ZNF385B |
| ZNF385C |
| ZNF470 |
| ZNF471 |
| ZNF502 |
| ZNF511 |
| ZNF536 |
| ZNF572 |
| ZNF577 |
| ZNF648 |
| ZNF662 |
| ZNF676 |
| ZNF682 |
| ZNF812 |
| ZNF879 |

| |
|---|
| ZPBP |
| ZPLD1 |
| ZSCAN18 |
| ZYG11A |

**Supplementary Table 5A.** Genes modulated by BLU0588 versus DMSO

| BLU588 Neg | shRNA Neg | Overlap | Hallmark |
|---|---|---|---|
| ROPN1L | PPARGC1A | ADH4 | XENOBIOTIC_METABOLISM |
| SCARA5 | FKBP11 | AKR1B15 | TNFA_SIGNALING_VIA_NFKB |
| BPIFA1 | PDE4B | AKR1C1 | HYPOXIA |
| CYP24A1 | ENTPD1 | ALDH1L1 | UV_RESPONSE_UP |
| C11orf86 | CPS1 | APOB | COMPLEMENT |
| SLC2A9 | DDIT4 | ATP1B1 | HEME_METABOLISM |
| MAPK4 | SLC16A11 | AVPI1 | |
| RIMKLB | RHOBTB1 | BASP1 | |
| IGFBP1 | C8G | BRD4 | |
| CD55 | COL5A3 | C11orf86 | |
| TSNAX-DISC1 | NEIL3 | CA2 | |
| SPATA31A6 | AKR1C3 | CABYR | |
| CPS1 | CGREF1 | CD55 | |
| KRT86 | TMOD1 | CDHR5 | |
| PTGS2 | GLA | CPB2 | |
| STC1 | HPD | CPLX2 | |
| CYSLTR1 | UBXN10 | CPS1 | |
| INSL4 | SLC5A6 | CU639417.2 | |
| PTPN5 | TLL2 | CYP24A1 | |
| CHST8 | KYNU | CYP2C18 | |
| SCNN1B | ATP1B1 | DDIT4 | |
| SPATA31A1 | E2F8 | DNAJC12 | |
| ASPG | TM4SF5 | DUSP1 | |
| SPATA31A7 | CTH | EPAS1 | |
| DUSP1 | PRKACA | EVA1C | |
| TNFSF11 | KCNE4 | FERMT1 | |
| ITM2A | GK | FMO5 | |
| GALNTL6 | EVA1C | FSTL4 | |
| LRRC32 | CA9 | G6PC | |
| ANGPTL3 | SLC7A1 | GDF7 | |
| KCNK3 | CORIN | GGH | |
| MPP1 | GPRIN3 | GK | |
| ANXA10 | SLC22A11 | GPRIN3 | |
| NTS | PBK | GPT | |
| RBP4 | PDE3A | HEPACAM | |
| GALNT8 | UTS2B | HMOX1 | |
| PPP1R3G | SDCBP2 | HPD | |
| ITIH4 | HKDC1 | HYAL1 | |
| CU639417.2 | CABYR | ID4 | |
| ACKR3 | NEB | KCNE3 | |

| | | |
|---|---|---|
| OXGR1 | KRT86 | KCNU1 |
| BASP1 | EPS8L3 | KHK |
| NR4A1 | TESC | KRT86 |
| SIK1 | FSTL4 | KYNU |
| SMIM9 | AKR1B15 | LYPD3 |
| KRT83 | S100P | MPP1 |
| TFCP2L1 | MUC5B | MUC13 |
| G6PC | KCNU1 | NAT6 |
| SLC5A7 | CALCA | NR4A1 |
| GK | CPLX2 | PAH |
| PTPRN | MUC13 | PBK |
| BMP2 | CPB2 | PC |
| MFAP5 | CLDN2 | PCK1 |
| PDK4 | PTGS2 | PDE3A |
| C11orf96 | SMIM9 | PDE4B |
| C3orf80 | SCARA5 | PDE7B |
| CD247 | CYP24A1 | PDK4 |
| DCLK1 | UGT2B10 | PER2 |
| VGF | ROPN1L | PEX11A |
| ADAMTS12 | APOB | PGC |
| ENTPD8 | C11orf86 | PLA2G4A |
| SCUBE1 | PDK4 | PPARGC1A |
| ADH4 | SLC39A5 | PREB |
| PDE10A | ADH4 | PTGS2 |
| PSG5 | MPP1 | RBFOX3 |
| APOA5 | HEPACAM | RHOBTB1 |
| FAM155A | CA2 | RND1 |
| HOPX | PCK1 | ROPN1L |
| PLA2G4A | C8A | S100P |
| ALOX15B | ADH6 | SCARA5 |
| PLIN1 | ASGR1 | SDCBP2 |
| NR4A3 | ABCG2 | SIK1 |
| HEPACAM | C3 | SLC13A3 |
| CLU | ALDOB | SLC16A11 |
| MAMLD1 | KHK | SLC22A11 |
| CA2 | PRODH2 | SLC5A6 |
| CPB2 | LYPD3 | SMIM9 |
| BMP6 | S100A14 | SRXN1 |
| PCK1 | PLA2G4A | TCF7L1 |
| FOXC1 | PKLR | TESC |
| KRT17 | ALDH1L1 | TGFBR3L |
| EVA1C | PDE7B | TNFSF11 |
| NEFL | SIK1 | TTPA |

| | | |
|---|---|---|
| KRT81 | CU639417.2 | UGT1A3 |
| RHOBTB1 | MTTP | UGT1A4 |
| S1PR1 | TNFSF11 | UGT2A3 |
| NUGGC | G6PC | UGT2B17 |
| GBP4 | SLC6A4 | ZFP36 |
| RBFOX3 | SULT1B1 | |
| TSNAX | ANPEP | |
| INHA | DEGS2 | |
| PDE3A | SLC23A1 | |
| IGFBP3 | PAGE4 | |
| TESC | UGT2B17 | |
| NOTCH3 | ANKS4B | |
| CPLX1 | GPT | |
| SLC16A6 | PGC | |
| KRT7 | MAT1A | |
| FA2H | HPN | |
| CRYAB | PKDCC | |
| KRT16 | ARHGEF16 | |
| HRH2 | TCF7L1 | |
| PLOD2 | CDHR5 | |
| SLC22A11 | RBFOX3 | |
| GOLGA6D | HMOX1 | |
| PABPC1L2A | UGT2A3 | |
| FXYD4 | F2 | |
| FST | AKR1B10 | |
| FXYD2 | PREB | |
| RASD1 | SLC16A13 | |
| SLC2A1 | UGT1A4 | |
| NPR3 | SSTR1 | |
| PEX11A | FMO5 | |
| SLC5A6 | XPNPEP2 | |
| C10orf90 | PAH | |
| KCNJ16 | NOSTRIN | |
| P4HA3 | SLCO2B1 | |
| HEY1 | GOLGA6A | |
| PGC | AL132639.3 | |
| PPARGC1A | DDC | |
| UGT1A4 | PCCA | |
| GRIP2 | HRCT1 | |
| APOA4 | MYO1A | |
| ID4 | AGMAT | |
| UGT2B15 | ANG | |
| LYPD3 | APOC4-APOC2 | |

| | |
|---|---|
| GPT | CYP4F3 |
| GDF7 | HYAL1 |
| PSG6 | AL163636.2 |
| PDE7B | SRGN |
| ODC1 | SLC13A3 |
| PDE4B | CD55 |
| KCNJ15 | GAS2 |
| PCP4 | RND1 |
| NR4A2 | FGL1 |
| RGCC | CDHR2 |
| IRS2 | CBR1 |
| ATP1B1 | HNF4A |
| VSIG1 | CLDN7 |
| LGALS3 | FERMT1 |
| ALDH1L1 | DNAJC12 |
| LRRK1 | PEX11A |
| GABRQ | CD68 |
| BLOC1S5-TXND | GSTA2 |
| EFCAB12 | SLC19A3 |
| CABYR | CMBL |
| HHLA3 | F7 |
| HYAL1 | GGH |
| PSTPIP2 | DACT2 |
| PTP4A1 | IQCH |
| PER2 | NR4A1 |
| CRABP1 | CREB3L3 |
| EPAS1 | THRB |
| F2RL2 | ENTPD2 |
| PBK | SLC43A1 |
| SPATA17 | CYP4F11 |
| UGT1A1 | SERPINA10 |
| SLC16A11 | PC |
| FGA | RNASE4 |
| CPA4 | TTPA |
| GABRP | WIF1 |
| SYT12 | TUBAL3 |
| ANGPTL4 | SOWAHA |
| AKR1B15 | LENG9 |
| SLC13A3 | PTGR1 |
| BCAS1 | PDE8B |
| KHK | NGEF |
| AVPI1 | ESRP2 |
| FSTL4 | CCL15 |

| | |
|---|---|
| PTPRB | PER2 |
| SGK1 | GCLM |
| CXCL5 | PIPOX |
| DNAJC12 | DUSP1 |
| GPRIN3 | MLKL |
| DIO2 | EDNRA |
| CCDC68 | TTC22 |
| KCNU1 | CYP2C18 |
| S100P | AVPI1 |
| KYNU | CIDEB |
| GJA1 | ID4 |
| CSGALNACT1 | BASP1 |
| B3GALT1 | EIF4EBP3 |
| RASSF9 | APOH |
| UGT1A3 | TFR2 |
| HYAL3 | FRRS1 |
| GGH | TGFBR3L |
| TCF7L1 | CES1 |
| ZBED2 | AKR1C4 |
| AKAP12 | MGST1 |
| FABP5 | GRTP1 |
| ADSSL1 | GJB1 |
| CPLX2 | SRXN1 |
| WNT11 | MST1 |
| PTPRM | APOC1 |
| RAB27B | MYO7B |
| SLC11A1 | UGDH |
| IRX3 | MTRR |
| RHOF | AMBP |
| CPEB4 | LDHD |
| GADD45B | NR1I2 |
| SDCBP2 | GLYCTK |
| MUC13 | AKR1C1 |
| PDZRN3 | ESPN |
| SLC6A12 | CHST13 |
| CH25H | METTL7A |
| HPD | A1CF |
| RPL17-C18orf32 | KCNE3 |
| MYOM2 | KLF15 |
| C4B | AMT |
| CST1 | SGK2 |
| AKR1C2 | SLC1A2 |
| CST7 | HSD11B2 |

| | |
|---|---|
| HMOX1 | UBA7 |
| SPP1 | EPAS1 |
| CST2 | GPAM |
| CXCL12 | TRIM15 |
| FMO5 | NAT6 |
| RASAL1 | CDH16 |
| BHLHE40 | ELMO3 |
| DUXA | BRD4 |
| SEMA3F | KCNQ1 |
| CMIP | LRRC8C |
| PREB | NDRG2 |
| PAH | GDF7 |
| TFPI | C2orf72 |
| SLC4A4 | HAAO |
| UGT2A3 | ZFP36 |
| ABCA1 | AGFG2 |
| ONECUT2 | NR1H3 |
| FMNL1 | EPB41L4B |
| LATS2 | LRIG3 |
| PLPP1 | AMN |
| PGM2L1 | BHMT2 |
| PC | ARHGAP4 |
| URB1 | ZNF331 |
| FSTL3 | COQ8A |
| DAB2 | NR0B1 |
| PLAC8 | SAT1 |
| IL10RA | SHROOM1 |
| IRX6 | UGT1A3 |
| PRKD3 | PECR |
| GCH1 | CGN |
| FERMT1 | |
| TTPA | |
| ZFP36 | |
| OSBP2 | |
| DNM3 | |
| ANXA1 | |
| HHAT | |
| MYO1E | |
| TBX3 | |
| SPANXD | |
| MN1 | |
| ENO3 | |
| CBR3 | |

| |
|---|
| MID1 |
| DGKD |
| SPANXA1 |
| CREG1 |
| CASP9 |
| HHIPL2 |
| RND1 |
| USP28 |
| CHCHD10 |
| NAT6 |
| APOB |
| RAB3B |
| TBX4 |
| KLB |
| ITGB8 |
| ADAM12 |
| DNAJC15 |
| FOXL1 |
| TLCD2 |
| ATF3 |
| SQSTM1 |
| LRP8 |
| SRXN1 |
| DGKI |
| CDHR5 |
| CKB |
| MSLN |
| TGFBR3L |
| RAB10 |
| SMAD2 |
| PIM3 |
| B4GALT1 |
| ISG20 |
| PLA2G4F |
| HES4 |
| PAG1 |
| PITX1 |
| BRD4 |
| TP53INP2 |
| BAG1 |
| KCNMB4 |
| PELI2 |
| SIAH2 |

| |
|---|
| PBX1 |
| HGD |
| ARG2 |
| UGT2B17 |
| C1QTNF6 |
| CYP2C18 |
| HSPB8 |
| LARGE1 |
| MAFK |
| CA12 |
| FAS |
| AKR1C1 |
| MESP1 |
| CNKSR3 |
| SCN1B |
| FAM167A |
| PRICKLE2 |
| DDIT4 |
| DDIT3 |
| PPP2R1B |
| GCLC |
| OAT |
| ITPRIP |
| NTRK2 |
| KCNE3 |

**Supplementary Table 5B.** Genes modulated by BLU0588 versus DMSO

| BLU588 Pos | shRNA Pos | Overlap | Hallmark |
|---|---|---|---|
| CMKLR1 | IGLON5 | ADAMTSL2 | COAGULATION |
| MUCL1 | ETNK2 | AHRR | EPITHELIAL_MESENCHYMAL_TRANSITION |
| CA4 | GJB3 | AMIGO2 | HEDGEHOG_SIGNALING |
| RSPO4 | CASC10 | AMOT | APOPTOSIS |
| OLFML3 | IGSF9 | ANGPTL2 | COMPLEMENT |
| MMP3 | IFIT1 | ANK1 | |
| CCM2L | KRT19 | ASAP3 | |
| CST6 | ECM1 | B3GNT7 | |
| NXPH3 | TMEM63C | BCAM | |
| DBH | SKIDA1 | BMF | |
| WNT16 | WNT11 | BMP7 | |
| VWA2 | MT1G | C15orf52 | |
| MMP12 | ADAMTSL2 | CACNA1S | |
| PNMT | TRIB1 | CACNB3 | |
| ANKRD1 | HMGCS2 | CAPN5 | |
| ADAMTSL2 | ANXA3 | CD8B | |
| LIX1 | NTN4 | CERCAM | |
| CARD11 | COL18A1 | CHRNA3 | |
| SERPINI1 | NGFR | CHST3 | |
| WFDC2 | CDH2 | CLIC5 | |
| LOXL4 | CYP2S1 | CLIP3 | |
| LYPD6B | JUN | CMTM3 | |
| CHRNA3 | RHOB | CPNE2 | |
| CTSE | CAPN5 | CRIP2 | |
| BMP7 | ARVCF | CYP2S1 | |
| PENK | SDK2 | DACT1 | |
| ANO1 | GPR162 | DNAJC18 | |
| CHGB | VPS37D | DPYSL2 | |
| CLIC5 | CXADR | DUSP6 | |
| SEZ6L | TINAGL1 | EBF4 | |
| AC243967.1 | AMOTL2 | ECM1 | |
| IGDCC3 | RGL1 | EEPD1 | |
| STRA6 | SHH | EFNB3 | |
| SMIM10 | LIMD2 | EGR1 | |
| IGFBPL1 | TP53INP1 | EMID1 | |
| SHH | NUDT4 | EPPK1 | |
| FGF20 | YPEL1 | FGF20 | |
| SLC16A12 | CD9 | FILIP1L | |
| CYP1A1 | EBF4 | FLRT3 | |
| CXCL14 | SRPX2 | FN1 | |
| WNT6 | FOXC2 | GALNT5 | |
| ST6GAL2 | SGPP1 | GAS6 | |
| APOD | PAPLN | GJB4 | |
| FGL2 | CTXN1 | GLB1L2 | |

| | | |
|---|---|---|
| NID1 | RIMKLA | GPR161 |
| ZAP70 | ATXN1 | GPR173 |
| SCARA3 | SORT1 | GSN |
| BCAM | SLC4A11 | HAPLN3 |
| ITGA1 | SEMA4F | HSPG2 |
| TMEM35A | MMP24 | IGSF9 |
| GALNT5 | DBNDD1 | IL17RD |
| BPIFB1 | FAM84A | IMPDH1 |
| CD70 | BAMBI | KCNH3 |
| IL17RD | DTNA | KIRREL |
| FZD2 | CITED2 | KLK5 |
| RIMKLA | MGAT5 | KRT20 |
| PDCD1 | TGFA | LIMD2 |
| GSN | DNAJB2 | LRRC75A |
| HS3ST4 | THBS1 | MAPK8IP1 |
| MYH7B | TRIM29 | MEIS3 |
| KRT71 | OSBP2 | MEX3A |
| VSIG2 | IMPDH1 | MFGE8 |
| EFNB3 | GATA6 | MGAT5 |
| AFAP1L2 | EMP3 | MT1X |
| COL26A1 | FLVCR2 | MYL9 |
| TNNT1 | FAM171A2 | NACAD |
| PLA2G2A | PTPN21 | NGFR |
| SOD3 | RHOQ | NKD2 |
| DPP6 | RTN2 | OLFML2A |
| BGN | CCDC148 | ORAI2 |
| SPRR1A | ZCCHC12 | PAM |
| KRT74 | ABHD2 | PCDHB15 |
| SLC9A4 | PCDHGA9 | PCDHB2 |
| LINGO1 | PIK3R3 | PCDHB8 |
| ZNF608 | PCDHGC3 | PCDHGC3 |
| GALNT14 | EEPD1 | PDGFB |
| MATN3 | PCDHGB2 | PDLIM1 |
| MFGE8 | GOLM1 | PDLIM4 |
| UPK2 | SLC35E4 | PIK3IP1 |
| FILIP1L | FLG | PLCB1 |
| DKK4 | ACTN1 | PMAIP1 |
| CLEC11A | ABTB2 | PNMA2 |
| SLC35F2 | MFSD6 | PPL |
| CRIP2 | MPP2 | PROCR |
| FN1 | PLCB1 | PRRT4 |
| MAGEA8 | PDLIM7 | PTPN14 |
| PRRT4 | CAPS | RASSF4 |
| YPEL1 | RCAN3 | RCOR2 |
| DACT1 | HSPG2 | REEP2 |
| PYGL | KRT80 | RGL1 |
| NGFR | PCDHB8 | RIMKLA |

| | | |
|---|---|---|
| LYZ | LRRN2 | SALL2 |
| ASTN1 | DNAJC18 | SBK1 |
| FRMPD1 | ADAMTS14 | SCARA3 |
| RGL1 | C15orf52 | SCRN1 |
| SMIM24 | OSR2 | SEMA4F |
| MGAT3 | PPL | SERPINE2 |
| MAL2 | NPTXR | SHH |
| SCUBE2 | IQSEC2 | SHROOM3 |
| ALOX5 | KLF6 | SLC34A2 |
| EMID1 | CHRNA3 | SLC4A11 |
| FLRT3 | FLNA | SORT1 |
| TBC1D4 | PCDHA4 | STK32B |
| VSTM2L | DPYSL2 | SULF2 |
| TUBB2B | SLC29A4 | SYT11 |
| MDFI | SHC2 | TIMP2 |
| ZDHHC2 | SLFN12 | TMEM173 |
| TRDC | SH3RF1 | TNNT1 |
| KRT13 | LTBP4 | TP53INP1 |
| NLRP7 | DNER | TPM1 |
| FZD10 | TLE1 | TUBA1A |
| SOBP | MAP1B | VPS37D |
| SEMA6B | TMEM132A | VWA5B2 |
| C1orf64 | EMID1 | WNT6 |
| TUBA1A | RASSF4 | YPEL1 |
| MMP2 | REEP1 | ZDHHC2 |
| IGSF9 | DUSP6 | ZNF423 |
| PNMA2 | SUGCT | ZNF608 |
| XG | BCAM | |
| EBF4 | CERCAM | |
| TMEM132E | FZD6 | |
| SLC2A10 | MAPK8IP2 | |
| C1QTNF5 | ZNF608 | |
| AC120057.3 | TUBB3 | |
| CD8B | ABHD6 | |
| CERCAM | PCDHB11 | |
| PLAU | VCL | |
| CDH3 | VWA5B2 | |
| TM4SF18 | SMURF2 | |
| KRT75 | SULF2 | |
| ARSJ | STMN3 | |
| NRGN | NAP1L3 | |
| TDGF1 | DUSP8 | |
| ADGRF1 | CACNA1S | |
| PAM | FAM13B | |
| ANOS1 | FLRT3 | |
| HAPLN3 | CCDC74A | |
| CCNJL | PCDHB14 | |

| | |
|---|---|
| CDH22 | RAB6B |
| CTSH | INHBB |
| GPR155 | SHROOM3 |
| FBN2 | FOXO6 |
| CEACAM6 | CPNE2 |
| ABHD12B | ABCC4 |
| ITGA10 | C12orf75 |
| MAP7D2 | FBXL2 |
| TRABD2B | PCDHB15 |
| HPSE | SERINC2 |
| LY6G6C | WNT10A |
| CERS6 | SOX4 |
| DLX2 | TNFRSF12A |
| MGAM2 | KIRREL |
| EPHA1 | SPON2 |
| JPH2 | ORAI2 |
| KIRREL | CMTM3 |
| ANGPTL2 | TMEM158 |
| ITGA3 | UNC5B |
| COMP | ZNF185 |
| SCD5 | PCDHB2 |
| COL13A1 | PCDHB7 |
| COL5A1 | GLI2 |
| PCDHA6 | MBOAT2 |
| HABP2 | SHANK1 |
| FAM46C | CRISPLD2 |
| LY6E | TMEM173 |
| ADGRF4 | PCDHB6 |
| SPOCK1 | ZNF423 |
| DIO1 | TANC2 |
| EFEMP2 | CDK5R1 |
| COCH | SERPINE2 |
| CIB2 | TNFRSF21 |
| CACNA1S | SALL2 |
| KRT20 | PDGFA |
| WLS | RGCC |
| KLK5 | SCRN1 |
| GPR173 | RPS6KA2 |
| TNS4 | MEIS3 |
| SPTSSB | TMSB4X |
| PTPRN2 | ASAP3 |
| FAIM2 | FAM110B |
| SYT11 | TPM1 |
| SEMA5B | APLP1 |
| TMEM178B | MAP2 |
| SCRN1 | DACT1 |
| AMOT | REEP2 |

| | |
|---|---|
| CRABP2 | RIMS3 |
| SSTR2 | PRRT4 |
| C5 | PIK3IP1 |
| ANK1 | PROCR |
| AQP3 | PLAT |
| SLC34A2 | MYO5A |
| NT5E | MFAP2 |
| LIPC | PHLDB1 |
| FXYD5 | ADAMTS15 |
| TMEM173 | VIM |
| RASGRF1 | LRRC75A |
| CUBN | IGSF9B |
| BMF | PTPN14 |
| P2RX4 | HEG1 |
| FUT1 | MAPK8IP1 |
| IL1RN | MEX3B |
| C1QL1 | AHNAK |
| STXBP6 | LRRC49 |
| SALL2 | RCOR2 |
| KCNA2 | IL17RD |
| GLB1L2 | TUBB6 |
| KCNH3 | COL6A1 |
| SLC4A11 | MICAL1 |
| NTSR1 | LRP12 |
| MMP7 | SEPT5 |
| EHD3 | NKD2 |
| NREP | KLF9 |
| SARDH | TUBB2A |
| NACAD | CACNB3 |
| FLI1 | PHLDA3 |
| THSD1 | ADGRB2 |
| MGLL | MT1X |
| CA11 | GAS6 |
| CYP1B1 | NKAIN1 |
| CEACAM5 | TPBG |
| VWA5B2 | GLCCI1 |
| LRRC2 | TNNT1 |
| CSRNP3 | PIFO |
| GPR161 | ADORA1 |
| CLDN5 | CRIP2 |
| PMAIP1 | CHST3 |
| REEP2 | TBC1D9 |
| SULF2 | GALNT5 |
| STK32B | MRAS |
| SEMA3C | EFR3B |
| PARM1 | UGT8 |
| PHGDH | SEMA7A |

| | |
|---|---|
| SLCO3A1 | EGR1 |
| CACNA2D2 | DLG3 |
| ARHGAP8 | MEX3A |
| B3GALNT1 | ATP2A3 |
| LAMC2 | PDLIM4 |
| C14orf105 | CDKN1C |
| F2R | NDRG4 |
| EPHB3 | PLK2 |
| KRT5 | FN1 |
| FIBCD1 | EPPK1 |
| CGNL1 | TMEM255A |
| RASSF4 | MT2A |
| ATP2C2 | CAMK1D |
| PDGFB | PCDHB10 |
| ADGRF5 | STK32B |
| FER1L6 | BMP7 |
| TRIB2 | TPPP3 |
| KCNIP3 | ANXA6 |
| CLGN | GLB1L2 |
| OTC | TIMP2 |
| FMOD | RUNX2 |
| COL6A3 | PDLIM1 |
| CPVL | TIMP3 |
| PLTP | PMAIP1 |
| MERTK | GJB4 |
| CYBRD1 | MYL9 |
| HSPG2 | GJA3 |
| AADAC | MPZL2 |
| SBK1 | TGFB2 |
| DKK1 | KRT20 |
| SLC16A2 | PAK1 |
| MT1X | KCNH3 |
| PTPRE | KCNN1 |
| EPHA7 | TAGLN3 |
| TRPM8 | ST8SIA2 |
| KLHL29 | MFGE8 |
| APOH | GLIPR1 |
| AMIGO2 | IGFL2 |
| ARNT2 | AMOT |
| C1orf115 | PAM |
| EGR1 | LFNG |
| TIMP2 | KLK5 |
| PROM2 | ANK1 |
| ZNF423 | NACAD |
| SV2A | ESM1 |
| FOXJ1 | WNT9A |
| KRT6B | AHNAK2 |

| | |
|---|---|
| KCNK9 | GPR173 |
| KIAA1324L | FILIP1L |
| COL17A1 | B3GNT7 |
| SLC29A1 | SYNGR3 |
| SERPINE2 | ANTXR2 |
| ELFN2 | SLC34A2 |
| ZNF618 | SYT11 |
| PCDHB5 | LTBP1 |
| MEGF6 | GSN |
| OSBPL6 | ANGPTL2 |
| SPNS2 | GPR161 |
| PTN | ACTA2 |
| MRC2 | FGF20 |
| PIK3AP1 | AHRR |
| TNXB | PCSK5 |
| DHRS9 | HDAC5 |
| UBASH3B | ZDHHC2 |
| TWSG1 | CREB5 |
| SYNPR | HAPLN3 |
| CAPN6 | CLIP3 |
| SRRM3 | CLSTN2 |
| HSPA2 | SBK1 |
| TMEM98 | PNMA2 |
| SLC14A1 | LOXL2 |
| MEIS3 | EFNB3 |
| VSIG10L | TMEM59L |
| ASAP3 | GLIS3 |
| PDP1 | TMEFF1 |
| ADAMTS9 | NPC2 |
| SPAG1 | GPR1 |
| ARMCX2 | CALHM1 |
| LCK | TMCC2 |
| CLDN10 | SPARC |
| EPS8L1 | AMIGO2 |
| APOBEC3B | OLFML2A |
| LIMD2 | AQP1 |
| SEMA4F | CD8B |
| DUSP6 | ZDHHC22 |
| CHN2 | SCARA3 |
| APBB1 | ISM1 |
| SYNE3 | PDGFB |
| LEF1 | ADAMTS7 |
| MMP11 | CLIC5 |
| GPRC5B | LTBP2 |
| PKN1 | TUBA1A |
| EEPD1 | MT1F |
| HEPHL1 | WNT6 |

| | |
|---|---|
| SHROOM3 | BMF |
| SNED1 | TMEM265 |
| DNAH2 | KRTAP2-3 |
| CCDC80 | KRTAP3-1 |
| KIAA1211L | |
| SCARF2 | |
| ECM1 | |
| CHST3 | |
| SRD5A2 | |
| UPK3B | |
| FRAS1 | |
| MYL9 | |
| C15orf52 | |
| CMTM3 | |
| CALHM2 | |
| DOCK11 | |
| VIPR1 | |
| DNAJC18 | |
| CLIP3 | |
| RCOR2 | |
| FBLN1 | |
| ARFGEF3 | |
| TM4SF4 | |
| B3GNT7 | |
| RAC2 | |
| HES6 | |
| MRAP2 | |
| EPB41L2 | |
| CYP39A1 | |
| LRRC75A | |
| OTOG | |
| VPS37D | |
| AGT | |
| GAS6 | |
| TNC | |
| LIPG | |
| IL15RA | |
| FHOD3 | |
| CACNB3 | |
| PCDHB15 | |
| FMR1NB | |
| SBSPON | |
| OLFML2A | |
| GATM | |
| ZDHHC14 | |
| PHC1 | |
| TGFB1I1 | |

| |
|---|
| SLC41A2 |
| PCDHB8 |
| DCBLD2 |
| PTGS1 |
| THBS3 |
| ARAP3 |
| PCNX2 |
| ENHO |
| PCDHB16 |
| KLK10 |
| MEX3A |
| PIK3IP1 |
| AHRR |
| TSPAN9 |
| KREMEN2 |
| HOXC5 |
| PPP1R1B |
| CPNE2 |
| NOS3 |
| RASSF3 |
| MAOB |
| ZNF22 |
| SETBP1 |
| TMEM125 |
| F5 |
| FAM174B |
| PLXNC1 |
| PROCR |
| LRRN4 |
| PDLIM4 |
| TP53INP1 |
| NKD2 |
| RTN4RL2 |
| PLCB1 |
| NEURL1B |
| ENTPD3 |
| PAQR8 |
| DPYSL2 |
| CLDN3 |
| QSOX1 |
| DISC1 |
| FLNC |
| NIN |
| TCEA3 |
| C16orf45 |
| PRSS8 |
| GNAL |

| |
|---|
| DNAAF3 |
| CHST7 |
| AKNA |
| FAM169A |
| GPD1L |
| MYO1D |
| SLC9A2 |
| CD302 |
| LARGE2 |
| MGAT5 |
| TGFBI |
| VASH1 |
| STEAP3 |
| MAPK8IP1 |
| LIPH |
| MMD |
| PPL |
| SLC2A6 |
| GJB4 |
| SFRP5 |
| SMPD1 |
| EPPK1 |
| EMP1 |
| TXNIP |
| TPBGL |
| MAP3K8 |
| PCDHGC3 |
| CYTH3 |
| PDLIM1 |
| CYP27A1 |
| IMPDH1 |
| MACC1 |
| EVC |
| SLC2A3 |
| TPPP |
| PKDCC |
| SH2D4A |
| CABLES1 |
| SLC2A12 |
| MCUB |
| CD59 |
| AC241585.3 |
| STBD1 |
| PCSK9 |
| CAPN5 |
| N4BP2 |
| NUP210 |

| |
|---|
| SORT1 |
| RAB36 |
| CTHRC1 |
| ARL4C |
| MUC1 |
| CTSV |
| ZSWIM5 |
| PTPN14 |
| PPM1H |
| AP3M2 |
| TTC39B |
| PXK |
| LARP6 |
| ABTB1 |
| CYP2S1 |
| GDPD5 |
| ORAI2 |
| MLXIPL |
| EPHA10 |
| KREMEN1 |
| LIMA1 |
| TPM1 |
| CHRNA5 |
| PCDHB2 |
| PTPRG |
| HES1 |
| MYO6 |

**Supplementary Table 6A.** Genes modulated by BLU0588, compared with FLC-related genes in the Simon et al dataset

| log2FC_Pos_Simon | Neg in BLU0588 | Overlap |
|---|---|---|
| MARCH4 | ROPN1L | CPS1 |
| MARCH10 | SCARA5 | KRT86 |
| AACS | BPIFA1 | STC1 |
| AAGAB | CYP24A1 | INSL4 |
| AAK1 | C11orf86 | PTPN5 |
| ABCA12 | SLC2A9 | CHST8 |
| ABCA2 | MAPK4 | GALNTL6 |
| ABCA3 | RIMKLB | NTS |
| ABCA7 | IGFBP1 | PPP1R3G |
| ABCB5 | CD55 | ACKR3 |
| ABCB8 | TSNAX-DISC1 | GK |
| ABCC5 | SPATA31A6 | PDE10A |
| ABCF2 | CPS1 | FAM155A |
| ABHD17C | KRT86 | HOPX |
| ABHD4 | PTGS2 | BMP6 |
| ABLIM2 | STC1 | FOXC1 |
| AC006132.1 | CYSLTR1 | EVA1C |
| AC008443.1 | INSL4 | KRT81 |
| AC010547.9 | PTPN5 | RHOBTB1 |
| AC096677.1 | CHST8 | PDE3A |
| AC145676.2 | SCNN1B | TESC |
| ACAN | SPATA31A1 | NOTCH3 |
| ACKR3 | ASPG | CPLX1 |
| ACLY | SPATA31A7 | KRT7 |
| ACSL4 | DUSP1 | FA2H |
| ACYP1 | TNFSF11 | SLC22A11 |
| ADAM12 | ITM2A | FST |
| ADAM22 | GALNTL6 | RASD1 |
| ADAM32 | LRRC32 | SLC2A1 |
| ADAMDEC1 | ANGPTL3 | SLC5A6 |
| ADAMTS14 | KCNK3 | C10orf90 |
| ADAMTS15 | MPP1 | PPARGC1A |
| ADAMTS18 | ANXA10 | PDE4B |
| ADAMTS6 | NTS | NR4A2 |
| ADAMTS9 | RBP4 | RGCC |
| ADAMTSL5 | GALNT8 | ATP1B1 |
| ADAT1 | PPP1R3G | VSIG1 |
| ADAT2 | ITIH4 | LGALS3 |
| ADCY2 | CU639417.2 | GABRQ |
| ADD2 | ACKR3 | EFCAB12 |
| ADRA2A | OXGR1 | CABYR |
| ADSSL1 | BASP1 | HHLA3 |
| AEN | NR4A1 | PBK |
| AF131215.5 | SIK1 | SPATA17 |
| AFAP1 | SMIM9 | SLC16A11 |
| AGA | KRT83 | AKR1B15 |
| AGR2 | TFCP2L1 | FSTL4 |
| AGRN | G6PC | GPRIN3 |
| AHI1 | SLC5A7 | DIO2 |
| AHNAK2 | GK | KCNU1 |
| AHRR | PTPRN | S100P |
| AIFM2 | BMP2 | KYNU |
| AIM1L | MFAP5 | CSGALNACT1 |
| AK8 | PDK4 | RASSF9 |
| AKAP12 | C11orf96 | AKAP12 |
| AKR1B15 | C3orf80 | ADSSL1 |
| AKR1C3 | CD247 | CPLX2 |
| AKTIP | DCLK1 | PTPRM |

| | | |
|---|---|---|
| AL133373.1 | VGF | IRX3 |
| AL358813.2 | ADAMTS12 | RHOF |
| AL359878.1 | ENTPD8 | SDCBP2 |
| ALDH18A1 | SCUBE1 | MUC13 |
| ALDH1A2 | ADH4 | HPD |
| ALDH1L2 | PDE10A | SEMA3F |
| ALDOA | PSG5 | URB1 |
| ALG1 | APOA5 | DAB2 |
| ALPK3 | FAM155A | DNM3 |
| AMFR | HOPX | MYO1E |
| ANKRD18A | PLA2G4A | MN1 |
| ANKRD22 | ALOX15B | MID1 |
| ANKRD29 | PLIN1 | HHIPL2 |
| ANKRD52 | NR4A3 | RAB3B |
| ANKS6 | HEPACAM | ADAM12 |
| ANLN | CLU | FOXL1 |
| ANO2 | MAMLD1 | TLCD2 |
| ANO4 | CA2 | LRP8 |
| ANTXR1 | CPB2 | DGKI |
| ANXA2 | BMP6 | CKB |
| ANXA5 | PCK1 | SMAD2 |
| AP1S3 | FOXC1 | ISG20 |
| APCDD1L | KRT17 | HES4 |
| APLN | EVA1C | PITX1 |
| APLP1 | NEFL | PBX1 |
| APOO | KRT81 | ARG2 |
| AQPEP | RHOBTB1 | MAFK |
| AREG | S1PR1 | CA12 |
| ARG2 | NUGGC | FAM167A |
| ARHGAP11A | GBP4 | DDIT4 |
| ARHGAP11B | RBFOX3 | OAT |
| ARHGAP18 | TSNAX | |
| ARHGAP22 | INHA | |
| ARHGAP36 | PDE3A | |
| ARHGAP39 | IGFBP3 | |
| ARHGEF35 | TESC | |
| ARHGEF39 | NOTCH3 | |
| ARHGEF5 | CPLX1 | |
| ARL14 | SLC16A6 | |
| ARL2BP | KRT7 | |
| ARMC9 | FA2H | |
| ARNTL2 | CRYAB | |
| AS3MT | KRT16 | |
| ASB15 | HRH2 | |
| ASIC1 | PLOD2 | |
| ASNS | SLC22A11 | |
| ASPHD1 | GOLGA6D | |
| ASRGL1 | PABPC1L2A | |
| ATIC | FXYD4 | |
| ATP1B1 | FST | |
| ATP2A1 | FXYD2 | |
| ATP2B4 | RASD1 | |
| ATP6V1B1 | SLC2A1 | |
| ATP8A2 | NPR3 | |
| ATP8B3 | PEX11A | |
| ATR | SLC5A6 | |
| AURKA | C10orf90 | |
| B3GNT3 | KCNJ16 | |
| B3GNT5 | P4HA3 | |
| B3GNTL1 | HEY1 | |
| B4GALNT1 | PGC | |

| | |
|---|---|
| B9D1 | PPARGC1A |
| BAG2 | UGT1A4 |
| BAG3 | GRIP2 |
| BAI2 | APOA4 |
| BAIAP2L2 | ID4 |
| BBC3 | UGT2B15 |
| BBS2 | LYPD3 |
| BCAR1 | GPT |
| BCAT1 | GDF7 |
| BCAT2 | PSG6 |
| BCL11A | PDE7B |
| BCL2L2-PABPN1 | ODC1 |
| BDKRB1 | PDE4B |
| BDKRB2 | KCNJ15 |
| BDNF | PCP4 |
| BEND3 | NR4A2 |
| BEND6 | RGCC |
| BEST3 | IRS2 |
| BFSP1 | ATP1B1 |
| BICD1 | VSIG1 |
| BIRC5 | LGALS3 |
| BLM | ALDH1L1 |
| BLVRA | LRRK1 |
| BMP6 | GABRQ |
| BMP8A | BLOC1S5-TXNDC5 |
| BMP8B | EFCAB12 |
| BNC2 | CABYR |
| BRE | HHLA3 |
| BRSK2 | HYAL1 |
| BSG | PSTPIP2 |
| BUB1 | PTP4A1 |
| BUB1B | PER2 |
| C10orf128 | CRABP1 |
| C10orf131 | EPAS1 |
| C10orf2 | F2RL2 |
| C10orf90 | PBK |
| C12orf39 | SPATA17 |
| C12orf5 | UGT1A1 |
| C14orf183 | SLC16A11 |
| C15orf65 | FGA |
| C16orf46 | CPA4 |
| C16orf59 | GABRP |
| C16orf93 | SYT12 |
| C18orf56 | ANGPTL4 |
| C19orf54 | AKR1B15 |
| C1orf105 | SLC13A3 |
| C1orf198 | BCAS1 |
| C1orf229 | KHK |
| C1QL1 | AVPI1 |
| C20orf96 | FSTL4 |
| C2CD4A | PTPRB |
| C2orf27A | SGK1 |
| C2orf66 | CXCL5 |
| C2orf81 | DNAJC12 |
| C3orf36 | GPRIN3 |
| C3orf52 | DIO2 |
| C3orf67 | CCDC68 |
| C4orf47 | KCNU1 |
| C5AR2 | S100P |
| C5orf46 | KYNU |
| C6orf163 | GJA1 |

| | |
|---|---|
| C6orf164 | CSGALNACT1 |
| C6orf195 | B3GALT1 |
| C8G | RASSF9 |
| C8orf87 | UGT1A3 |
| C9orf116 | HYAL3 |
| C9orf163 | GGH |
| C9orf57 | TCF7L1 |
| C9orf66 | ZBED2 |
| CA12 | AKAP12 |
| CA5A | FABP5 |
| CA5B | ADSSL1 |
| CA8 | CPLX2 |
| CA9 | WNT11 |
| CABYR | PTPRM |
| CACNA1C | RAB27B |
| CACNA1D | SLC11A1 |
| CACNB2 | IRX3 |
| CACNB4 | RHOF |
| CALCA | CPEB4 |
| CALCB | GADD45B |
| CAMK2N2 | SDCBP2 |
| CAP2 | MUC13 |
| CAPN11 | PDZRN3 |
| CATSPERB | SLC6A12 |
| CBX2 | CH25H |
| CBX8 | HPD |
| CC2D2B | RPL17-C18orf32 |
| CCDC102B | MYOM2 |
| CCDC113 | C4B |
| CCDC13 | CST1 |
| CCDC136 | AKR1C2 |
| CCDC169 | CST7 |
| CCDC170 | HMOX1 |
| CCDC177 | SPP1 |
| CCDC64 | CST2 |
| CCDC78 | CXCL12 |
| CCDC80 | FMO5 |
| CCDC85A | RASAL1 |
| CCDC88C | BHLHE40 |
| CCNA2 | DUXA |
| CCNB1 | SEMA3F |
| CCNB2 | CMIP |
| CCNE1 | PREB |
| CCNF | PAH |
| CCNO | TFPI |
| CCR8 | SLC4A4 |
| CCRN4L | UGT2A3 |
| CCT3 | ABCA1 |
| CCT5 | ONECUT2 |
| CCT6A | FMNL1 |
| CD109 | LATS2 |
| CD200 | PLPP1 |
| CD248 | PGM2L1 |
| CD34 | PC |
| CD46 | URB1 |
| CDC20 | FSTL3 |
| CDC20B | DAB2 |
| CDC25C | PLAC8 |
| CDC45 | IL10RA |
| CDC6 | IRX6 |
| CDCA2 | PRKD3 |

| | |
|---|---|
| CDCA5 | GCH1 |
| CDCA7 | FERMT1 |
| CDH11 | TTPA |
| CDH13 | ZFP36 |
| CDH17 | OSBP2 |
| CDH24 | DNM3 |
| CDH6 | ANXA1 |
| CDK1 | HHAT |
| CDK6 | MYO1E |
| CDKN2A | TBX3 |
| CDKN2B | SPANXD |
| CDKN3 | MN1 |
| CDX1 | ENO3 |
| CDYL2 | CBR3 |
| CELSR3 | MID1 |
| CEND1 | DGKD |
| CENPF | SPANXA1 |
| CENPI | CREG1 |
| CENPK | CASP9 |
| CENPL | HHIPL2 |
| CENPO | RND1 |
| CEP128 | USP28 |
| CEP152 | CHCHD10 |
| CEP55 | NAT6 |
| CERCAM | APOB |
| CERKL | RAB3B |
| CGA | TBX4 |
| CGREF1 | KLB |
| CHAC1 | ITGB8 |
| CHCHD3 | ADAM12 |
| CHEK1 | DNAJC15 |
| CHI3L1 | FOXL1 |
| CHML | TLCD2 |
| CHN1 | ATF3 |
| CHRDL2 | SQSTM1 |
| CHRNA5 | LRP8 |
| CHST10 | SRXN1 |
| CHST6 | DGKI |
| CHST8 | CDHR5 |
| CIAPIN1 | CKB |
| CIDEC | MSLN |
| CKB | TGFBR3L |
| CKS2 | RAB10 |
| CLCF1 | SMAD2 |
| CLDN5 | PIM3 |
| CLEC18B | B4GALT1 |
| CLGN | ISG20 |
| CLIC5 | PLA2G4F |
| CLIP4 | HES4 |
| CLSPN | PAG1 |
| CMTM4 | PITX1 |
| CNNM2 | BRD4 |
| COL10A1 | TP53INP2 |
| COL11A1 | BAG1 |
| COL15A1 | KCNMB4 |
| COL17A1 | PELI2 |
| COL1A1 | SIAH2 |
| COL1A2 | PBX1 |
| COL22A1 | HGD |
| COL4A1 | ARG2 |
| COL4A2 | UGT2B17 |

| | |
|---|---|
| COL5A1 | C1QTNF6 |
| COL5A2 | CYP2C18 |
| COL5A3 | HSPB8 |
| COL6A3 | LARGE1 |
| COL6A5 | MAFK |
| COL8A1 | CA12 |
| COL9A1 | FAS |
| COMP | AKR1C1 |
| COQ9 | MESP1 |
| CORIN | CNKSR3 |
| CORO2A | SCN1B |
| COX4I2 | FAM167A |
| CPA6 | PRICKLE2 |
| CPD | DDIT4 |
| CPE | DDIT3 |
| CPLX1 | PPP2R1B |
| CPLX2 | GCLC |
| CPNE5 | OAT |
| CPNE7 | ITPRIP |
| CPS1 | NTRK2 |
| CPXM1 | KCNE3 |
| CRCP | |
| CREB3L1 | |
| CREG2 | |
| CRLF2 | |
| CRMP1 | |
| CRP | |
| CRYM | |
| CSGALNACT1 | |
| CSMD2 | |
| CSNK2A2 | |
| CSPG4 | |
| CTD-2600O9.1 | |
| CTH | |
| CTHRC1 | |
| CXCL16 | |
| CXorf36 | |
| CYBA | |
| CYCS | |
| CYP17A1 | |
| CYP19A1 | |
| CYP1B1 | |
| CYP21A2 | |
| CYP27B1 | |
| CYSTM1 | |
| DAB2 | |
| DAGLA | |
| DCAF4L1 | |
| DCDC2 | |
| DCLK2 | |
| DCX | |
| DDIT4 | |
| DDIT4L | |
| DEFB1 | |
| DEFB132 | |
| DEPDC1 | |
| DFNA5 | |
| DGKI | |
| DHRS7 | |
| DIAPH3 | |
| DIO2 | |

| |
|---|
| DKK1 |
| DKK2 |
| DKK4 |
| DLGAP4 |
| DLGAP5 |
| DNAAF1 |
| DNAAF3 |
| DNAH17 |
| DNAH5 |
| DNAH7 |
| DNAJC6 |
| DNER |
| DNM3 |
| DOCK5 |
| DOK4 |
| DOK5 |
| DPCD |
| DPEP1 |
| DPP10 |
| DPY19L1 |
| DPYSL4 |
| DSEL |
| DTL |
| DTNA |
| DUSP4 |
| DUSP8 |
| DYDC2 |
| E2F1 |
| E2F3 |
| E2F5 |
| E2F7 |
| E2F8 |
| EARS2 |
| EBF1 |
| EBF2 |
| EBF3 |
| ECEL1 |
| ECT2 |
| EDA2R |
| EDIL3 |
| EEF1A2 |
| EFCAB12 |
| EFEMP1 |
| EFNA3 |
| EFNA5 |
| EGF |
| EGFL6 |
| EGLN3 |
| EHD2 |
| EIF4G3 |
| ELOVL3 |
| ELOVL7 |
| EMC7 |
| EME1 |
| ENDOD1 |
| ENO1 |
| ENOX1 |
| ENTPD1 |
| EPB41L1 |
| EPDR1 |
| EPHA10 |

| |
|---|
| EPHX4 |
| EPS8L3 |
| ERBB2 |
| ERC2 |
| ERCC6L |
| EREG |
| ERMP1 |
| ESCO2 |
| ESM1 |
| ESRRG |
| ETV1 |
| ETV4 |
| EVA1C |
| EVC |
| EXO1 |
| EYS |
| EZH2 |
| EZR |
| F13A1 |
| F2RL1 |
| F2RL3 |
| FA2H |
| FAHD2B |
| FAM101A |
| FAM111B |
| FAM115C |
| FAM127A |
| FAM127B |
| FAM127C |
| FAM132B |
| FAM135A |
| FAM155A |
| FAM155B |
| FAM167A |
| FAM169B |
| FAM177B |
| FAM179A |
| FAM188B |
| FAM189B |
| FAM196A |
| FAM19A5 |
| FAM227A |
| FAM64A |
| FAM71F2 |
| FAM78B |
| FAM81A |
| FAM83A |
| FAM86A |
| FAM86B1 |
| FANCI |
| FAP |
| FASTKD1 |
| FAT1 |
| FBN1 |
| FBXL18 |
| FBXL8 |
| FBXO25 |
| FBXO36 |
| FBXO39 |
| FGF7 |
| FHDC1 |

| |
|---|
| FKBP11 |
| FKBP14 |
| FKBP1C |
| FLAD1 |
| FLVCR1 |
| FMNL2 |
| FMNL3 |
| FNDC1 |
| FNDC3A |
| FOXC1 |
| FOXF2 |
| FOXL1 |
| FOXM1 |
| FOXQ1 |
| FOXS1 |
| FRRS1L |
| FRZB |
| FSD1L |
| FST |
| FSTL4 |
| FTO |
| FURIN |
| FZD10 |
| FZD2 |
| FZD6 |
| FZD8 |
| G6PD |
| GABRB3 |
| GABRD |
| GABRE |
| GABRQ |
| GAD1 |
| GAL3ST4 |
| GALNT10 |
| GALNT12 |
| GALNT5 |
| GALNT7 |
| GALNTL6 |
| GALR2 |
| GAPDH |
| GAREML |
| GARS |
| GAS8 |
| GATSL2 |
| GCNT3 |
| GDA |
| GDF15 |
| GGN |
| GHRHR |
| GINS1 |
| GJA5 |
| GJC1 |
| GK |
| GLA |
| GLCE |
| GLIS3 |
| GLP1R |
| GLRB |
| GLRX |
| GLS |
| GMDS |

| |
|---|
| GNA12 |
| GNAL |
| GNAZ |
| GNB5 |
| GNG4 |
| GOLGA6L9 |
| GOLM1 |
| GOLT1B |
| GOT1 |
| GPATCH4 |
| GPC2 |
| GPD1L |
| GPI |
| GPR1 |
| GPR107 |
| GPR150 |
| GPR37 |
| GPR4 |
| GPR56 |
| GPR63 |
| GPR64 |
| GPR68 |
| GPR97 |
| GPRIN1 |
| GPRIN3 |
| GPSM2 |
| GPX8 |
| GRAMD1A |
| GRAMD1B |
| GRAMD4 |
| GREM2 |
| GRIA3 |
| GRK7 |
| GRM5 |
| GRM7 |
| GSDMC |
| GSN |
| GTF2E1 |
| GTF2IRD1 |
| GTPBP4 |
| GUCY2C |
| GULP1 |
| GXYLT2 |
| H2AFY2 |
| HAGHL |
| HAPLN1 |
| HAS3 |
| HAVCR1 |
| HCN1 |
| HEATR2 |
| HEG1 |
| HELLS |
| HES2 |
| HES4 |
| HEYL |
| HHATL |
| HHIPL2 |
| HHLA3 |
| HIGD1B |
| HIST1H1A |
| HIST1H1C |

| |
|---|
| HIST1H1T |
| HIST1H2AA |
| HIST1H2AB |
| HIST1H2AC |
| HIST1H2AG |
| HIST1H2AH |
| HIST1H2AI |
| HIST1H2AK |
| HIST1H2BB |
| HIST1H2BG |
| HIST1H2BJ |
| HIST1H2BK |
| HIST1H2BN |
| HIST1H3A |
| HIST1H3D |
| HIST1H4I |
| HIST2H3D |
| HJURP |
| HK2 |
| HKDC1 |
| HMGB3 |
| HNF1B |
| HOMER1 |
| HOPX |
| HOXD1 |
| HOXD3 |
| HOXD9 |
| HPD |
| HSF4 |
| HSP90AA1 |
| HSP90AB1 |
| HSPA12A |
| HSPA12B |
| HSPA4L |
| HSPA5 |
| HSPB6 |
| HSPH1 |
| HTATIP2 |
| HTR2A |
| HTRA1 |
| HYOU1 |
| IARS |
| ICAM5 |
| IER3 |
| IER5L |
| IGDCC4 |
| IGF2BP3 |
| IGFBPL1 |
| IGSF9B |
| IL17D |
| IL2RA |
| IL31RA |
| IL32 |
| INPP4B |
| INSL4 |
| INTS9 |
| INTU |
| IQCA1 |
| IQCD |
| IQCE |
| IQCK |

| |
|---|
| IQGAP3 |
| IRAK1 |
| IRF4 |
| IRX3 |
| IRX5 |
| ISG15 |
| ISG20 |
| ITGA11 |
| ITGA2 |
| ITGA6 |
| ITGA7 |
| ITGAV |
| ITGB4 |
| ITPKA |
| JAG1 |
| JAG2 |
| JPH1 |
| KAAG1 |
| KAL1 |
| KCNE1L |
| KCNE4 |
| KCNF1 |
| KCNJ5 |
| KCNJ6 |
| KCNK9 |
| KCNN3 |
| KCNQ3 |
| KCNU1 |
| KDELR3 |
| KIAA0100 |
| KIAA0319 |
| KIAA0556 |
| KIAA0895L |
| KIAA1024 |
| KIAA1199 |
| KIAA1211L |
| KIAA1244 |
| KIAA1324 |
| KIAA1462 |
| KIAA1522 |
| KIAA1549 |
| KIAA1549L |
| KIF14 |
| KIF18B |
| KIF20A |
| KIF21B |
| KIF23 |
| KIF24 |
| KIF26B |
| KIF2C |
| KIF3A |
| KIF4A |
| KIF5A |
| KLC2 |
| KLF5 |
| KLHDC7B |
| KLHL21 |
| KLHL29 |
| KNDC1 |
| KPNA2 |
| KRT222 |

| |
|---|
| KRT7 |
| KRT81 |
| KRT86 |
| KSR1 |
| KYNU |
| LAMA1 |
| LAMA3 |
| LAMA4 |
| LAMB4 |
| LAMC1 |
| LANCL1 |
| LANCL3 |
| LAPTM4B |
| LBP |
| LCN2 |
| LDHB |
| LEF1 |
| LEP |
| LETM2 |
| LGALS3 |
| LGI2 |
| LHCGR |
| LHFPL5 |
| LHX6 |
| LIF |
| LIMCH1 |
| LIMK1 |
| LIMK2 |
| LIPH |
| LITAF |
| LOH12CR1 |
| LONRF2 |
| LOX |
| LOXL2 |
| LPAR3 |
| LPAR4 |
| LPCAT1 |
| LPL |
| LPPR4 |
| LRP8 |
| LRRC1 |
| LRRC16A |
| LRRC37A3 |
| LRRC69 |
| LRRC73 |
| LRRC8E |
| LSAMP |
| LTBP2 |
| LUZP2 |
| LYPD1 |
| LZTS1 |
| MAB21L3 |
| MAFK |
| MALL |
| MANF |
| MAP1A |
| MAP1B |
| MAP1LC3B |
| MAP1LC3B2 |
| MAP3K9 |
| MAP9 |

| |
|---|
| MAPK12 |
| MAPT |
| MARS2 |
| MATN3 |
| MBOAT4 |
| MCAM |
| MCF2L2 |
| MCM10 |
| MCM4 |
| MCTP2 |
| MECOM |
| MED9 |
| MELK |
| MEX3B |
| MFSD6 |
| MGAT3 |
| MGAT5B |
| MGC4294 |
| MID1 |
| MKI67 |
| MLEC |
| MMP1 |
| MMP10 |
| MMP11 |
| MMP14 |
| MMP2 |
| MN1 |
| MND1 |
| MPHOSPH6 |
| MPP3 |
| MPP4 |
| MPV17 |
| MPV17L2 |
| MPZ |
| MRAP |
| MRAP2 |
| MRAS |
| MRPS12 |
| MRVI1 |
| MST1R |
| MT-ATP8 |
| MT-ND5 |
| MT-ND6 |
| MT3 |
| MTFR2 |
| MTHFD1L |
| MTRNR2L2 |
| MUC13 |
| MUC3A |
| MUC5B |
| MURC |
| MVP |
| MYBPHL |
| MYCN |
| MYEF2 |
| MYH4 |
| MYO1E |
| MYO5C |
| MYOF |
| MYOM3 |
| MYRF |

| |
|---|
| N4BP3 |
| NAV1 |
| NAV2 |
| NBEA |
| NCAPG |
| NCR3LG1 |
| NDUFA4L2 |
| NEB |
| NEIL3 |
| NEK2 |
| NETO2 |
| NHS |
| NIPAL2 |
| NKX1-2 |
| NKX2-3 |
| NME1 |
| NMNAT2 |
| NOL3 |
| NOMO1 |
| NOMO2 |
| NOMO3 |
| NOTCH3 |
| NOV |
| NOVA1 |
| NOX1 |
| NOX4 |
| NPAS2 |
| NPCDR1 |
| NPFFR2 |
| NPNT |
| NPTX2 |
| NPTXR |
| NPY5R |
| NQO1 |
| NR4A2 |
| NRG2 |
| NRIP2 |
| NRXN3 |
| NT5C3A |
| NT5DC2 |
| NTM |
| NTS |
| NUF2 |
| NXPH4 |
| OAT |
| OGDH |
| OGFOD1 |
| OLAH |
| OLFML2A |
| OLFML2B |
| OPHN1 |
| OPN3 |
| OR1F1 |
| OR2A7 |
| OR2AG2 |
| OR2B6 |
| OR2D2 |
| OR51E1 |
| OR51E2 |
| OR6A2 |
| ORC1 |

| |
|---|
| ORC6 |
| OSBPL3 |
| OSBPL9 |
| OSMR |
| OSR1 |
| OSR2 |
| OTOG |
| OTUB2 |
| OTUD7A |
| OTX1 |
| OXCT1 |
| P4HA2 |
| PAEP |
| PAK3 |
| PALLD |
| PAPPA |
| PAQR4 |
| PAQR5 |
| PARM1 |
| PARPBP |
| PBK |
| PBX1 |
| PCDH17 |
| PCDHB10 |
| PCDHB11 |
| PCDHB13 |
| PCDHB8 |
| PCDHGA1 |
| PCDHGA12 |
| PCDHGA4 |
| PCDHGA5 |
| PCDHGA7 |
| PCDHGA8 |
| PCDHGB1 |
| PCDHGB2 |
| PCDHGC5 |
| PCDP1 |
| PCED1B |
| PCM1 |
| PCNT |
| PCNXL2 |
| PCSK1 |
| PDE10A |
| PDE1C |
| PDE3A |
| PDE3B |
| PDE4A |
| PDE4B |
| PDE4D |
| PDF |
| PDGFA |
| PDGFRB |
| PDGFRL |
| PDK1 |
| PDLIM7 |
| PDX1 |
| PDZD2 |
| PFKFB2 |
| PFKM |
| PFKP |
| PHEX |

| |
|---|
| PHF17 |
| PHLDA2 |
| PHLDA3 |
| PHPT1 |
| PIAS3 |
| PIGA |
| PIP4K2C |
| PITPNM1 |
| PITX1 |
| PKP1 |
| PLA2G2A |
| PLA2G2C |
| PLA2G4E |
| PLCB4 |
| PLCD3 |
| PLCE1 |
| PLCH1 |
| PLEKHA8 |
| PLEKHG2 |
| PLEKHH1 |
| PLEKHN1 |
| PLK1 |
| PLOD3 |
| PLP2 |
| PLVAP |
| PLXDC1 |
| PM20D2 |
| PMEPA1 |
| PMFBP1 |
| PNMA1 |
| PODNL1 |
| PODXL |
| POLN |
| POLR1A |
| POLR2C |
| POLR3G |
| PON2 |
| POSTN |
| POU5F1 |
| PPA1 |
| PPARGC1A |
| PPAT |
| PPIA |
| PPP1R13L |
| PPP1R36 |
| PPP1R3D |
| PPP1R3G |
| PPP2R2C |
| PRAMEF10 |
| PRAMEF2 |
| PRAMEF4 |
| PRC1 |
| PRDM6 |
| PRDM7 |
| PRKACA |
| PRND |
| PRODH |
| PRR15 |
| PRR16 |
| PRR26 |
| PRRG3 |

| |
|---|
| PRRX1 |
| PRSS27 |
| PSMD14 |
| PSME3 |
| PSORS1C1 |
| PTCHD4 |
| PTGFR |
| PTGFRN |
| PTP4A3 |
| PTPDC1 |
| PTPLA |
| PTPN14 |
| PTPN5 |
| PTPRM |
| PTPRR |
| PTPRU |
| PVRL1 |
| PYCR1 |
| PYGB |
| QRFPR |
| RAB3A |
| RAB3B |
| RAB6B |
| RACGAP1 |
| RAET1E |
| RANBP17 |
| RAP1GAP |
| RARRES1 |
| RASD1 |
| RASD2 |
| RASEF |
| RASGEF1A |
| RASGRF2 |
| RASL11B |
| RASL12 |
| RASSF6 |
| RASSF9 |
| RBM20 |
| RBM24 |
| RBM44 |
| RBPMS |
| RECQL4 |
| RELL2 |
| RFX8 |
| RGAG4 |
| RGCC |
| RGS17 |
| RGS5 |
| RGS6 |
| RGS9 |
| RHBDD2 |
| RHOBTB1 |
| RHOBTB2 |
| RHOF |
| RHOQ |
| RHPN1 |
| RMI2 |
| RNF157 |
| RNFT2 |
| ROBO1 |
| ROR2 |

| |
|---|
| RP1-102H19.8 |
| RP11-17M16.1 |
| RP11-212D19.4 |
| RP11-248J23.6 |
| RP11-366L20.2 |
| RP11-796G6.2 |
| RPGR |
| RPGRIP1L |
| RPP40 |
| RPRML |
| RPS6KA2 |
| RPS6KL1 |
| RRM2 |
| RRS1 |
| RUNX1 |
| S100A3 |
| S100P |
| SAPCD2 |
| SATB2 |
| SCG2 |
| SCIN |
| SCML4 |
| SCN4A |
| SDCBP2 |
| SDIM1 |
| SDSL |
| SEMA3F |
| SEMA3G |
| SEMA5B |
| SERHL2 |
| SERPINB3 |
| SERTAD4 |
| SEZ6L2 |
| SFMBT2 |
| SFN |
| SFRP2 |
| SFRP4 |
| SGIP1 |
| SGPP2 |
| SH3PXD2B |
| SH3RF3 |
| SHC1 |
| SHCBP1 |
| SHISA2 |
| SHROOM4 |
| SIAE |
| SIX1 |
| SIX4 |
| SKA1 |
| SLC16A11 |
| SLC16A14 |
| SLC22A11 |
| SLC22A12 |
| SLC22A15 |
| SLC22A23 |
| SLC22A5 |
| SLC25A12 |
| SLC25A15 |
| SLC25A24 |
| SLC26A2 |
| SLC26A7 |

| |
|---|
| SLC26A9 |
| SLC27A4 |
| SLC2A1 |
| SLC2A5 |
| SLC35B4 |
| SLC35C1 |
| SLC35E4 |
| SLC35G2 |
| SLC36A1 |
| SLC38A1 |
| SLC44A3 |
| SLC45A1 |
| SLC4A11 |
| SLC52A3 |
| SLC5A6 |
| SLC6A11 |
| SLC6A17 |
| SLC6A3 |
| SLC6A6 |
| SLC6A8 |
| SLC6A9 |
| SLC7A1 |
| SLC7A11 |
| SLC7A2 |
| SLC7A5 |
| SLC7A6 |
| SLCO1C1 |
| SLCO2A1 |
| SLCO5A1 |
| SLIT2 |
| SLITRK4 |
| SMAD2 |
| SMCO2 |
| SMKR1 |
| SMOC2 |
| SMOX |
| SNAP25 |
| SNPH |
| SOGA1 |
| SORT1 |
| SOX11 |
| SOX12 |
| SOX21 |
| SOX4 |
| SP140 |
| SP6 |
| SPA17 |
| SPATA17 |
| SPATC1L |
| SPATS2 |
| SPECC1 |
| SPINK1 |
| SPIRE1 |
| SPOCD1 |
| SPOCK1 |
| SPRED3 |
| SPTBN5 |
| SPTSSA |
| SRGAP1 |
| SRM |
| SRPX2 |

| |
|---|
| SRRM3 |
| SRSF12 |
| ST6GALNAC4 |
| ST8SIA5 |
| STAMBPL1 |
| STC1 |
| STC2 |
| STEAP1 |
| STEAP1B |
| STK32B |
| STRA6 |
| STRIP2 |
| STS |
| STXBP1 |
| STXBP4 |
| STXBP5 |
| SULF1 |
| SULF2 |
| SULT1C2 |
| SULT2B1 |
| SULT4A1 |
| SYNDIG1 |
| SYT13 |
| TAF4B |
| TAF6 |
| TAF7L |
| TAGLN2 |
| TANC2 |
| TAS2R3 |
| TAS2R4 |
| TAS2R5 |
| TAX1BP3 |
| TBC1D16 |
| TBC1D30 |
| TBC1D31 |
| TC2N |
| TCF23 |
| TCN1 |
| TCTN3 |
| TEAD4 |
| TENM4 |
| TESC |
| TEX9 |
| TG |
| TGFB2 |
| TGFBR1 |
| TGIF1 |
| TGM2 |
| TGM3 |
| THBS2 |
| THBS4 |
| THSD7A |
| THY1 |
| TICAM2 |
| TICRR |
| TIMP1 |
| TK1 |
| TLCD1 |
| TLCD2 |
| TLDC1 |
| TLE6 |

| |
|---|
| TLL2 |
| TM4SF5 |
| TMC1 |
| TMC5 |
| TMC7 |
| TMED3 |
| TMEM104 |
| TMEM108 |
| TMEM119 |
| TMEM132A |
| TMEM136 |
| TMEM144 |
| TMEM145 |
| TMEM156 |
| TMEM163 |
| TMEM165 |
| TMEM178B |
| TMEM2 |
| TMEM200A |
| TMEM217 |
| TMEM245 |
| TMEM246 |
| TMEM45A |
| TMEM55A |
| TMEM59L |
| TMOD1 |
| TMTC2 |
| TNFRSF10A |
| TNFRSF12A |
| TNFRSF21 |
| TNFRSF4 |
| TNFSF4 |
| TNRC6C |
| TOMM40L |
| TOP2A |
| TOX |
| TPBG |
| TPBGL |
| TPM4 |
| TRAF2 |
| TRIM31 |
| TRIM40 |
| TRIM46 |
| TRIM59 |
| TRIM7 |
| TRIM8 |
| TRIP13 |
| TRPC4 |
| TRPC6 |
| TRPM2 |
| TRPS1 |
| TSHZ2 |
| TSPAN13 |
| TTC26 |
| TTC39A |
| TTK |
| TTL |
| TTYH2 |
| TUBA4A |
| TXNDC17 |
| TXNRD1 |

| |
|---|
| TYMS |
| UBE2C |
| UBE2T |
| UBFD1 |
| UBXN10 |
| UCHL1 |
| UCN |
| UGT2B11 |
| ULBP1 |
| ULBP2 |
| ULBP3 |
| UNC119B |
| UNC13A |
| UNC13B |
| UNC5B |
| UNC5D |
| URB1 |
| USB1 |
| USH1C |
| USP49 |
| USP54 |
| UTS2B |
| VARS |
| VASH2 |
| VAX2 |
| VCAN |
| VLDLR |
| VSIG1 |
| VSX1 |
| VWA7 |
| VWF |
| WDR13 |
| WIPF3 |
| WNT16 |
| WWC1 |
| XKR3 |
| XKR6 |
| XKRX |
| XPR1 |
| XRCC2 |
| YIPF6 |
| YWHAG |
| ZBTB38 |
| ZC3H12B |
| ZFHX3 |
| ZFP69B |
| ZHX1-C8ORF76 |
| ZIC2 |
| ZMAT3 |
| ZMYND15 |
| ZNF154 |
| ZNF215 |
| ZNF23 |
| ZNF233 |
| ZNF365 |
| ZNF382 |
| ZNF385D |
| ZNF57 |
| ZNF683 |
| ZNF695 |
| ZNF703 |

| |
|---|
| ZNF724P |
| ZNF827 |
| ZP3 |
| ZSCAN5A |
| ZSCAN9 |
| ZWINT |
| ZXDB |

**Supplementary Table 6B.** Genes modulated by BLU0588, compared with FLC-related genes in the Simon et al dataset

| log2FC_Neg_Simon | Pos in BLU0588 | Overlap |
|---|---|---|
| MARCH2 | CMKLR1 | OLFML3 |
| SEPT4 | MUCL1 | DBH |
| A1BG | CA4 | ADAMTSL2 |
| A1CF | RSPO4 | ANO1 |
| AADAC | OLFML3 | SEZ6L |
| AADAT | MMP3 | SHH |
| ABAT | CCM2L | SLC16A12 |
| ABCA13 | CST6 | CYP1A1 |
| ABCA6 | NXPH3 | CXCL14 |
| ABCA9 | DBH | FGL2 |
| ABCB11 | WNT16 | HS3ST4 |
| ABCG2 | VWA2 | MYH7B |
| ABCG5 | MMP12 | GALNT14 |
| ABCG8 | PNMT | YPEL1 |
| ABHD1 | ANKRD1 | PYGL |
| ABHD15 | ADAMTSL2 | NGFR |
| ABHD5 | LIX1 | ASTN1 |
| AC007405.2 | CARD11 | RGL1 |
| AC010368.2 | SERPINI1 | SOBP |
| AC011484.1 | WFDC2 | IGSF9 |
| AC011841.1 | LOXL4 | EBF4 |
| AC018755.1 | LYPD6B | CEACAM6 |
| AC021860.1 | CHRNA3 | TRABD2B |
| AC023590.1 | CTSE | EPHA1 |
| AC103801.2 | BMP7 | HABP2 |
| AC104667.3 | PENK | FAM46C |
| AC104809.3 | ANO1 | LY6E |
| AC110781.3 | CHGB | DIO1 |
| ACAA1 | CLIC5 | PTPRN2 |
| ACAA2 | SEZ6L | SSTR2 |
| ACACB | AC243967.1 | NT5E |
| ACADL | IGDCC3 | LIPC |
| ACADM | STRA6 | P2RX4 |
| ACADS | SMIM10 | IL1RN |
| ACADSB | IGFBPL1 | MMP7 |
| ACAT1 | SHH | EHD3 |
| ACAT2 | FGF20 | NREP |
| ACBD4 | SLC16A12 | SARDH |
| ACE2 | CYP1A1 | PHGDH |
| ACKR4 | CXCL14 | CACNA2D2 |
| ACOT1 | WNT6 | CGNL1 |
| ACOT12 | ST6GAL2 | OTC |
| ACOT13 | APOD | AADAC |
| ACOT2 | FGL2 | SLC16A2 |
| ACOT4 | NID1 | EPHA7 |
| ACOX2 | ZAP70 | TRPM8 |
| ACP5 | SCARA3 | APOH |
| ACSF2 | BCAM | C1orf115 |
| ACSL1 | ITGA1 | OSBPL6 |
| ACSL5 | TMEM35A | PTN |
| ACSL6 | GALNT5 | PIK3AP1 |
| ACSM2A | BPIFB1 | CLDN10 |
| ACSM2B | CD70 | LIMD2 |
| ACSM3 | IL17RD | CHN2 |
| ACSM5 | FZD2 | ECM1 |

| | | |
|---|---|---|
| ACSS2 | RIMKLA | FRAS1 |
| ACY3 | PDCD1 | VIPR1 |
| ADAM11 | GSN | CYP39A1 |
| ADAMTS13 | HS3ST4 | VPS37D |
| ADAMTSL2 | MYH7B | LIPG |
| ADAMTSL3 | KRT71 | FHOD3 |
| ADAMTSL4 | VSIG2 | GATM |
| ADAP1 | EFNB3 | ENHO |
| ADCY1 | AFAP1L2 | TSPAN9 |
| ADCY10 | COL26A1 | MAOB |
| ADCYAP1R1 | TNNT1 | TMEM125 |
| ADH1A | PLA2G2A | LRRN4 |
| ADH1B | SOD3 | TP53INP1 |
| ADH4 | DPP6 | RTN4RL2 |
| ADH6 | BGN | CLDN3 |
| ADH7 | SPRR1A | QSOX1 |
| ADI1 | KRT74 | TCEA3 |
| ADIRF | SLC9A4 | PRSS8 |
| ADM | LINGO1 | CHST7 |
| ADRA1A | ZNF608 | FAM169A |
| ADRA1B | GALNT14 | CD302 |
| ADRA2B | MATN3 | STEAP3 |
| ADRB1 | MFGE8 | SFRP5 |
| ADRB2 | UPK2 | TXNIP |
| ADTRP | FILIP1L | CYP27A1 |
| AFF3 | DKK4 | SH2D4A |
| AFM | CLEC11A | CABLES1 |
| AGBL2 | SLC35F2 | SLC2A12 |
| AGL | CRIP2 | PCSK9 |
| AGMAT | FN1 | CAPN5 |
| AGMO | MAGEA8 | CYP2S1 |
| AGPAT2 | PRRT4 | |
| AGTR1 | YPEL1 | |
| AGXT | DACT1 | |
| AGXT2 | PYGL | |
| AHSG | NGFR | |
| AIG1 | LYZ | |
| AJAP1 | ASTN1 | |
| AKAP3 | FRMPD1 | |
| AKR1A1 | RGL1 | |
| AKR1C4 | SMIM24 | |
| AKR1D1 | MGAT3 | |
| AL078585.1 | MAL2 | |
| AL590714.1 | SCUBE2 | |
| ALAD | ALOX5 | |
| ALAS1 | EMID1 | |
| ALAS2 | FLRT3 | |
| ALB | TBC1D4 | |
| ALDH1L1 | VSTM2L | |
| ALDH2 | TUBB2B | |
| ALDH5A1 | MDFI | |
| ALDH6A1 | ZDHHC2 | |
| ALDH7A1 | TRDC | |
| ALDH9A1 | KRT13 | |
| ALDOC | NLRP7 | |
| ALLC | FZD10 | |
| ALPL | SOBP | |
| AMACR | SEMA6B | |

| | |
|---|---|
| AMBP | C1orf64 |
| AMHR2 | TUBA1A |
| AMN | MMP2 |
| AMOTL2 | IGSF9 |
| AMT | PNMA2 |
| ANG | XG |
| ANGPTL6 | EBF4 |
| ANGPTL7 | TMEM132E |
| ANK3 | SLC2A10 |
| ANKRD20A3 | C1QTNF5 |
| ANKRD55 | AC120057.3 |
| ANO1 | CD8B |
| ANPEP | CERCAM |
| ANXA10 | PLAU |
| ANXA3 | CDH3 |
| ANXA9 | TM4SF18 |
| AOC1 | KRT75 |
| AP1M2 | ARSJ |
| APBA1 | NRGN |
| APH1A | TDGF1 |
| APOA1 | ADGRF1 |
| APOA2 | PAM |
| APOA5 | ANOS1 |
| APOBEC3A | HAPLN3 |
| APOC1 | CCNJL |
| APOC3 | CDH22 |
| APOF | CTSH |
| APOH | GPR155 |
| APOL1 | FBN2 |
| APOL6 | CEACAM6 |
| APOM | ABHD12B |
| AQP11 | ITGA10 |
| AQP7 | MAP7D2 |
| AQP9 | TRABD2B |
| AR | HPSE |
| ARC | LY6G6C |
| ARHGAP20 | CERS6 |
| ARHGEF10L | DLX2 |
| ARHGEF26 | MGAM2 |
| ARHGEF38 | EPHA1 |
| ARHGEF40 | JPH2 |
| ARID3C | KIRREL |
| ARID5A | ANGPTL2 |
| ARL4D | ITGA3 |
| ARMC3 | COMP |
| ARMC5 | SCD5 |
| ARRDC3 | COL13A1 |
| ARSF | COL5A1 |
| ART3 | PCDHA6 |
| ART4 | HABP2 |
| ART5 | FAM46C |
| ARVCF | LY6E |
| ASB4 | ADGRF4 |
| ASB9 | SPOCK1 |
| ASCL2 | DIO1 |
| ASGR1 | EFEMP2 |
| ASMTL | COCH |
| ASPA | CIB2 |

| | |
|---|---|
| ASPDH | CACNA1S |
| ASTL | KRT20 |
| ASTN1 | WLS |
| ASXL3 | KLK5 |
| ATAD3C | GPR173 |
| ATF3 | TNS4 |
| ATF5 | SPTSSB |
| ATF7IP2 | PTPRN2 |
| ATHL1 | FAIM2 |
| ATOH8 | SYT11 |
| ATP11C | SEMA5B |
| ATP13A4 | TMEM178B |
| ATP7B | SCRN1 |
| ATRNL1 | AMOT |
| AUTS2 | CRABP2 |
| AVPR1A | SSTR2 |
| AXL | C5 |
| AZGP1 | ANK1 |
| AZU1 | AQP3 |
| B3GAT1 | SLC34A2 |
| B3GAT2 | NT5E |
| B3GNT8 | LIPC |
| B4GALNT3 | FXYD5 |
| BAALC | TMEM173 |
| BACH2 | RASGRF1 |
| BAI3 | CUBN |
| BAIAP2 | BMF |
| BAIAP3 | P2RX4 |
| BBOX1 | FUT1 |
| BCHE | IL1RN |
| BCL2A1 | C1QL1 |
| BCL6 | STXBP6 |
| BCO2 | SALL2 |
| BDH1 | KCNA2 |
| BEND4 | GLB1L2 |
| BEND7 | KCNH3 |
| BEX1 | SLC4A11 |
| BEX4 | NTSR1 |
| BHMT | MMP7 |
| BHMT2 | EHD3 |
| BLK | NREP |
| BLNK | SARDH |
| BMP10 | NACAD |
| BMP3 | FLI1 |
| BMP5 | THSD1 |
| BMPER | MGLL |
| BNC1 | CA11 |
| BOK | CYP1B1 |
| BPHL | CEACAM5 |
| BPI | VWA5B2 |
| BRINP2 | LRRC2 |
| BRIP1 | CSRNP3 |
| BTD | GPR161 |
| BTNL8 | CLDN5 |
| BZRAP1 | PMAIP1 |
| C10orf11 | REEP2 |
| C10orf25 | SULF2 |
| C10orf67 | STK32B |

| | |
|---|---|
| C11orf31 | SEMA3C |
| C11orf35 | PARM1 |
| C11orf54 | PHGDH |
| C11orf65 | SLCO3A1 |
| C11orf71 | CACNA2D2 |
| C11orf85 | ARHGAP8 |
| C12orf42 | B3GALNT1 |
| C12orf68 | LAMC2 |
| C12orf74 | C14orf105 |
| C14orf164 | F2R |
| C14orf180 | EPHB3 |
| C14orf80 | KRT5 |
| C15orf26 | FIBCD1 |
| C15orf43 | CGNL1 |
| C16orf96 | RASSF4 |
| C19orf12 | ATP2C2 |
| C19orf38 | PDGFB |
| C19orf66 | ADGRF5 |
| C19orf71 | FER1L6 |
| C19orf80 | TRIB2 |
| C1orf115 | KCNIP3 |
| C1orf116 | CLGN |
| C1orf162 | OTC |
| C1orf168 | FMOD |
| C1orf172 | COL6A3 |
| C1orf173 | CPVL |
| C1orf21 | PLTP |
| C1orf210 | MERTK |
| C1orf226 | CYBRD1 |
| C1orf53 | HSPG2 |
| C1QB | AADAC |
| C1QC | SBK1 |
| C1R | DKK1 |
| C1RL | SLC16A2 |
| C1S | MT1X |
| C2 | PTPRE |
| C21orf2 | EPHA7 |
| C21orf33 | TRPM8 |
| C21orf37 | KLHL29 |
| C21orf62 | APOH |
| C21orf67 | AMIGO2 |
| C21orf88 | ARNT2 |
| C21orf90 | C1orf115 |
| C21orf91 | EGR1 |
| C2CD4B | TIMP2 |
| C2orf40 | PROM2 |
| C2orf54 | ZNF423 |
| C2orf72 | SV2A |
| C4BPA | FOXJ1 |
| C5AR1 | KRT6B |
| C5orf27 | KCNK9 |
| C5orf49 | KIAA1324L |
| C6 | COL17A1 |
| C7orf10 | SLC29A1 |
| C8A | SERPINE2 |
| C8B | ELFN2 |
| C8orf82 | ZNF618 |
| C9 | PCDHB5 |

| | |
|---|---|
| C9orf43 | MEGF6 |
| C9orf72 | OSBPL6 |
| C9orf96 | SPNS2 |
| CA1 | PTN |
| CA14 | MRC2 |
| CABLES1 | PIK3AP1 |
| CABP4 | TNXB |
| CACNA2D2 | DHRS9 |
| CADM2 | UBASH3B |
| CALN1 | TWSG1 |
| CAMK2B | SYNPR |
| CAMK2N1 | CAPN6 |
| CAMK4 | SRRM3 |
| CAND2 | HSPA2 |
| CAPN12 | TMEM98 |
| CAPN5 | SLC14A1 |
| CAPZA3 | MEIS3 |
| CARHSP1 | VSIG10L |
| CASC10 | ASAP3 |
| CASR | PDP1 |
| CAT | ADAMTS9 |
| CBFA2T3 | SPAG1 |
| CBLC | ARMCX2 |
| CBLN4 | LCK |
| CBR4 | CLDN10 |
| CBS | EPS8L1 |
| CBX7 | APOBEC3B |
| CCBE1 | LIMD2 |
| CCDC150 | SEMA4F |
| CCDC151 | DUSP6 |
| CCDC152 | CHN2 |
| CCDC158 | APBB1 |
| CCDC178 | SYNE3 |
| CCDC64B | LEF1 |
| CCDC68 | MMP11 |
| CCDC73 | GPRC5B |
| CCL14 | PKN1 |
| CCL16 | EEPD1 |
| CCL2 | HEPHL1 |
| CCL23 | SHROOM3 |
| CCL24 | SNED1 |
| CCL28 | DNAH2 |
| CCL3 | CCDC80 |
| CCL4L1 | KIAA1211L |
| CCNB1IP1 | SCARF2 |
| CCR1 | ECM1 |
| CCR9 | CHST3 |
| CCT6B | SRD5A2 |
| CD14 | UPK3B |
| CD160 | FRAS1 |
| CD163 | MYL9 |
| CD180 | C15orf52 |
| CD1C | CMTM3 |
| CD1D | CALHM2 |
| CD1E | DOCK11 |
| CD244 | VIPR1 |
| CD274 | DNAJC18 |
| CD300A | CLIP3 |

| | |
|---|---|
| CD300C | RCOR2 |
| CD300E | FBLN1 |
| CD300LB | ARFGEF3 |
| CD300LG | TM4SF4 |
| CD302 | B3GNT7 |
| CD36 | RAC2 |
| CD4 | HES6 |
| CD5L | MRAP2 |
| CD81 | EPB41L2 |
| CD82 | CYP39A1 |
| CD83 | LRRC75A |
| CD9 | OTOG |
| CDH19 | VPS37D |
| CDH2 | AGT |
| CDH23 | GAS6 |
| CDH4 | TNC |
| CDHR5 | LIPG |
| CDK18 | IL15RA |
| CDK3 | FHOD3 |
| CEACAM3 | CACNB3 |
| CEACAM4 | PCDHB15 |
| CEACAM6 | FMR1NB |
| CEBPA | SBSPON |
| CEBPD | OLFML2A |
| CECR2 | GATM |
| CENPV | ZDHHC14 |
| CERS4 | PHC1 |
| CES1 | TGFB1I1 |
| CES2 | SLC41A2 |
| CES3 | PCDHB8 |
| CES4A | DCBLD2 |
| CES5A | PTGS1 |
| CETP | THBS3 |
| CFB | ARAP3 |
| CFD | PCNX2 |
| CFHR1 | ENHO |
| CFHR2 | PCDHB16 |
| CFHR3 | KLK10 |
| CFHR4 | MEX3A |
| CFHR5 | PIK3IP1 |
| CFL2 | AHRR |
| CFP | TSPAN9 |
| CFTR | KREMEN2 |
| CGN | HOXC5 |
| CGNL1 | PPP1R1B |
| CHAD | CPNE2 |
| CHADL | NOS3 |
| CHDC2 | RASSF3 |
| CHDH | MAOB |
| CHN2 | ZNF22 |
| CHRDL1 | SETBP1 |
| CHRM2 | TMEM125 |
| CHRNA4 | F5 |
| CHST13 | FAM174B |
| CHST4 | PLXNC1 |
| CHST7 | PROCR |
| CIDEB | LRRN4 |
| CILP | PDLIM4 |

| | |
|---|---|
| CISH | TP53INP1 |
| CLDN1 | NKD2 |
| CLDN10 | RTN4RL2 |
| CLDN14 | PLCB1 |
| CLDN19 | NEURL1B |
| CLDN3 | ENTPD3 |
| CLDN4 | PAQR8 |
| CLEC1B | DPYSL2 |
| CLEC3B | CLDN3 |
| CLEC4C | QSOX1 |
| CLEC4D | DISC1 |
| CLEC4E | FLNC |
| CLEC4G | NIN |
| CLEC4M | TCEA3 |
| CLVS2 | C16orf45 |
| CMA1 | PRSS8 |
| CMBL | GNAL |
| CMPK2 | DNAAF3 |
| CMTM2 | CHST7 |
| CMTM8 | AKNA |
| CMYA5 | FAM169A |
| CNBD1 | GPD1L |
| CNDP1 | MYO1D |
| CNGA1 | SLC9A2 |
| CNKSR2 | CD302 |
| CNPY3 | LARGE2 |
| CNST | MGAT5 |
| CNTLN | TGFBI |
| CNTN5 | VASH1 |
| COBLL1 | STEAP3 |
| COL11A2 | MAPK8IP1 |
| COL18A1 | LIPH |
| COL25A1 | MMD |
| COL27A1 | PPL |
| COL28A1 | SLC2A6 |
| COL6A6 | GJB4 |
| COLEC10 | SFRP5 |
| COLEC11 | SMPD1 |
| COMT | EPPK1 |
| COQ10A | EMP1 |
| COX6A2 | TXNIP |
| CP | TPBGL |
| CPA3 | MAP3K8 |
| CPAMD8 | PCDHGC3 |
| CPB2 | CYTH3 |
| CPEB3 | PDLIM1 |
| CPED1 | CYP27A1 |
| CPN1 | IMPDH1 |
| CPN2 | MACC1 |
| CPNE6 | EVC |
| CR1 | SLC2A3 |
| CR1L | TPPP |
| CRHBP | PKDCC |
| CRLS1 | SH2D4A |
| CRYAA | CABLES1 |
| CRYL1 | SLC2A12 |
| CSAD | MCUB |
| CSF3 | CD59 |

| | |
|---|---|
| CSF3R | AC241585.3 |
| CSRNP1 | STBD1 |
| CTD-2014B16.3 | PCSK9 |
| CTF1 | CAPN5 |
| CTNNA3 | N4BP2 |
| CTSG | NUP210 |
| CTSL | SORT1 |
| CTTNBP2 | RAB36 |
| CUX2 | CTHRC1 |
| CX3CL1 | ARL4C |
| CXADR | MUC1 |
| CXCL1 | CTSV |
| CXCL12 | ZSWIM5 |
| CXCL14 | PTPN14 |
| CXCL2 | PPM1H |
| CXCL3 | AP3M2 |
| CXCR1 | TTC39B |
| CXCR2 | PXK |
| CXorf22 | LARP6 |
| CXorf66 | ABTB1 |
| CYB5A | CYP2S1 |
| CYP11A1 | GDPD5 |
| CYP1A1 | ORAI2 |
| CYP1A2 | MLXIPL |
| CYP26A1 | EPHA10 |
| CYP26B1 | KREMEN1 |
| CYP27A1 | LIMA1 |
| CYP27C1 | TPM1 |
| CYP2A13 | CHRNA5 |
| CYP2A6 | PCDHB2 |
| CYP2A7 | PTPRG |
| CYP2B6 | HES1 |
| CYP2C19 | MYO6 |
| CYP2C8 | |
| CYP2C9 | |
| CYP2D6 | |
| CYP2J2 | |
| CYP2S1 | |
| CYP39A1 | |
| CYP3A4 | |
| CYP3A43 | |
| CYP3A5 | |
| CYP3A7 | |
| CYP4A11 | |
| CYP4A22 | |
| CYP4F12 | |
| CYP4F2 | |
| CYP4F3 | |
| CYP4V2 | |
| CYP4X1 | |
| CYP4Z1 | |
| CYP7A1 | |
| CYP7B1 | |
| CYP8B1 | |
| DAB1 | |
| DACH1 | |
| DAK | |
| DAO | |

| |
|---|
| DBH |
| DBI |
| DCAF11 |
| DCHS2 |
| DCT |
| DCXR |
| DDC |
| DDT |
| DDTL |
| DDX25 |
| DEFA3 |
| DEPDC7 |
| DGAT2 |
| DGCR6L |
| DHCR24 |
| DHCR7 |
| DHFR |
| DHODH |
| DHRS1 |
| DHRS12 |
| DHRS2 |
| DHRS4 |
| DHRS4L2 |
| DHTKD1 |
| DIO1 |
| DIRAS3 |
| DLEU7 |
| DLGAP2 |
| DMD |
| DMGDH |
| DMRTA1 |
| DNAH6 |
| DNAJC12 |
| DNAJC19 |
| DNALI1 |
| DNASE1L3 |
| DNMT3L |
| DOK2 |
| DOK6 |
| DPEP2 |
| DPEP3 |
| DPF3 |
| DPP4 |
| DPPA4 |
| DPT |
| DPYD |
| DPYS |
| DRD1 |
| DSC2 |
| DSCAM |
| DSG1 |
| DSG4 |
| DTX1 |
| DUSP10 |
| DUSP14 |
| DYNLRB2 |
| EBF4 |
| EBI3 |

| |
|---|
| EBP |
| EBPL |
| ECH1 |
| ECHDC2 |
| ECHDC3 |
| ECI2 |
| ECM1 |
| EDAR |
| EDN1 |
| EDNRB |
| EFCAB1 |
| EFCC1 |
| EFHD1 |
| EFNA2 |
| EHD3 |
| EHHADH |
| EIF1AY |
| ELAC1 |
| ELAVL4 |
| ELF5 |
| ELFN1 |
| ELOVL2 |
| ELOVL6 |
| EMILIN3 |
| EMP2 |
| EMR1 |
| EMR3 |
| ENDOU |
| ENHO |
| ENO3 |
| ENPEP |
| ENPP3 |
| ENPP6 |
| ENPP7 |
| ENTPD5 |
| ENTPD8 |
| EPB41L4B |
| EPHA1 |
| EPHA7 |
| EPHX1 |
| EPHX2 |
| EPO |
| EPOR |
| EPS8L2 |
| ERF |
| ERVFRD-1 |
| ESPN |
| ESPNL |
| ESR1 |
| ESRP1 |
| ETFDH |
| ETNK2 |
| EVA1A |
| EVPLL |
| EXOC3L4 |
| EXPH5 |
| EYA4 |
| F10 |

| |
|---|
| F11 |
| F12 |
| F13B |
| F2 |
| F7 |
| F8 |
| F9 |
| FAAH |
| FABP1 |
| FABP3 |
| FADS2 |
| FADS6 |
| FAH |
| FAM107A |
| FAM110C |
| FAM124A |
| FAM129C |
| FAM134B |
| FAM135B |
| FAM13A |
| FAM149A |
| FAM150B |
| FAM151A |
| FAM163B |
| FAM169A |
| FAM180A |
| FAM198A |
| FAM211B |
| FAM213A |
| FAM227B |
| FAM229B |
| FAM26F |
| FAM3B |
| FAM46C |
| FAM65C |
| FAM69B |
| FAM71D |
| FAM83B |
| FAM83E |
| FAM9B |
| FANCC |
| FAT3 |
| FAXDC2 |
| FBN3 |
| FBP1 |
| FBXO15 |
| FBXO2 |
| FBXO40 |
| FBXO6 |
| FBXO7 |
| FCAMR |
| FCAR |
| FCER1A |
| FCER2 |
| FCGR2B |
| FCGR3A |
| FCGR3B |
| FCGRT |

| |
|---|
| FCN1 |
| FCN2 |
| FCN3 |
| FCRL1 |
| FCRLB |
| FDPS |
| FERMT2 |
| FETUB |
| FEZ1 |
| FFAR2 |
| FGD4 |
| FGF10 |
| FGF21 |
| FGF23 |
| FGFBP2 |
| FGFR2 |
| FGFR3 |
| FGFR4 |
| FGL2 |
| FGR |
| FHL1 |
| FHOD3 |
| FITM1 |
| FLRT1 |
| FMO3 |
| FMO4 |
| FMO5 |
| FNDC5 |
| FNIP2 |
| FOLH1 |
| FOS |
| FOSB |
| FOXA2 |
| FOXA3 |
| FOXD3 |
| FOXN2 |
| FOXN3 |
| FOXO1 |
| FPR1 |
| FPR2 |
| FRAS1 |
| FRAT1 |
| FREM1 |
| FREM2 |
| FRMD1 |
| FRMD4B |
| FRMD7 |
| FRMPD4 |
| FRRS1 |
| FTCD |
| FTCDNL1 |
| FUOM |
| FUT3 |
| FUT6 |
| FUZ |
| FXN |
| FXYD1 |
| FXYD3 |

| |
|---|
| FXYD7 |
| FZD9 |
| G0S2 |
| GABRA2 |
| GABRG1 |
| GABRP |
| GADD45A |
| GADD45G |
| GAL3ST2 |
| GALK1 |
| GALM |
| GALNT14 |
| GALNT3 |
| GALT |
| GAMT |
| GAS2 |
| GATA4 |
| GATA5 |
| GATM |
| GBP1 |
| GBP7 |
| GCAT |
| GCDH |
| GCGR |
| GCHFR |
| GCK |
| GCKR |
| GCOM1 |
| GCSAML |
| GDF2 |
| GDPD4 |
| GFI1B |
| GFRA1 |
| GFRA2 |
| GFRA3 |
| GGACT |
| GGT6 |
| GHR |
| GHRL |
| GIPC2 |
| GJB2 |
| GJB3 |
| GJC3 |
| GLIPR1L2 |
| GLOD5 |
| GLP2R |
| GLUD1 |
| GLUL |
| GLYAT |
| GLYATL1 |
| GLYATL3 |
| GLYCTK |
| GMPR |
| GNA14 |
| GNAO1 |
| GNAT1 |
| GNE |
| GNG5P2 |

| |
|---|
| GNMT |
| GNPNAT1 |
| GNRH2 |
| GOLGA6A |
| GOLGA6B |
| GOLGA8M |
| GP1BA |
| GPAM |
| GPC6 |
| GPD1 |
| GPER1 |
| GPHN |
| GPLD1 |
| GPM6A |
| GPM6B |
| GPR123 |
| GPR125 |
| GPR126 |
| GPR128 |
| GPR142 |
| GPR143 |
| GPR158 |
| GPR162 |
| GPR182 |
| GPR75 |
| GPR82 |
| GPR83 |
| GPR88 |
| GPR98 |
| GPRIN2 |
| GRAMD1C |
| GRAP |
| GREB1 |
| GREB1L |
| GRHL1 |
| GRHL2 |
| GRHL3 |
| GRHPR |
| GRIK3 |
| GRM8 |
| GRTP1 |
| GSDMB |
| GSTA1 |
| GSTA2 |
| GSTM5 |
| GSTT2 |
| GSTT2B |
| GSTZ1 |
| GYLTL1B |
| GYS2 |
| GZMM |
| H1FX |
| HAAO |
| HABP2 |
| HAGH |
| HAMP |
| HAO1 |
| HAO2 |

| |
|---|
| HAP1 |
| HAS2 |
| HAUS4 |
| HBA1 |
| HBA2 |
| HBB |
| HBCBP |
| HBD |
| HBG2 |
| HCAR2 |
| HCAR3 |
| HDAC6 |
| HDC |
| HEMGN |
| HEPACAM |
| HERC5 |
| HEY2 |
| HFE2 |
| HGF |
| HGFAC |
| HHIP |
| HK3 |
| HLF |
| HLX |
| HMGCL |
| HMGCLL1 |
| HMGCR |
| HMGCS1 |
| HMGCS2 |
| HMOX1 |
| HNMT |
| HOGA1 |
| HOMER2 |
| HORMAD2 |
| HP |
| HPGDS |
| HPN |
| HPR |
| HPRT1 |
| HPX |
| HRASLS2 |
| HRG |
| HS3ST3A1 |
| HS3ST3B1 |
| HS3ST4 |
| HSBP1L1 |
| HSD11B1 |
| HSD17B11 |
| HSD17B13 |
| HSD17B14 |
| HSD17B6 |
| HSD17B8 |
| HSD3B1 |
| HSD3B2 |
| HSDL2 |
| HSPB9 |
| HYDIN |
| ICAM4 |

| |
|---|
| ID2 |
| IDNK |
| IDO2 |
| IFI27 |
| IFI44L |
| IFIT1 |
| IFIT1B |
| IFIT2 |
| IFIT3 |
| IFITM10 |
| IFNLR1 |
| IFT46 |
| IGF2 |
| IGFALS |
| IGFBP1 |
| IGFBP2 |
| IGFBP3 |
| IGJ |
| IGLON5 |
| IGSF10 |
| IGSF23 |
| IGSF9 |
| IHH |
| IL10 |
| IL11RA |
| IL13RA2 |
| IL17RB |
| IL17RC |
| IL17RE |
| IL1B |
| IL1RAP |
| IL1RN |
| IL20RA |
| IL22RA1 |
| IL23A |
| IL27 |
| IL33 |
| IMPA2 |
| INHBC |
| INMT |
| INS-IGF2 |
| INSIG2 |
| IQGAP2 |
| IQSEC3 |
| IRF6 |
| IRF7 |
| ISPD |
| ITCH |
| ITGA2B |
| ITGA9 |
| ITGAD |
| ITIH1 |
| ITIH2 |
| ITIH3 |
| ITLN1 |
| ITPR2 |
| IYD |
| JAKMIP2 |

| |
|---|
| JDP2 |
| JUN |
| JUNB |
| JUND |
| KANK4 |
| KAZN |
| KBTBD11 |
| KCNAB1 |
| KCND3 |
| KCNE1 |
| KCNH7 |
| KCNH8 |
| KCNJ10 |
| KCNJ13 |
| KCNJ15 |
| KCNJ3 |
| KCNK1 |
| KCNK17 |
| KCNK3 |
| KCNK5 |
| KCNMB2 |
| KCNN2 |
| KCTD16 |
| KDM8 |
| KHK |
| KIAA1161 |
| KIF12 |
| KIF17 |
| KIF19 |
| KIF1A |
| KIF25 |
| KIF26A |
| KIF6 |
| KIRREL3 |
| KLB |
| KLC4 |
| KLF10 |
| KLF11 |
| KLF12 |
| KLF4 |
| KLHL13 |
| KLHL33 |
| KLHL4 |
| KLKB1 |
| KLLN |
| KLRF1 |
| KNG1 |
| KRBOX1 |
| KRT1 |
| KRT19 |
| KRT73 |
| KRTAP5-9 |
| KRTCAP3 |
| L1CAM |
| LAG3 |
| LARGE |
| LBX2 |
| LCAT |

| |
|---|
| LDLR |
| LDLRAD4 |
| LEAP2 |
| LEPREL1 |
| LGALS12 |
| LGALS4 |
| LGI1 |
| LGI4 |
| LGSN |
| LHX2 |
| LIFR |
| LILRA1 |
| LILRA2 |
| LILRA5 |
| LILRA6 |
| LILRB1 |
| LILRB2 |
| LILRB3 |
| LILRB5 |
| LIMD2 |
| LIME1 |
| LINC00923 |
| LINGO4 |
| LIPA |
| LIPC |
| LIPG |
| LIPJ |
| LIPN |
| LNP1 |
| LONRF3 |
| LPA |
| LPIN2 |
| LRAT |
| LRFN5 |
| LRIG3 |
| LRP1B |
| LRP3 |
| LRRC16B |
| LRRC19 |
| LRRC25 |
| LRRC3 |
| LRRC31 |
| LRRC3DN |
| LRRC4 |
| LRRC55 |
| LRRFIP2 |
| LRRIQ1 |
| LRRK2 |
| LRRN1 |
| LRRN3 |
| LRRN4 |
| LRRTM1 |
| LRRTM2 |
| LRRTM4 |
| LSR |
| LSS |
| LST1 |
| LST3 |

| |
|---|
| LTK |
| LY6E |
| LYNX1 |
| LYVE1 |
| LZTFL1 |
| MAATS1 |
| MACROD1 |
| MAFF |
| MAG |
| MAGEH1 |
| MAK |
| MAN1C1 |
| MAOA |
| MAOB |
| MAP1LC3A |
| MAP3K13 |
| MAPK4 |
| MARCO |
| MARVELD3 |
| MASP1 |
| MASP2 |
| MAT1A |
| MATN2 |
| MBL2 |
| MBNL3 |
| MBOAT1 |
| MCHR1 |
| MDGA2 |
| MEFV |
| MEGF10 |
| MEI4 |
| MEIOB |
| MEP1B |
| MEST |
| METTL20 |
| METTL7A |
| MFAP3L |
| MFI2 |
| MFSD2A |
| MFSD4 |
| MGAM |
| MGMT |
| MGST2 |
| MLANA |
| MLF1 |
| MLIP |
| MLK4 |
| MMAB |
| MME |
| MMP19 |
| MMP25 |
| MMP7 |
| MMP8 |
| MMRN1 |
| MOGAT2 |
| MOGAT3 |
| MORC1 |
| MORC3 |

| |
|---|
| MPDZ |
| MPPED1 |
| MPST |
| MRC1L1 |
| MREG |
| MRGPRF |
| MRO |
| MROH2A |
| MROH2B |
| MROH7 |
| MROH8 |
| MRPL23 |
| MRPS6 |
| MS4A2 |
| MS4A6A |
| MSMO1 |
| MST1 |
| MT1G |
| MT1H |
| MT1M |
| MTHFD1 |
| MTHFS |
| MTTP |
| MTUS2 |
| MUSK |
| MUT |
| MVK |
| MYCL |
| MYCT1 |
| MYH7B |
| MYL3 |
| MYO15A |
| MYO16 |
| MYO1B |
| MYO1F |
| MYO3A |
| MYRIP |
| MYT1L |
| N4BP2L1 |
| NAAA |
| NAALAD2 |
| NAB2 |
| NAGS |
| NANOS1 |
| NAT1 |
| NAT2 |
| NAT8 |
| NCAM1 |
| NCF1 |
| NCKAP5 |
| NCMAP |
| NCR1 |
| NDNF |
| NDRG2 |
| NDST3 |
| NECAB2 |
| NEIL1 |
| NEU4 |

| |
|---|
| NFAM1 |
| NFATC2 |
| NFE2 |
| NFIA |
| NFKBIA |
| NFKBIZ |
| NGF |
| NGFR |
| NGFRAP1 |
| NHSL1 |
| NIPAL1 |
| NIPSNAP1 |
| NLRC4 |
| NLRP11 |
| NLRP12 |
| NLRP14 |
| NLRP3 |
| NLRP6 |
| NMRK1 |
| NOL4 |
| NOS1 |
| NOS1AP |
| NOTUM |
| NOXO1 |
| NPAS3 |
| NPC1L1 |
| NPR1 |
| NPR2 |
| NR0B2 |
| NR1H3 |
| NR1I2 |
| NR1I3 |
| NR2F6 |
| NR5A2 |
| NRAP |
| NREP |
| NRG1 |
| NRG3 |
| NRG4 |
| NRN1 |
| NRXN1 |
| NSUN6 |
| NSUN7 |
| NT5DC3 |
| NT5E |
| NTF3 |
| NTHL1 |
| NTN1 |
| NTN3 |
| NTN4 |
| NTRK1 |
| NUDT10 |
| NUDT4 |
| NUDT6 |
| NUGGC |
| NUP62CL |
| NXF3 |
| NYAP1 |

| |
|---|
| OAF |
| OAS1 |
| OCLN |
| ODF3L1 |
| OGDHL |
| OIT3 |
| OLFM1 |
| OLFM4 |
| OLFML3 |
| OPRK1 |
| OR10J5 |
| OR13C3 |
| OR13C4 |
| OR13C5 |
| OR13C9 |
| OR2W3 |
| ORMDL3 |
| OSBPL6 |
| OSGIN1 |
| OTC |
| OTOA |
| OXER1 |
| OXT |
| P2RX3 |
| P2RX4 |
| P2RX6 |
| P2RY12 |
| P2RY13 |
| P2RY2 |
| P4HA1 |
| PACRG |
| PACSIN1 |
| PACSIN3 |
| PADI4 |
| PAGE5 |
| PAK7 |
| PALM3 |
| PAMR1 |
| PANK1 |
| PANX2 |
| PAQR7 |
| PAQR9 |
| PATZ1 |
| PBLD |
| PC |
| PCDH15 |
| PCDH20 |
| PCDH9 |
| PCDHAC1 |
| PCDHAC2 |
| PCK2 |
| PCLO |
| PCOLCE2 |
| PCP2 |
| PCP4L1 |
| PCSK2 |
| PCSK9 |
| PCYT2 |

| |
|---|
| PDCD1LG2 |
| PDE11A |
| PDE2A |
| PDE6G |
| PDE8B |
| PDIA5 |
| PDK2 |
| PDLIM2 |
| PDXP |
| PDZRN4 |
| PEBP1 |
| PEBP4 |
| PECR |
| PEG10 |
| PEG3 |
| PEMT |
| PER2 |
| PER3 |
| PEX11G |
| PFKFB1 |
| PGAM2 |
| PGAP3 |
| PGLYRP2 |
| PGM1 |
| PGM2 |
| PGM5 |
| PGRMC1 |
| PHACTR3 |
| PHGDH |
| PHLDA1 |
| PHLPP1 |
| PHOSPHO1 |
| PHYH |
| PHYHD1 |
| PHYHIPL |
| PI16 |
| PID1 |
| PIGR |
| PIK3AP1 |
| PIK3C2G |
| PIK3R1 |
| PIM1 |
| PIPOX |
| PITPNM3 |
| PKD2L1 |
| PKHD1L1 |
| PKLR |
| PKP2 |
| PLA1A |
| PLA2G12B |
| PLA2G16 |
| PLAC8 |
| PLCL2 |
| PLCXD2 |
| PLD5 |
| PLEK2 |
| PLEKHA4 |
| PLEKHA6 |

| |
|---|
| PLEKHB1 |
| PLEKHF1 |
| PLEKHG6 |
| PLEKHG7 |
| PLG |
| PLGLB1 |
| PLGLB2 |
| PLIN1 |
| PLIN2 |
| PLK3 |
| PLP1 |
| PLSCR4 |
| PM20D1 |
| PMEL |
| PMM1 |
| PMP2 |
| PNMA3 |
| PNMA6C |
| PNMAL2 |
| PNPLA3 |
| PNPLA7 |
| POFUT1 |
| POLR3GL |
| POMC |
| PON3 |
| POU6F2 |
| PPAP2B |
| PPARA |
| PPBP |
| PPFIBP2 |
| PPP1R15A |
| PPP1R1A |
| PPP1R3B |
| PPP1R3C |
| PPP4R4 |
| PPP6R2 |
| PQLC1 |
| PRAM1 |
| PRAP1 |
| PRELID2 |
| PRG4 |
| PRH2 |
| PRIMA1 |
| PRKAR2B |
| PRKCB |
| PRLR |
| PROC |
| PRODH2 |
| PROK2 |
| PROM1 |
| PROSER2 |
| PROZ |
| PRPSAP1 |
| PRR18 |
| PRR22 |
| PRR5 |
| PRRG4 |
| PRSS12 |

| |
|---|
| PRSS22 |
| PRSS36 |
| PRSS42 |
| PRSS45 |
| PRSS50 |
| PRSS53 |
| PRSS8 |
| PSAT1 |
| PTCHD3 |
| PTCRA |
| PTGDR2 |
| PTGR1 |
| PTGS2 |
| PTH1R |
| PTK6 |
| PTMS |
| PTN |
| PTPRB |
| PTPRD |
| PTPRN2 |
| PTPRS |
| PTPRT |
| PVALB |
| PVRL3 |
| PXMP2 |
| PYGL |
| PYROXD2 |
| PZP |
| QPRT |
| QRICH2 |
| QSOX1 |
| RAB17 |
| RAB25 |
| RAB26 |
| RAB27B |
| RAB39A |
| RAB3IL1 |
| RAD51AP2 |
| RAD54L2 |
| RAD9B |
| RAG1 |
| RALGPS2 |
| RALYL |
| RANBP3L |
| RARRES2 |
| RARRES3 |
| RASGEF1B |
| RASGRP2 |
| RASGRP4 |
| RASL10A |
| RASL10B |
| RASL11A |
| RASSF5 |
| RBP5 |
| RCAN1 |
| RCL1 |
| RD3L |
| RDH16 |

| |
|---|
| RENBP |
| RET |
| RFNG |
| RFPL1 |
| RGL1 |
| RGN |
| RGPD2 |
| RGS18 |
| RGS3 |
| RGSL1 |
| RHBG |
| RHCE |
| RHOB |
| RIC3 |
| RIPK4 |
| RIPPLY1 |
| RIPPLY3 |
| RMDN2 |
| RMND5A |
| RND2 |
| RNF125 |
| RNF144B |
| RNF152 |
| RNF165 |
| ROBO2 |
| ROPN1L |
| RORC |
| ROS1 |
| RP11-10A14.4 |
| RP11-146D12.2 |
| RP11-181C3.1 |
| RP11-242G20.1 |
| RP11-321F6.1 |
| RP11-422N16.3 |
| RP11-595B24.2 |
| RP11-650K20.3 |
| RP11-676J12.7 |
| RP11-766F14.2 |
| RP11-817J15.3 |
| RP11-867G23.8 |
| RP11-986E7.7 |
| RP11-998D10.1 |
| RPGRIP1 |
| RPS29 |
| RPS4Y1 |
| RPS6KA6 |
| RSAD2 |
| RSPH10B |
| RSPH4A |
| RSPO2 |
| RTN4RL1 |
| RTN4RL2 |
| RTP3 |
| RTP4 |
| RTTN |
| RUNDC3B |
| RXFP1 |
| RXRG |

| |
|---|
| S100A1 |
| S100A12 |
| S100A14 |
| S100A8 |
| S100A9 |
| S1PR5 |
| SAA4 |
| SALL4 |
| SAMD4A |
| SAMD5 |
| SARDH |
| SAT2 |
| SATB1 |
| SC5D |
| SCG5 |
| SCGB3A1 |
| SCIMP |
| SCN11A |
| SCN2A |
| SCN3A |
| SCN7A |
| SCN9A |
| SCNN1B |
| SCNN1D |
| SCP2 |
| SCRN2 |
| SDC3 |
| SDK2 |
| SDPR |
| SEC14L2 |
| SEC14L3 |
| SEC14L4 |
| SELE |
| SELENBP1 |
| SELO |
| SEMA3D |
| SEMA3E |
| SEMA4A |
| SEMA4G |
| SEMA6A |
| SEMA6C |
| SEMA6D |
| SEPP1 |
| SERP2 |
| SERPINA10 |
| SERPINA11 |
| SERPINA4 |
| SERPINA5 |
| SERPINA6 |
| SERPINA7 |
| SERPINC1 |
| SERPIND1 |
| SERPINE1 |
| SERPINF1 |
| SERPINF2 |
| SEZ6L |
| SFRP1 |
| SFRP5 |

| |
|---|
| SFTPD |
| SFXN5 |
| SGCE |
| SGCZ |
| SH2D1B |
| SH2D4A |
| SH2D6 |
| SH3BP2 |
| SH3BP5 |
| SH3GL2 |
| SH3RF2 |
| SHB |
| SHBG |
| SHD |
| SHF |
| SHH |
| SHMT1 |
| SHROOM2 |
| SIGIRR |
| SIGLEC1 |
| SIGLEC11 |
| SIGLEC14 |
| SIGLEC15 |
| SIGLEC7 |
| SIGLEC9 |
| SIM1 |
| SIRPB1 |
| SIRT5 |
| SIVA1 |
| SKAP1 |
| SKIDA1 |
| SKOR1 |
| SLAIN1 |
| SLC10A1 |
| SLC13A5 |
| SLC15A1 |
| SLC16A1 |
| SLC16A10 |
| SLC16A12 |
| SLC16A2 |
| SLC16A4 |
| SLC16A9 |
| SLC17A2 |
| SLC17A8 |
| SLC18A2 |
| SLC19A1 |
| SLC19A2 |
| SLC19A3 |
| SLC22A1 |
| SLC22A10 |
| SLC22A24 |
| SLC22A25 |
| SLC22A7 |
| SLC22A9 |
| SLC23A1 |
| SLC24A2 |
| SLC25A1 |
| SLC25A13 |

| |
|---|
| SLC25A18 |
| SLC25A20 |
| SLC25A21 |
| SLC25A21-AS1 |
| SLC25A27 |
| SLC25A34 |
| SLC25A47 |
| SLC26A1 |
| SLC26A5 |
| SLC27A2 |
| SLC27A3 |
| SLC27A5 |
| SLC28A1 |
| SLC28A2 |
| SLC2A12 |
| SLC2A2 |
| SLC2A4RG |
| SLC2A9 |
| SLC30A1 |
| SLC30A10 |
| SLC31A2 |
| SLC34A1 |
| SLC35D1 |
| SLC37A4 |
| SLC38A11 |
| SLC38A4 |
| SLC39A4 |
| SLC39A5 |
| SLC3A1 |
| SLC43A3 |
| SLC45A3 |
| SLC47A1 |
| SLC4A1 |
| SLC51A |
| SLC52A1 |
| SLC5A1 |
| SLC5A7 |
| SLC5A9 |
| SLC6A1 |
| SLC6A12 |
| SLC6A13 |
| SLC6A19 |
| SLC6A20 |
| SLC6A4 |
| SLC7A8 |
| SLC7A9 |
| SLC8A1 |
| SLC9A3R2 |
| SLC9B2 |
| SLCO1A2 |
| SLCO1B3 |
| SLCO1B7 |
| SLCO2B1 |
| SLCO4C1 |
| SLITRK2 |
| SLITRK3 |
| SLITRK6 |
| SMAD6 |

| |
|---|
| SMCO3 |
| SMIM1 |
| SMIM14 |
| SMIM19 |
| SMIM9 |
| SMLR1 |
| SMO |
| SMPD3 |
| SNCA |
| SNTB1 |
| SNTG1 |
| SOAT2 |
| SOBP |
| SOCS2 |
| SOCS6 |
| SOD1 |
| SORCS1 |
| SORD |
| SORL1 |
| SOX10 |
| SOX5 |
| SPDYC |
| SPECC1L-ADORA2A |
| SPI1 |
| SPIB |
| SPIC |
| SPINT2 |
| SPOCK3 |
| SPP2 |
| SPSB3 |
| SPSB4 |
| SPTBN2 |
| SQLE |
| SRCIN1 |
| SRD5A1 |
| SRPX |
| SSTR1 |
| SSTR2 |
| ST14 |
| ST3GAL1 |
| ST3GAL6 |
| ST6GAL1 |
| ST6GALNAC2 |
| ST6GALNAC3 |
| ST8SIA3 |
| STAB1 |
| STAB2 |
| STAG3 |
| STARD10 |
| STARD4 |
| STEAP3 |
| STEAP4 |
| STMND1 |
| STPG2 |
| SUCNR1 |
| SULT1A1 |
| SULT1A2 |
| SULT1E1 |

| |
|---|
| SULT2A1 |
| SUN2 |
| SUSD4 |
| SYBU |
| SYCE1 |
| SYDE2 |
| SYNE4 |
| SYNGR1 |
| SYT1 |
| SYT10 |
| SYT12 |
| SYT15 |
| SYT17 |
| SYT7 |
| SYTL4 |
| TACSTD2 |
| TADA1 |
| TAS1R3 |
| TBL1Y |
| TBX20 |
| TBXA2R |
| TBXAS1 |
| TCEA3 |
| TCEAL2 |
| TCF21 |
| TCHH |
| TCL1A |
| TCP10L |
| TCP10L2 |
| TCTEX1D1 |
| TCTEX1D4 |
| TDRD10 |
| TDRD6 |
| TEF |
| TEK |
| TEKT2 |
| TEKT5 |
| TENM1 |
| TENM2 |
| TESK2 |
| TEX30 |
| TF |
| TFPI2 |
| TFR2 |
| TGFBR3 |
| THEMIS2 |
| THNSL1 |
| THOP1 |
| THRSP |
| TIAM1 |
| TIMD4 |
| TINAGL1 |
| TIPARP |
| TJP2 |
| TKTL1 |
| TLR4 |
| TM6SF2 |
| TMCO6 |

| |
|---|
| TMEFF2 |
| TMEM105 |
| TMEM121 |
| TMEM125 |
| TMEM132C |
| TMEM132D |
| TMEM139 |
| TMEM150C |
| TMEM170B |
| TMEM176B |
| TMEM200B |
| TMEM200C |
| TMEM220 |
| TMEM232 |
| TMEM25 |
| TMEM252 |
| TMEM26 |
| TMEM27 |
| TMEM30B |
| TMEM37 |
| TMEM45B |
| TMEM47 |
| TMEM52 |
| TMEM56 |
| TMEM63C |
| TMEM71 |
| TMEM82 |
| TMEM86B |
| TMEM97 |
| TMIE |
| TMPO |
| TMPRSS2 |
| TMPRSS4 |
| TMPRSS6 |
| TMPRSS9 |
| TMSB4Y |
| TNF |
| TNFAIP8L1 |
| TNFRSF11B |
| TNFSF10 |
| TNFSF11 |
| TNN |
| TNNC1 |
| TNR |
| TOM1L1 |
| TOX2 |
| TP53I13 |
| TP53INP1 |
| TPH2 |
| TPPP2 |
| TPRG1 |
| TPSAB1 |
| TPST2 |
| TRABD2B |
| TRAPPC3L |
| TRDN |
| TREH |
| TREM1 |

| |
|---|
| TREML2 |
| TRHDE |
| TRIB1 |
| TRIM58 |
| TRIM63 |
| TRPC5 |
| TRPM6 |
| TRPM8 |
| TRPV4 |
| TRPV6 |
| TSHR |
| TSLP |
| TSPAN11 |
| TSPAN12 |
| TSPAN7 |
| TSPAN9 |
| TST |
| TSTD1 |
| TTBK1 |
| TTC36 |
| TTC38 |
| TTC40 |
| TTC7B |
| TTPA |
| TTR |
| TUBB1 |
| TUBE1 |
| TUSC1 |
| TXNDC16 |
| TXNIP |
| TXNRD2 |
| UAP1 |
| UGP2 |
| UGT1A1 |
| UGT2B10 |
| UGT2B15 |
| UGT2B17 |
| UGT2B7 |
| UGT3A1 |
| UNC13D |
| UNC79 |
| UNC93A |
| UPB1 |
| UROC1 |
| USH2A |
| USP18 |
| USP44 |
| USP51 |
| UTY |
| VCAM1 |
| VEPH1 |
| VIL1 |
| VIPR1 |
| VIPR2 |
| VMO1 |
| VNN1 |
| VNN2 |
| VNN3 |

| |
|---|
| VPS37B |
| VPS37D |
| VSIG4 |
| VSNL1 |
| VTCN1 |
| VWA3B |
| VWCE |
| VWDE |
| WBSCR27 |
| WDR17 |
| WDR72 |
| WNK2 |
| WNK3 |
| WNT11 |
| WNT5A |
| WNT5B |
| WNT7A |
| XAGE3 |
| XDH |
| XKR4 |
| XPNPEP2 |
| XRCC6BP1 |
| XYLB |
| YBX2 |
| YPEL1 |
| YPEL2 |
| ZBTB18 |
| ZC3H12C |
| ZCCHC6 |
| ZCWPW1 |
| ZDHHC19 |
| ZFP1 |
| ZFY |
| ZG16 |
| ZIC1 |
| ZMYND12 |
| ZNF175 |
| ZNF268 |
| ZNF311 |
| ZNF334 |
| ZNF354C |
| ZNF358 |
| ZNF367 |
| ZNF385B |
| ZNF385C |
| ZNF470 |
| ZNF471 |
| ZNF502 |
| ZNF511 |
| ZNF536 |
| ZNF572 |
| ZNF577 |
| ZNF648 |
| ZNF662 |
| ZNF676 |
| ZNF682 |
| ZNF812 |
| ZNF879 |

| |
|---|
| ZPBP |
| ZPLD1 |
| ZSCAN18 |
| ZYG11A |

**Supplementary Table 7A.** Genes modulated by both BLU0588 treatment and shRNA knockdown, and with inverse modulation in FLC

| log2FC_Pos_Simon | Neg in both BLU0588 and shRNA | Overlap |
|---|---|---|
| MARCH4 | ROPN1L | AKR1B15 |
| MARCH10 | SCARA5 | ATP1B1 |
| AACS | CYP24A1 | CABYR |
| AAGAB | C11orf86 | CPLX2 |
| AAK1 | CD55 | CPS1 |
| ABCA12 | CPS1 | DDIT4 |
| ABCA2 | KRT86 | EVA1C |
| ABCA3 | PTGS2 | FSTL4 |
| ABCA7 | DUSP1 | GK |
| ABCB5 | TNFSF11 | GPRIN3 |
| ABCB8 | MPP1 | HPD |
| ABCC5 | CU639417.2 | KCNU1 |
| ABCF2 | BASP1 | KRT86 |
| ABHD17C | NR4A1 | KYNU |
| ABHD4 | SIK1 | MUC13 |
| ABLIM2 | SMIM9 | PBK |
| AC006132.1 | G6PC | PDE3A |
| AC008443.1 | GK | PDE4B |
| AC010547.9 | PDK4 | PPARGC1A |
| AC096677.1 | ADH4 | RHOBTB1 |
| AC145676.2 | PLA2G4A | S100P |
| ACAN | HEPACAM | SDCBP2 |
| ACKR3 | CA2 | SLC16A11 |
| ACLY | CPB2 | SLC22A11 |
| ACSL4 | PCK1 | SLC5A6 |
| ACYP1 | EVA1C | TESC |
| ADAM12 | RHOBTB1 | |
| ADAM22 | RBFOX3 | |
| ADAM32 | PDE3A | |
| ADAMDEC1 | TESC | |
| ADAMTS14 | SLC22A11 | |
| ADAMTS15 | PEX11A | |
| ADAMTS18 | SLC5A6 | |
| ADAMTS6 | PGC | |
| ADAMTS9 | PPARGC1A | |
| ADAMTSL5 | UGT1A4 | |
| ADAT1 | ID4 | |
| ADAT2 | LYPD3 | |
| ADCY2 | GPT | |
| ADD2 | GDF7 | |
| ADRA2A | PDE7B | |
| ADSSL1 | PDE4B | |
| AEN | ATP1B1 | |
| AF131215.5 | ALDH1L1 | |
| AFAP1 | CABYR | |
| AGA | HYAL1 | |
| AGR2 | PER2 | |
| AGRN | EPAS1 | |
| AHI1 | PBK | |
| AHNAK2 | SLC16A11 | |
| AHRR | AKR1B15 | |
| AIFM2 | SLC13A3 | |
| AIM1L | KHK | |
| AK8 | AVPI1 | |
| AKAP12 | FSTL4 | |
| AKR1B15 | DNAJC12 | |
| AKR1C3 | GPRIN3 | |
| AKTIP | KCNU1 | |
| AL133373.1 | S100P | |
| AL358813.2 | KYNU | |
| AL359878.1 | UGT1A3 | |
| ALDH18A1 | GGH | |
| ALDH1A2 | TCF7L1 | |

| | |
|---|---|
| ALDH1L2 | CPLX2 |
| ALDOA | SDCBP2 |
| ALG1 | MUC13 |
| ALPK3 | HPD |
| AMFR | HMOX1 |
| ANKRD18A | FMO5 |
| ANKRD22 | PREB |
| ANKRD29 | PAH |
| ANKRD52 | UGT2A3 |
| ANKS6 | PC |
| ANLN | FERMT1 |
| ANO2 | TTPA |
| ANO4 | ZFP36 |
| ANTXR1 | RND1 |
| ANXA2 | NAT6 |
| ANXA5 | APOB |
| AP1S3 | SRXN1 |
| APCDD1L | CDHR5 |
| APLN | TGFBR3L |
| APLP1 | BRD4 |
| APOO | UGT2B17 |
| AQPEP | CYP2C18 |
| AREG | AKR1C1 |
| ARG2 | DDIT4 |
| ARHGAP11A | KCNE3 |
| ARHGAP11B | |
| ARHGAP18 | |
| ARHGAP22 | |
| ARHGAP36 | |
| ARHGAP39 | |
| ARHGEF35 | |
| ARHGEF39 | |
| ARHGEF5 | |
| ARL14 | |
| ARL2BP | |
| ARMC9 | |
| ARNTL2 | |
| AS3MT | |
| ASB15 | |
| ASIC1 | |
| ASNS | |
| ASPHD1 | |
| ASRGL1 | |
| ATIC | |
| ATP1B1 | |
| ATP2A1 | |
| ATP2B4 | |
| ATP6V1B1 | |
| ATP8A2 | |
| ATP8B3 | |
| ATR | |
| AURKA | |
| B3GNT3 | |
| B3GNT5 | |
| B3GNTL1 | |
| B4GALNT1 | |
| B9D1 | |
| BAG2 | |
| BAG3 | |
| BAI2 | |
| BAIAP2L2 | |
| BBC3 | |
| BBS2 | |
| BCAR1 | |
| BCAT1 | |
| BCAT2 | |

| |
|---|
| BCL11A |
| BCL2L2-PABPN1 |
| BDKRB1 |
| BDKRB2 |
| BDNF |
| BEND3 |
| BEND6 |
| BEST3 |
| BFSP1 |
| BICD1 |
| BIRC5 |
| BLM |
| BLVRA |
| BMP6 |
| BMP8A |
| BMP8B |
| BNC2 |
| BRE |
| BRSK2 |
| BSG |
| BUB1 |
| BUB1B |
| C10orf128 |
| C10orf131 |
| C10orf2 |
| C10orf90 |
| C12orf39 |
| C12orf5 |
| C14orf183 |
| C15orf65 |
| C16orf46 |
| C16orf59 |
| C16orf93 |
| C18orf56 |
| C19orf54 |
| C1orf105 |
| C1orf198 |
| C1orf229 |
| C1QL1 |
| C20orf96 |
| C2CD4A |
| C2orf27A |
| C2orf66 |
| C2orf81 |
| C3orf36 |
| C3orf52 |
| C3orf67 |
| C4orf47 |
| C5AR2 |
| C5orf46 |
| C6orf163 |
| C6orf164 |
| C6orf195 |
| C8G |
| C8orf87 |
| C9orf116 |
| C9orf163 |
| C9orf57 |
| C9orf66 |
| CA12 |
| CA5A |
| CA5B |
| CA8 |
| CA9 |
| CABYR |
| CACNA1C |

| |
|---|
| CACNA1D |
| CACNB2 |
| CACNB4 |
| CALCA |
| CALCB |
| CAMK2N2 |
| CAP2 |
| CAPN11 |
| CATSPERB |
| CBX2 |
| CBX8 |
| CC2D2B |
| CCDC102B |
| CCDC113 |
| CCDC13 |
| CCDC136 |
| CCDC169 |
| CCDC170 |
| CCDC177 |
| CCDC64 |
| CCDC78 |
| CCDC80 |
| CCDC85A |
| CCDC88C |
| CCNA2 |
| CCNB1 |
| CCNB2 |
| CCNE1 |
| CCNF |
| CCNO |
| CCR8 |
| CCRN4L |
| CCT3 |
| CCT5 |
| CCT6A |
| CD109 |
| CD200 |
| CD248 |
| CD34 |
| CD46 |
| CDC20 |
| CDC20B |
| CDC25C |
| CDC45 |
| CDC6 |
| CDCA2 |
| CDCA5 |
| CDCA7 |
| CDH11 |
| CDH13 |
| CDH17 |
| CDH24 |
| CDH6 |
| CDK1 |
| CDK6 |
| CDKN2A |
| CDKN2B |
| CDKN3 |
| CDX1 |
| CDYL2 |
| CELSR3 |
| CEND1 |
| CENPF |
| CENPI |
| CENPK |
| CENPL |

| |
|---|
| CENPO |
| CEP128 |
| CEP152 |
| CEP55 |
| CERCAM |
| CERKL |
| CGA |
| CGREF1 |
| CHAC1 |
| CHCHD3 |
| CHEK1 |
| CHI3L1 |
| CHML |
| CHN1 |
| CHRDL2 |
| CHRNA5 |
| CHST10 |
| CHST6 |
| CHST8 |
| CIAPIN1 |
| CIDEC |
| CKB |
| CKS2 |
| CLCF1 |
| CLDN5 |
| CLEC18B |
| CLGN |
| CLIC5 |
| CLIP4 |
| CLSPN |
| CMTM4 |
| CNNM2 |
| COL10A1 |
| COL11A1 |
| COL15A1 |
| COL17A1 |
| COL1A1 |
| COL1A2 |
| COL22A1 |
| COL4A1 |
| COL4A2 |
| COL5A1 |
| COL5A2 |
| COL5A3 |
| COL6A3 |
| COL6A5 |
| COL8A1 |
| COL9A1 |
| COMP |
| COQ9 |
| CORIN |
| CORO2A |
| COX4I2 |
| CPA6 |
| CPD |
| CPE |
| CPLX1 |
| CPLX2 |
| CPNE5 |
| CPNE7 |
| CPS1 |
| CPXM1 |
| CRCP |
| CREB3L1 |
| CREG2 |
| CRLF2 |

| |
|---|
| CRMP1 |
| CRP |
| CRYM |
| CSGALNACT1 |
| CSMD2 |
| CSNK2A2 |
| CSPG4 |
| CTD-2600O9.1 |
| CTH |
| CTHRC1 |
| CXCL16 |
| CXorf36 |
| CYBA |
| CYCS |
| CYP17A1 |
| CYP19A1 |
| CYP1B1 |
| CYP21A2 |
| CYP27B1 |
| CYSTM1 |
| DAB2 |
| DAGLA |
| DCAF4L1 |
| DCDC2 |
| DCLK2 |
| DCX |
| DDIT4 |
| DDIT4L |
| DEFB1 |
| DEFB132 |
| DEPDC1 |
| DFNA5 |
| DGKI |
| DHRS7 |
| DIAPH3 |
| DIO2 |
| DKK1 |
| DKK2 |
| DKK4 |
| DLGAP4 |
| DLGAP5 |
| DNAAF1 |
| DNAAF3 |
| DNAH17 |
| DNAH5 |
| DNAH7 |
| DNAJC6 |
| DNER |
| DNM3 |
| DOCK5 |
| DOK4 |
| DOK5 |
| DPCD |
| DPEP1 |
| DPP10 |
| DPY19L1 |
| DPYSL4 |
| DSEL |
| DTL |
| DTNA |
| DUSP4 |
| DUSP8 |
| DYDC2 |
| E2F1 |
| E2F3 |
| E2F5 |

| |
|---|
| E2F7 |
| E2F8 |
| EARS2 |
| EBF1 |
| EBF2 |
| EBF3 |
| ECEL1 |
| ECT2 |
| EDA2R |
| EDIL3 |
| EEF1A2 |
| EFCAB12 |
| EFEMP1 |
| EFNA3 |
| EFNA5 |
| EGF |
| EGFL6 |
| EGLN3 |
| EHD2 |
| EIF4G3 |
| ELOVL3 |
| ELOVL7 |
| EMC7 |
| EME1 |
| ENDOD1 |
| ENO1 |
| ENOX1 |
| ENTPD1 |
| EPB41L1 |
| EPDR1 |
| EPHA10 |
| EPHX4 |
| EPS8L3 |
| ERBB2 |
| ERC2 |
| ERCC6L |
| EREG |
| ERMP1 |
| ESCO2 |
| ESM1 |
| ESRRG |
| ETV1 |
| ETV4 |
| EVA1C |
| EVC |
| EXO1 |
| EYS |
| EZH2 |
| EZR |
| F13A1 |
| F2RL1 |
| F2RL3 |
| FA2H |
| FAHD2B |
| FAM101A |
| FAM111B |
| FAM115C |
| FAM127A |
| FAM127B |
| FAM127C |
| FAM132B |
| FAM135A |
| FAM155A |
| FAM155B |
| FAM167A |
| FAM169B |

| |
|---|
| FAM177B |
| FAM179A |
| FAM188B |
| FAM189B |
| FAM196A |
| FAM19A5 |
| FAM227A |
| FAM64A |
| FAM71F2 |
| FAM78B |
| FAM81A |
| FAM83A |
| FAM86A |
| FAM86B1 |
| FANCI |
| FAP |
| FASTKD1 |
| FAT1 |
| FBN1 |
| FBXL18 |
| FBXL8 |
| FBXO25 |
| FBXO36 |
| FBXO39 |
| FGF7 |
| FHDC1 |
| FKBP11 |
| FKBP14 |
| FKBP1C |
| FLAD1 |
| FLVCR1 |
| FMNL2 |
| FMNL3 |
| FNDC1 |
| FNDC3A |
| FOXC1 |
| FOXF2 |
| FOXL1 |
| FOXM1 |
| FOXQ1 |
| FOXS1 |
| FRRS1L |
| FRZB |
| FSD1L |
| FST |
| FSTL4 |
| FTO |
| FURIN |
| FZD10 |
| FZD2 |
| FZD6 |
| FZD8 |
| G6PD |
| GABRB3 |
| GABRD |
| GABRE |
| GABRQ |
| GAD1 |
| GAL3ST4 |
| GALNT10 |
| GALNT12 |
| GALNT5 |
| GALNT7 |
| GALNTL6 |
| GALR2 |
| GAPDH |

| |
|---|
| GAREML |
| GARS |
| GAS8 |
| GATSL2 |
| GCNT3 |
| GDA |
| GDF15 |
| GGN |
| GHRHR |
| GINS1 |
| GJA5 |
| GJC1 |
| GK |
| GLA |
| GLCE |
| GLIS3 |
| GLP1R |
| GLRB |
| GLRX |
| GLS |
| GMDS |
| GNA12 |
| GNAL |
| GNAZ |
| GNB5 |
| GNG4 |
| GOLGA6L9 |
| GOLM1 |
| GOLT1B |
| GOT1 |
| GPATCH4 |
| GPC2 |
| GPD1L |
| GPI |
| GPR1 |
| GPR107 |
| GPR150 |
| GPR37 |
| GPR4 |
| GPR56 |
| GPR63 |
| GPR64 |
| GPR68 |
| GPR97 |
| GPRIN1 |
| GPRIN3 |
| GPSM2 |
| GPX8 |
| GRAMD1A |
| GRAMD1B |
| GRAMD4 |
| GREM2 |
| GRIA3 |
| GRK7 |
| GRM5 |
| GRM7 |
| GSDMC |
| GSN |
| GTF2E1 |
| GTF2IRD1 |
| GTPBP4 |
| GUCY2C |
| GULP1 |
| GXYLT2 |
| H2AFY2 |
| HAGHL |

| |
|---|
| HAPLN1 |
| HAS3 |
| HAVCR1 |
| HCN1 |
| HEATR2 |
| HEG1 |
| HELLS |
| HES2 |
| HES4 |
| HEYL |
| HHATL |
| HHIPL2 |
| HHLA3 |
| HIGD1B |
| HIST1H1A |
| HIST1H1C |
| HIST1H1T |
| HIST1H2AA |
| HIST1H2AB |
| HIST1H2AC |
| HIST1H2AG |
| HIST1H2AH |
| HIST1H2AI |
| HIST1H2AK |
| HIST1H2BB |
| HIST1H2BG |
| HIST1H2BJ |
| HIST1H2BK |
| HIST1H2BN |
| HIST1H3A |
| HIST1H3D |
| HIST1H4I |
| HIST2H3D |
| HJURP |
| HK2 |
| HKDC1 |
| HMGB3 |
| HNF1B |
| HOMER1 |
| HOPX |
| HOXD1 |
| HOXD3 |
| HOXD9 |
| HPD |
| HSF4 |
| HSP90AA1 |
| HSP90AB1 |
| HSPA12A |
| HSPA12B |
| HSPA4L |
| HSPA5 |
| HSPB6 |
| HSPH1 |
| HTATIP2 |
| HTR2A |
| HTRA1 |
| HYOU1 |
| IARS |
| ICAM5 |
| IER3 |
| IER5L |
| IGDCC4 |
| IGF2BP3 |
| IGFBPL1 |
| IGSF9B |
| IL17D |

| |
|---|
| IL2RA |
| IL31RA |
| IL32 |
| INPP4B |
| INSL4 |
| INTS9 |
| INTU |
| IQCA1 |
| IQCD |
| IQCE |
| IQCK |
| IQGAP3 |
| IRAK1 |
| IRF4 |
| IRX3 |
| IRX5 |
| ISG15 |
| ISG20 |
| ITGA11 |
| ITGA2 |
| ITGA6 |
| ITGA7 |
| ITGAV |
| ITGB4 |
| ITPKA |
| JAG1 |
| JAG2 |
| JPH1 |
| KAAG1 |
| KAL1 |
| KCNE1L |
| KCNE4 |
| KCNF1 |
| KCNJ5 |
| KCNJ6 |
| KCNK9 |
| KCNN3 |
| KCNQ3 |
| KCNU1 |
| KDELR3 |
| KIAA0100 |
| KIAA0319 |
| KIAA0556 |
| KIAA0895L |
| KIAA1024 |
| KIAA1199 |
| KIAA1211L |
| KIAA1244 |
| KIAA1324 |
| KIAA1462 |
| KIAA1522 |
| KIAA1549 |
| KIAA1549L |
| KIF14 |
| KIF18B |
| KIF20A |
| KIF21B |
| KIF23 |
| KIF24 |
| KIF26B |
| KIF2C |
| KIF3A |
| KIF4A |
| KIF5A |
| KLC2 |
| KLF5 |

| |
|---|
| KLHDC7B |
| KLHL21 |
| KLHL29 |
| KNDC1 |
| KPNA2 |
| KRT222 |
| KRT7 |
| KRT81 |
| KRT86 |
| KSR1 |
| KYNU |
| LAMA1 |
| LAMA3 |
| LAMA4 |
| LAMB4 |
| LAMC1 |
| LANCL1 |
| LANCL3 |
| LAPTM4B |
| LBP |
| LCN2 |
| LDHB |
| LEF1 |
| LEP |
| LETM2 |
| LGALS3 |
| LGI2 |
| LHCGR |
| LHFPL5 |
| LHX6 |
| LIF |
| LIMCH1 |
| LIMK1 |
| LIMK2 |
| LIPH |
| LITAF |
| LOH12CR1 |
| LONRF2 |
| LOX |
| LOXL2 |
| LPAR3 |
| LPAR4 |
| LPCAT1 |
| LPL |
| LPPR4 |
| LRP8 |
| LRRC1 |
| LRRC16A |
| LRRC37A3 |
| LRRC69 |
| LRRC73 |
| LRRC8E |
| LSAMP |
| LTBP2 |
| LUZP2 |
| LYPD1 |
| LZTS1 |
| MAB21L3 |
| MAFK |
| MALL |
| MANF |
| MAP1A |
| MAP1B |
| MAP1LC3B |
| MAP1LC3B2 |
| MAP3K9 |

| |
|---|
| MAP9 |
| MAPK12 |
| MAPT |
| MARS2 |
| MATN3 |
| MBOAT4 |
| MCAM |
| MCF2L2 |
| MCM10 |
| MCM4 |
| MCTP2 |
| MECOM |
| MED9 |
| MELK |
| MEX3B |
| MFSD6 |
| MGAT3 |
| MGAT5B |
| MGC4294 |
| MID1 |
| MKI67 |
| MLEC |
| MMP1 |
| MMP10 |
| MMP11 |
| MMP14 |
| MMP2 |
| MN1 |
| MND1 |
| MPHOSPH6 |
| MPP3 |
| MPP4 |
| MPV17 |
| MPV17L2 |
| MPZ |
| MRAP |
| MRAP2 |
| MRAS |
| MRPS12 |
| MRVI1 |
| MST1R |
| MT-ATP8 |
| MT-ND5 |
| MT-ND6 |
| MT3 |
| MTFR2 |
| MTHFD1L |
| MTRNR2L2 |
| MUC13 |
| MUC3A |
| MUC5B |
| MURC |
| MVP |
| MYBPHL |
| MYCN |
| MYEF2 |
| MYH4 |
| MYO1E |
| MYO5C |
| MYOF |
| MYOM3 |
| MYRF |
| N4BP3 |
| NAV1 |
| NAV2 |
| NBEA |

| |
|---|
| NCAPG |
| NCR3LG1 |
| NDUFA4L2 |
| NEB |
| NEIL3 |
| NEK2 |
| NETO2 |
| NHS |
| NIPAL2 |
| NKX1-2 |
| NKX2-3 |
| NME1 |
| NMNAT2 |
| NOL3 |
| NOMO1 |
| NOMO2 |
| NOMO3 |
| NOTCH3 |
| NOV |
| NOVA1 |
| NOX1 |
| NOX4 |
| NPAS2 |
| NPCDR1 |
| NPFFR2 |
| NPNT |
| NPTX2 |
| NPTXR |
| NPY5R |
| NQO1 |
| NR4A2 |
| NRG2 |
| NRIP2 |
| NRXN3 |
| NT5C3A |
| NT5DC2 |
| NTM |
| NTS |
| NUF2 |
| NXPH4 |
| OAT |
| OGDH |
| OGFOD1 |
| OLAH |
| OLFML2A |
| OLFML2B |
| OPHN1 |
| OPN3 |
| OR1F1 |
| OR2A7 |
| OR2AG2 |
| OR2B6 |
| OR2D2 |
| OR51E1 |
| OR51E2 |
| OR6A2 |
| ORC1 |
| ORC6 |
| OSBPL3 |
| OSBPL9 |
| OSMR |
| OSR1 |
| OSR2 |
| OTOG |
| OTUB2 |
| OTUD7A |

| |
|---|
| OTX1 |
| OXCT1 |
| P4HA2 |
| PAEP |
| PAK3 |
| PALLD |
| PAPPA |
| PAQR4 |
| PAQR5 |
| PARM1 |
| PARPBP |
| PBK |
| PBX1 |
| PCDH17 |
| PCDHB10 |
| PCDHB11 |
| PCDHB13 |
| PCDHB8 |
| PCDHGA1 |
| PCDHGA12 |
| PCDHGA4 |
| PCDHGA5 |
| PCDHGA7 |
| PCDHGA8 |
| PCDHGB1 |
| PCDHGB2 |
| PCDHGC5 |
| PCDP1 |
| PCED1B |
| PCM1 |
| PCNT |
| PCNXL2 |
| PCSK1 |
| PDE10A |
| PDE1C |
| PDE3A |
| PDE3B |
| PDE4A |
| PDE4B |
| PDE4D |
| PDF |
| PDGFA |
| PDGFRB |
| PDGFRL |
| PDK1 |
| PDLIM7 |
| PDX1 |
| PDZD2 |
| PFKFB2 |
| PFKM |
| PFKP |
| PHEX |
| PHF17 |
| PHLDA2 |
| PHLDA3 |
| PHPT1 |
| PIAS3 |
| PIGA |
| PIP4K2C |
| PITPNM1 |
| PITX1 |
| PKP1 |
| PLA2G2A |
| PLA2G2C |
| PLA2G4E |
| PLCB4 |

| |
|---|
| PLCD3 |
| PLCE1 |
| PLCH1 |
| PLEKHA8 |
| PLEKHG2 |
| PLEKHH1 |
| PLEKHN1 |
| PLK1 |
| PLOD3 |
| PLP2 |
| PLVAP |
| PLXDC1 |
| PM20D2 |
| PMEPA1 |
| PMFBP1 |
| PNMA1 |
| PODNL1 |
| PODXL |
| POLN |
| POLR1A |
| POLR2C |
| POLR3G |
| PON2 |
| POSTN |
| POU5F1 |
| PPA1 |
| PPARGC1A |
| PPAT |
| PPIA |
| PPP1R13L |
| PPP1R36 |
| PPP1R3D |
| PPP1R3G |
| PPP2R2C |
| PRAMEF10 |
| PRAMEF2 |
| PRAMEF4 |
| PRC1 |
| PRDM6 |
| PRDM7 |
| PRKACA |
| PRND |
| PRODH |
| PRR15 |
| PRR16 |
| PRR26 |
| PRRG3 |
| PRRX1 |
| PRSS27 |
| PSMD14 |
| PSME3 |
| PSORS1C1 |
| PTCHD4 |
| PTGFR |
| PTGFRN |
| PTP4A3 |
| PTPDC1 |
| PTPLA |
| PTPN14 |
| PTPN5 |
| PTPRM |
| PTPRR |
| PTPRU |
| PVRL1 |
| PYCR1 |
| PYGB |

| |
|---|
| QRFPR |
| RAB3A |
| RAB3B |
| RAB6B |
| RACGAP1 |
| RAET1E |
| RANBP17 |
| RAP1GAP |
| RARRES1 |
| RASD1 |
| RASD2 |
| RASEF |
| RASGEF1A |
| RASGRF2 |
| RASL11B |
| RASL12 |
| RASSF6 |
| RASSF9 |
| RBM20 |
| RBM24 |
| RBM44 |
| RBPMS |
| RECQL4 |
| RELL2 |
| RFX8 |
| RGAG4 |
| RGCC |
| RGS17 |
| RGS5 |
| RGS6 |
| RGS9 |
| RHBDD2 |
| RHOBTB1 |
| RHOBTB2 |
| RHOF |
| RHOQ |
| RHPN1 |
| RMI2 |
| RNF157 |
| RNFT2 |
| ROBO1 |
| ROR2 |
| RP1-102H19.8 |
| RP11-17M16.1 |
| RP11-212D19.4 |
| RP11-248J23.6 |
| RP11-366L20.2 |
| RP11-796G6.2 |
| RPGR |
| RPGRIP1L |
| RPP40 |
| RPRML |
| RPS6KA2 |
| RPS6KL1 |
| RRM2 |
| RRS1 |
| RUNX1 |
| S100A3 |
| S100P |
| SAPCD2 |
| SATB2 |
| SCG2 |
| SCIN |
| SCML4 |
| SCN4A |
| SDCBP2 |

| |
|---|
| SDIM1 |
| SDSL |
| SEMA3F |
| SEMA3G |
| SEMA5B |
| SERHL2 |
| SERPINB3 |
| SERTAD4 |
| SEZ6L2 |
| SFMBT2 |
| SFN |
| SFRP2 |
| SFRP4 |
| SGIP1 |
| SGPP2 |
| SH3PXD2B |
| SH3RF3 |
| SHC1 |
| SHCBP1 |
| SHISA2 |
| SHROOM4 |
| SIAE |
| SIX1 |
| SIX4 |
| SKA1 |
| SLC16A11 |
| SLC16A14 |
| SLC22A11 |
| SLC22A12 |
| SLC22A15 |
| SLC22A23 |
| SLC22A5 |
| SLC25A12 |
| SLC25A15 |
| SLC25A24 |
| SLC26A2 |
| SLC26A7 |
| SLC26A9 |
| SLC27A4 |
| SLC2A1 |
| SLC2A5 |
| SLC35B4 |
| SLC35C1 |
| SLC35E4 |
| SLC35G2 |
| SLC36A1 |
| SLC38A1 |
| SLC44A3 |
| SLC45A1 |
| SLC4A11 |
| SLC52A3 |
| SLC5A6 |
| SLC6A11 |
| SLC6A17 |
| SLC6A3 |
| SLC6A6 |
| SLC6A8 |
| SLC6A9 |
| SLC7A1 |
| SLC7A11 |
| SLC7A2 |
| SLC7A5 |
| SLC7A6 |
| SLCO1C1 |
| SLCO2A1 |
| SLCO5A1 |

| |
|---|
| SLIT2 |
| SLITRK4 |
| SMAD2 |
| SMCO2 |
| SMKR1 |
| SMOC2 |
| SMOX |
| SNAP25 |
| SNPH |
| SOGA1 |
| SORT1 |
| SOX11 |
| SOX12 |
| SOX21 |
| SOX4 |
| SP140 |
| SP6 |
| SPA17 |
| SPATA17 |
| SPATC1L |
| SPATS2 |
| SPECC1 |
| SPINK1 |
| SPIRE1 |
| SPOCD1 |
| SPOCK1 |
| SPRED3 |
| SPTBN5 |
| SPTSSA |
| SRGAP1 |
| SRM |
| SRPX2 |
| SRRM3 |
| SRSF12 |
| ST6GALNAC4 |
| ST8SIA5 |
| STAMBPL1 |
| STC1 |
| STC2 |
| STEAP1 |
| STEAP1B |
| STK32B |
| STRA6 |
| STRIP2 |
| STS |
| STXBP1 |
| STXBP4 |
| STXBP5 |
| SULF1 |
| SULF2 |
| SULT1C2 |
| SULT2B1 |
| SULT4A1 |
| SYNDIG1 |
| SYT13 |
| TAF4B |
| TAF6 |
| TAF7L |
| TAGLN2 |
| TANC2 |
| TAS2R3 |
| TAS2R4 |
| TAS2R5 |
| TAX1BP3 |
| TBC1D16 |
| TBC1D30 |

| |
|---|
| TBC1D31 |
| TC2N |
| TCF23 |
| TCN1 |
| TCTN3 |
| TEAD4 |
| TENM4 |
| TESC |
| TEX9 |
| TG |
| TGFB2 |
| TGFBR1 |
| TGIF1 |
| TGM2 |
| TGM3 |
| THBS2 |
| THBS4 |
| THSD7A |
| THY1 |
| TICAM2 |
| TICRR |
| TIMP1 |
| TK1 |
| TLCD1 |
| TLCD2 |
| TLDC1 |
| TLE6 |
| TLL2 |
| TM4SF5 |
| TMC1 |
| TMC5 |
| TMC7 |
| TMED3 |
| TMEM104 |
| TMEM108 |
| TMEM119 |
| TMEM132A |
| TMEM136 |
| TMEM144 |
| TMEM145 |
| TMEM156 |
| TMEM163 |
| TMEM165 |
| TMEM178B |
| TMEM2 |
| TMEM200A |
| TMEM217 |
| TMEM245 |
| TMEM246 |
| TMEM45A |
| TMEM55A |
| TMEM59L |
| TMOD1 |
| TMTC2 |
| TNFRSF10A |
| TNFRSF12A |
| TNFRSF21 |
| TNFRSF4 |
| TNFSF4 |
| TNRC6C |
| TOMM40L |
| TOP2A |
| TOX |
| TPBG |
| TPBGL |
| TPM4 |

| |
|---|
| TRAF2 |
| TRIM31 |
| TRIM40 |
| TRIM46 |
| TRIM59 |
| TRIM7 |
| TRIM8 |
| TRIP13 |
| TRPC4 |
| TRPC6 |
| TRPM2 |
| TRPS1 |
| TSHZ2 |
| TSPAN13 |
| TTC26 |
| TTC39A |
| TTK |
| TTL |
| TTYH2 |
| TUBA4A |
| TXNDC17 |
| TXNRD1 |
| TYMS |
| UBE2C |
| UBE2T |
| UBFD1 |
| UBXN10 |
| UCHL1 |
| UCN |
| UGT2B11 |
| ULBP1 |
| ULBP2 |
| ULBP3 |
| UNC119B |
| UNC13A |
| UNC13B |
| UNC5B |
| UNC5D |
| URB1 |
| USB1 |
| USH1C |
| USP49 |
| USP54 |
| UTS2B |
| VARS |
| VASH2 |
| VAX2 |
| VCAN |
| VLDLR |
| VSIG1 |
| VSX1 |
| VWA7 |
| VWF |
| WDR13 |
| WIPF3 |
| WNT16 |
| WWC1 |
| XKR3 |
| XKR6 |
| XKRX |
| XPR1 |
| XRCC2 |
| YIPF6 |
| YWHAG |
| ZBTB38 |
| ZC3H12B |

| |
|---|
| ZFHX3 |
| ZFP69B |
| ZHX1-C8ORF76 |
| ZIC2 |
| ZMAT3 |
| ZMYND15 |
| ZNF154 |
| ZNF215 |
| ZNF23 |
| ZNF233 |
| ZNF365 |
| ZNF382 |
| ZNF385D |
| ZNF57 |
| ZNF683 |
| ZNF695 |
| ZNF703 |
| ZNF724P |
| ZNF827 |
| ZP3 |
| ZSCAN5A |
| ZSCAN9 |
| ZWINT |
| ZXDB |

**Supplementary Table 7B.** Genes modulated by both BLU0588 treatment and shRNA knockdown, and with inverse modulation in FLC

| log2FC_Neg_Simon | Pos in both BLU0588 and shRNA | Overlap |
|---|---|---|
| MARCH2 | ADAMTSL2 | ADAMTSL2 |
| SEPT4 | CHRNA3 | CAPN5 |
| A1BG | BMP7 | CYP2S1 |
| A1CF | CLIC5 | EBF4 |
| AADAC | SHH | ECM1 |
| AADAT | FGF20 | IGSF9 |
| ABAT | WNT6 | LIMD2 |
| ABCA13 | SCARA3 | NGFR |
| ABCA6 | BCAM | RGL1 |
| ABCA9 | GALNT5 | SHH |
| ABCB11 | IL17RD | TP53INP1 |
| ABCG2 | RIMKLA | VPS37D |
| ABCG5 | GSN | YPEL1 |
| ABCG8 | EFNB3 | |
| ABHD1 | TNNT1 | |
| ABHD15 | ZNF608 | |
| ABHD5 | MFGE8 | |
| AC007405.2 | FILIP1L | |
| AC010368.2 | CRIP2 | |
| AC011484.1 | FN1 | |
| AC011841.1 | PRRT4 | |
| AC018755.1 | YPEL1 | |
| AC021860.1 | DACT1 | |
| AC023590.1 | NGFR | |
| AC103801.2 | RGL1 | |
| AC104667.3 | EMID1 | |
| AC104809.3 | FLRT3 | |
| AC110781.3 | ZDHHC2 | |
| ACAA1 | TUBA1A | |
| ACAA2 | IGSF9 | |
| ACACB | PNMA2 | |
| ACADL | EBF4 | |
| ACADM | CD8B | |
| ACADS | CERCAM | |
| ACADSB | PAM | |
| ACAT1 | HAPLN3 | |
| ACAT2 | KIRREL | |
| ACBD4 | ANGPTL2 | |
| ACE2 | CACNA1S | |
| ACKR4 | KRT20 | |
| ACOT1 | KLK5 | |
| ACOT12 | GPR173 | |
| ACOT13 | SYT11 | |
| ACOT2 | SCRN1 | |
| ACOT4 | AMOT | |
| ACOX2 | ANK1 | |
| ACP5 | SLC34A2 | |
| ACSF2 | TMEM173 | |
| ACSL1 | BMF | |
| ACSL5 | SALL2 | |
| ACSL6 | GLB1L2 | |
| ACSM2A | KCNH3 | |
| ACSM2B | SLC4A11 | |
| ACSM3 | NACAD | |
| ACSM5 | VWA5B2 | |
| ACSS2 | GPR161 | |
| ACY3 | PMAIP1 | |
| ADAM11 | REEP2 | |
| ADAMTS13 | SULF2 | |
| ADAMTSL2 | STK32B | |
| ADAMTSL3 | RASSF4 | |
| ADAMTSL4 | PDGFB | |
| ADAP1 | HSPG2 | |
| ADCY1 | SBK1 | |
| ADCY10 | MT1X | |
| ADCYAP1R1 | AMIGO2 | |
| ADH1A | EGR1 | |

| | |
|---|---|
| ADH1B | TIMP2 |
| ADH4 | ZNF423 |
| ADH6 | SERPINE2 |
| ADH7 | MEIS3 |
| ADI1 | ASAP3 |
| ADIRF | LIMD2 |
| ADM | SEMA4F |
| ADRA1A | DUSP6 |
| ADRA1B | EEPD1 |
| ADRA2B | SHROOM3 |
| ADRB1 | ECM1 |
| ADRB2 | CHST3 |
| ADTRP | MYL9 |
| AFF3 | C15orf52 |
| AFM | CMTM3 |
| AGBL2 | DNAJC18 |
| AGL | CLIP3 |
| AGMAT | RCOR2 |
| AGMO | B3GNT7 |
| AGPAT2 | LRRC75A |
| AGTR1 | VPS37D |
| AGXT | GAS6 |
| AGXT2 | CACNB3 |
| AHSG | PCDHB15 |
| AIG1 | OLFML2A |
| AJAP1 | PCDHB8 |
| AKAP3 | MEX3A |
| AKR1A1 | PIK3IP1 |
| AKR1C4 | AHRR |
| AKR1D1 | CPNE2 |
| AL078585.1 | PROCR |
| AL590714.1 | PDLIM4 |
| ALAD | TP53INP1 |
| ALAS1 | NKD2 |
| ALAS2 | PLCB1 |
| ALB | DPYSL2 |
| ALDH1L1 | MGAT5 |
| ALDH2 | MAPK8IP1 |
| ALDH5A1 | PPL |
| ALDH6A1 | GJB4 |
| ALDH7A1 | EPPK1 |
| ALDH9A1 | PCDHGC3 |
| ALDOC | PDLIM1 |
| ALLC | IMPDH1 |
| ALPL | CAPN5 |
| AMACR | SORT1 |
| AMBP | PTPN14 |
| AMHR2 | CYP2S1 |
| AMN | ORAI2 |
| AMOTL2 | TPM1 |
| AMT | PCDHB2 |
| ANG | |
| ANGPTL6 | |
| ANGPTL7 | |
| ANK3 | |
| ANKRD20A3 | |
| ANKRD55 | |
| ANO1 | |
| ANPEP | |
| ANXA10 | |
| ANXA3 | |
| ANXA9 | |
| AOC1 | |
| AP1M2 | |
| APBA1 | |
| APH1A | |
| APOA1 | |
| APOA2 | |
| APOA5 | |
| APOBEC3A | |

| |
|---|
| APOC1 |
| APOC3 |
| APOF |
| APOH |
| APOL1 |
| APOL6 |
| APOM |
| AQP11 |
| AQP7 |
| AQP9 |
| AR |
| ARC |
| ARHGAP20 |
| ARHGEF10L |
| ARHGEF26 |
| ARHGEF38 |
| ARHGEF40 |
| ARID3C |
| ARID5A |
| ARL4D |
| ARMC3 |
| ARMC5 |
| ARRDC3 |
| ARSF |
| ART3 |
| ART4 |
| ART5 |
| ARVCF |
| ASB4 |
| ASB9 |
| ASCL2 |
| ASGR1 |
| ASMTL |
| ASPA |
| ASPDH |
| ASTL |
| ASTN1 |
| ASXL3 |
| ATAD3C |
| ATF3 |
| ATF5 |
| ATF7IP2 |
| ATHL1 |
| ATOH8 |
| ATP11C |
| ATP13A4 |
| ATP7B |
| ATRNL1 |
| AUTS2 |
| AVPR1A |
| AXL |
| AZGP1 |
| AZU1 |
| B3GAT1 |
| B3GAT2 |
| B3GNT8 |
| B4GALNT3 |
| BAALC |
| BACH2 |
| BAI3 |
| BAIAP2 |
| BAIAP3 |
| BBOX1 |
| BCHE |
| BCL2A1 |
| BCL6 |
| BCO2 |
| BDH1 |
| BEND4 |
| BEND7 |

| |
|---|
| BEX1 |
| BEX4 |
| BHMT |
| BHMT2 |
| BLK |
| BLNK |
| BMP10 |
| BMP3 |
| BMP5 |
| BMPER |
| BNC1 |
| BOK |
| BPHL |
| BPI |
| BRINP2 |
| BRIP1 |
| BTD |
| BTNL8 |
| BZRAP1 |
| C10orf11 |
| C10orf25 |
| C10orf67 |
| C11orf31 |
| C11orf35 |
| C11orf54 |
| C11orf65 |
| C11orf71 |
| C11orf85 |
| C12orf42 |
| C12orf68 |
| C12orf74 |
| C14orf164 |
| C14orf180 |
| C14orf80 |
| C15orf26 |
| C15orf43 |
| C16orf96 |
| C19orf12 |
| C19orf38 |
| C19orf66 |
| C19orf71 |
| C19orf80 |
| C1orf115 |
| C1orf116 |
| C1orf162 |
| C1orf168 |
| C1orf172 |
| C1orf173 |
| C1orf21 |
| C1orf210 |
| C1orf226 |
| C1orf53 |
| C1QB |
| C1QC |
| C1R |
| C1RL |
| C1S |
| C2 |
| C21orf2 |
| C21orf33 |
| C21orf37 |
| C21orf62 |
| C21orf67 |
| C21orf88 |
| C21orf90 |
| C21orf91 |
| C2CD4B |
| C2orf40 |
| C2orf54 |
| C2orf72 |

| |
|---|
| C4BPA |
| C5AR1 |
| C5orf27 |
| C5orf49 |
| C6 |
| C7orf10 |
| C8A |
| C8B |
| C8orf82 |
| C9 |
| C9orf43 |
| C9orf72 |
| C9orf96 |
| CA1 |
| CA14 |
| CABLES1 |
| CABP4 |
| CACNA2D2 |
| CADM2 |
| CALN1 |
| CAMK2B |
| CAMK2N1 |
| CAMK4 |
| CAND2 |
| CAPN12 |
| CAPN5 |
| CAPZA3 |
| CARHSP1 |
| CASC10 |
| CASR |
| CAT |
| CBFA2T3 |
| CBLC |
| CBLN4 |
| CBR4 |
| CBS |
| CBX7 |
| CCBE1 |
| CCDC150 |
| CCDC151 |
| CCDC152 |
| CCDC158 |
| CCDC178 |
| CCDC64B |
| CCDC68 |
| CCDC73 |
| CCL14 |
| CCL16 |
| CCL2 |
| CCL23 |
| CCL24 |
| CCL28 |
| CCL3 |
| CCL4L1 |
| CCNB1IP1 |
| CCR1 |
| CCR9 |
| CCT6B |
| CD14 |
| CD160 |
| CD163 |
| CD180 |
| CD1C |
| CD1D |
| CD1E |
| CD244 |
| CD274 |
| CD300A |
| CD300C |
| CD300E |

| |
|---|
| CD300LB |
| CD300LG |
| CD302 |
| CD36 |
| CD4 |
| CD5L |
| CD81 |
| CD82 |
| CD83 |
| CD9 |
| CDH19 |
| CDH2 |
| CDH23 |
| CDH4 |
| CDHR5 |
| CDK18 |
| CDK3 |
| CEACAM3 |
| CEACAM4 |
| CEACAM6 |
| CEBPA |
| CEBPD |
| CECR2 |
| CENPV |
| CERS4 |
| CES1 |
| CES2 |
| CES3 |
| CES4A |
| CES5A |
| CETP |
| CFB |
| CFD |
| CFHR1 |
| CFHR2 |
| CFHR3 |
| CFHR4 |
| CFHR5 |
| CFL2 |
| CFP |
| CFTR |
| CGN |
| CGNL1 |
| CHAD |
| CHADL |
| CHDC2 |
| CHDH |
| CHN2 |
| CHRDL1 |
| CHRM2 |
| CHRNA4 |
| CHST13 |
| CHST4 |
| CHST7 |
| CIDEB |
| CILP |
| CISH |
| CLDN1 |
| CLDN10 |
| CLDN14 |
| CLDN19 |
| CLDN3 |
| CLDN4 |
| CLEC1B |
| CLEC3B |
| CLEC4C |
| CLEC4D |
| CLEC4E |
| CLEC4G |
| CLEC4M |

| |
|---|
| CLVS2 |
| CMA1 |
| CMBL |
| CMPK2 |
| CMTM2 |
| CMTM8 |
| CMYA5 |
| CNBD1 |
| CNDP1 |
| CNGA1 |
| CNKSR2 |
| CNPY3 |
| CNST |
| CNTLN |
| CNTN5 |
| COBLL1 |
| COL11A2 |
| COL18A1 |
| COL25A1 |
| COL27A1 |
| COL28A1 |
| COL6A6 |
| COLEC10 |
| COLEC11 |
| COMT |
| COQ10A |
| COX6A2 |
| CP |
| CPA3 |
| CPAMD8 |
| CPB2 |
| CPEB3 |
| CPED1 |
| CPN1 |
| CPN2 |
| CPNE6 |
| CR1 |
| CR1L |
| CRHBP |
| CRLS1 |
| CRYAA |
| CRYL1 |
| CSAD |
| CSF3 |
| CSF3R |
| CSRNP1 |
| CTD-2014B16.3 |
| CTF1 |
| CTNNA3 |
| CTSG |
| CTSL |
| CTTNBP2 |
| CUX2 |
| CX3CL1 |
| CXADR |
| CXCL1 |
| CXCL12 |
| CXCL14 |
| CXCL2 |
| CXCL3 |
| CXCR1 |
| CXCR2 |
| CXorf22 |
| CXorf66 |
| CYB5A |
| CYP11A1 |
| CYP1A1 |
| CYP1A2 |
| CYP26A1 |
| CYP26B1 |

| |
|---|
| CYP27A1 |
| CYP27C1 |
| CYP2A13 |
| CYP2A6 |
| CYP2A7 |
| CYP2B6 |
| CYP2C19 |
| CYP2C8 |
| CYP2C9 |
| CYP2D6 |
| CYP2J2 |
| CYP2S1 |
| CYP39A1 |
| CYP3A4 |
| CYP3A43 |
| CYP3A5 |
| CYP3A7 |
| CYP4A11 |
| CYP4A22 |
| CYP4F12 |
| CYP4F2 |
| CYP4F3 |
| CYP4V2 |
| CYP4X1 |
| CYP4Z1 |
| CYP7A1 |
| CYP7B1 |
| CYP8B1 |
| DAB1 |
| DACH1 |
| DAK |
| DAO |
| DBH |
| DBI |
| DCAF11 |
| DCHS2 |
| DCT |
| DCXR |
| DDC |
| DDT |
| DDTL |
| DDX25 |
| DEFA3 |
| DEPDC7 |
| DGAT2 |
| DGCR6L |
| DHCR24 |
| DHCR7 |
| DHFR |
| DHODH |
| DHRS1 |
| DHRS12 |
| DHRS2 |
| DHRS4 |
| DHRS4L2 |
| DHTKD1 |
| DIO1 |
| DIRAS3 |
| DLEU7 |
| DLGAP2 |
| DMD |
| DMGDH |
| DMRTA1 |
| DNAH6 |
| DNAJC12 |
| DNAJC19 |
| DNALI1 |
| DNASE1L3 |
| DNMT3L |
| DOK2 |

| |
|---|
| DOK6 |
| DPEP2 |
| DPEP3 |
| DPF3 |
| DPP4 |
| DPPA4 |
| DPT |
| DPYD |
| DPYS |
| DRD1 |
| DSC2 |
| DSCAM |
| DSG1 |
| DSG4 |
| DTX1 |
| DUSP10 |
| DUSP14 |
| DYNLRB2 |
| EBF4 |
| EBI3 |
| EBP |
| EBPL |
| ECH1 |
| ECHDC2 |
| ECHDC3 |
| ECI2 |
| ECM1 |
| EDAR |
| EDN1 |
| EDNRB |
| EFCAB1 |
| EFCC1 |
| EFHD1 |
| EFNA2 |
| EHD3 |
| EHHADH |
| EIF1AY |
| ELAC1 |
| ELAVL4 |
| ELF5 |
| ELFN1 |
| ELOVL2 |
| ELOVL6 |
| EMILIN3 |
| EMP2 |
| EMR1 |
| EMR3 |
| ENDOU |
| ENHO |
| ENO3 |
| ENPEP |
| ENPP3 |
| ENPP6 |
| ENPP7 |
| ENTPD5 |
| ENTPD8 |
| EPB41L4B |
| EPHA1 |
| EPHA7 |
| EPHX1 |
| EPHX2 |
| EPO |
| EPOR |
| EPS8L2 |
| ERF |
| ERVFRD-1 |
| ESPN |
| ESPNL |
| ESR1 |
| ESRP1 |

| |
|---|
| ETFDH |
| ETNK2 |
| EVA1A |
| EVPLL |
| EXOC3L4 |
| EXPH5 |
| EYA4 |
| F10 |
| F11 |
| F12 |
| F13B |
| F2 |
| F7 |
| F8 |
| F9 |
| FAAH |
| FABP1 |
| FABP3 |
| FADS2 |
| FADS6 |
| FAH |
| FAM107A |
| FAM110C |
| FAM124A |
| FAM129C |
| FAM134B |
| FAM135B |
| FAM13A |
| FAM149A |
| FAM150B |
| FAM151A |
| FAM163B |
| FAM169A |
| FAM180A |
| FAM198A |
| FAM211B |
| FAM213A |
| FAM227B |
| FAM229B |
| FAM26F |
| FAM3B |
| FAM46C |
| FAM65C |
| FAM69B |
| FAM71D |
| FAM83B |
| FAM83E |
| FAM9B |
| FANCC |
| FAT3 |
| FAXDC2 |
| FBN3 |
| FBP1 |
| FBXO15 |
| FBXO2 |
| FBXO40 |
| FBXO6 |
| FBXO7 |
| FCAMR |
| FCAR |
| FCER1A |
| FCER2 |
| FCGR2B |
| FCGR3A |
| FCGR3B |
| FCGRT |
| FCN1 |
| FCN2 |
| FCN3 |
| FCRL1 |

| |
|---|
| FCRLB |
| FDPS |
| FERMT2 |
| FETUB |
| FEZ1 |
| FFAR2 |
| FGD4 |
| FGF10 |
| FGF21 |
| FGF23 |
| FGFBP2 |
| FGFR2 |
| FGFR3 |
| FGFR4 |
| FGL2 |
| FGR |
| FHL1 |
| FHOD3 |
| FITM1 |
| FLRT1 |
| FMO3 |
| FMO4 |
| FMO5 |
| FNDC5 |
| FNIP2 |
| FOLH1 |
| FOS |
| FOSB |
| FOXA2 |
| FOXA3 |
| FOXD3 |
| FOXN2 |
| FOXN3 |
| FOXO1 |
| FPR1 |
| FPR2 |
| FRAS1 |
| FRAT1 |
| FREM1 |
| FREM2 |
| FRMD1 |
| FRMD4B |
| FRMD7 |
| FRMPD4 |
| FRRS1 |
| FTCD |
| FTCDNL1 |
| FUOM |
| FUT3 |
| FUT6 |
| FUZ |
| FXN |
| FXYD1 |
| FXYD3 |
| FXYD7 |
| FZD9 |
| G0S2 |
| GABRA2 |
| GABRG1 |
| GABRP |
| GADD45A |
| GADD45G |
| GAL3ST2 |
| GALK1 |
| GALM |
| GALNT14 |
| GALNT3 |
| GALT |
| GAMT |
| GAS2 |

| |
|---|
| GATA4 |
| GATA5 |
| GATM |
| GBP1 |
| GBP7 |
| GCAT |
| GCDH |
| GCGR |
| GCHFR |
| GCK |
| GCKR |
| GCOM1 |
| GCSAML |
| GDF2 |
| GDPD4 |
| GFI1B |
| GFRA1 |
| GFRA2 |
| GFRA3 |
| GGACT |
| GGT6 |
| GHR |
| GHRL |
| GIPC2 |
| GJB2 |
| GJB3 |
| GJC3 |
| GLIPR1L2 |
| GLOD5 |
| GLP2R |
| GLUD1 |
| GLUL |
| GLYAT |
| GLYATL1 |
| GLYATL3 |
| GLYCTK |
| GMPR |
| GNA14 |
| GNAO1 |
| GNAT1 |
| GNE |
| GNG5P2 |
| GNMT |
| GNPNAT1 |
| GNRH2 |
| GOLGA6A |
| GOLGA6B |
| GOLGA8M |
| GP1BA |
| GPAM |
| GPC6 |
| GPD1 |
| GPER1 |
| GPHN |
| GPLD1 |
| GPM6A |
| GPM6B |
| GPR123 |
| GPR125 |
| GPR126 |
| GPR128 |
| GPR142 |
| GPR143 |
| GPR158 |
| GPR162 |
| GPR182 |
| GPR75 |
| GPR82 |
| GPR83 |
| GPR88 |

| |
|---|
| GPR98 |
| GPRIN2 |
| GRAMD1C |
| GRAP |
| GREB1 |
| GREB1L |
| GRHL1 |
| GRHL2 |
| GRHL3 |
| GRHPR |
| GRIK3 |
| GRM8 |
| GRTP1 |
| GSDMB |
| GSTA1 |
| GSTA2 |
| GSTM5 |
| GSTT2 |
| GSTT2B |
| GSTZ1 |
| GYLTL1B |
| GYS2 |
| GZMM |
| H1FX |
| HAAO |
| HABP2 |
| HAGH |
| HAMP |
| HAO1 |
| HAO2 |
| HAP1 |
| HAS2 |
| HAUS4 |
| HBA1 |
| HBA2 |
| HBB |
| HBCBP |
| HBD |
| HBG2 |
| HCAR2 |
| HCAR3 |
| HDAC6 |
| HDC |
| HEMGN |
| HEPACAM |
| HERC5 |
| HEY2 |
| HFE2 |
| HGF |
| HGFAC |
| HHIP |
| HK3 |
| HLF |
| HLX |
| HMGCL |
| HMGCLL1 |
| HMGCR |
| HMGCS1 |
| HMGCS2 |
| HMOX1 |
| HNMT |
| HOGA1 |
| HOMER2 |
| HORMAD2 |
| HP |
| HPGDS |
| HPN |
| HPR |
| HPRT1 |
| HPX |

| |
|---|
| HRASLS2 |
| HRG |
| HS3ST3A1 |
| HS3ST3B1 |
| HS3ST4 |
| HSBP1L1 |
| HSD11B1 |
| HSD17B11 |
| HSD17B13 |
| HSD17B14 |
| HSD17B6 |
| HSD17B8 |
| HSD3B1 |
| HSD3B2 |
| HSDL2 |
| HSPB9 |
| HYDIN |
| ICAM4 |
| ID2 |
| IDNK |
| IDO2 |
| IFI27 |
| IFI44L |
| IFIT1 |
| IFIT1B |
| IFIT2 |
| IFIT3 |
| IFITM10 |
| IFNLR1 |
| IFT46 |
| IGF2 |
| IGFALS |
| IGFBP1 |
| IGFBP2 |
| IGFBP3 |
| IGJ |
| IGLON5 |
| IGSF10 |
| IGSF23 |
| IGSF9 |
| IHH |
| IL10 |
| IL11RA |
| IL13RA2 |
| IL17RB |
| IL17RC |
| IL17RE |
| IL1B |
| IL1RAP |
| IL1RN |
| IL20RA |
| IL22RA1 |
| IL23A |
| IL27 |
| IL33 |
| IMPA2 |
| INHBC |
| INMT |
| INS-IGF2 |
| INSIG2 |
| IQGAP2 |
| IQSEC3 |
| IRF6 |
| IRF7 |
| ISPD |
| ITCH |
| ITGA2B |
| ITGA9 |
| ITGAD |
| ITIH1 |

| |
|---|
| ITIH2 |
| ITIH3 |
| ITLN1 |
| ITPR2 |
| IYD |
| JAKMIP2 |
| JDP2 |
| JUN |
| JUNB |
| JUND |
| KANK4 |
| KAZN |
| KBTBD11 |
| KCNAB1 |
| KCND3 |
| KCNE1 |
| KCNH7 |
| KCNH8 |
| KCNJ10 |
| KCNJ13 |
| KCNJ15 |
| KCNJ3 |
| KCNK1 |
| KCNK17 |
| KCNK3 |
| KCNK5 |
| KCNMB2 |
| KCNN2 |
| KCTD16 |
| KDM8 |
| KHK |
| KIAA1161 |
| KIF12 |
| KIF17 |
| KIF19 |
| KIF1A |
| KIF25 |
| KIF26A |
| KIF6 |
| KIRREL3 |
| KLB |
| KLC4 |
| KLF10 |
| KLF11 |
| KLF12 |
| KLF4 |
| KLHL13 |
| KLHL33 |
| KLHL4 |
| KLKB1 |
| KLLN |
| KLRF1 |
| KNG1 |
| KRBOX1 |
| KRT1 |
| KRT19 |
| KRT73 |
| KRTAP5-9 |
| KRTCAP3 |
| L1CAM |
| LAG3 |
| LARGE |
| LBX2 |
| LCAT |
| LDLR |
| LDLRAD4 |
| LEAP2 |
| LEPREL1 |
| LGALS12 |
| LGALS4 |

| |
|---|
| LGI1 |
| LGI4 |
| LGSN |
| LHX2 |
| LIFR |
| LILRA1 |
| LILRA2 |
| LILRA5 |
| LILRA6 |
| LILRB1 |
| LILRB2 |
| LILRB3 |
| LILRB5 |
| LIMD2 |
| LIME1 |
| LINC00923 |
| LINGO4 |
| LIPA |
| LIPC |
| LIPG |
| LIPJ |
| LIPN |
| LNP1 |
| LONRF3 |
| LPA |
| LPIN2 |
| LRAT |
| LRFN5 |
| LRIG3 |
| LRP1B |
| LRP3 |
| LRRC16B |
| LRRC19 |
| LRRC25 |
| LRRC3 |
| LRRC31 |
| LRRC3DN |
| LRRC4 |
| LRRC55 |
| LRRFIP2 |
| LRRIQ1 |
| LRRK2 |
| LRRN1 |
| LRRN3 |
| LRRN4 |
| LRRTM1 |
| LRRTM2 |
| LRRTM4 |
| LSR |
| LSS |
| LST1 |
| LST3 |
| LTK |
| LY6E |
| LYNX1 |
| LYVE1 |
| LZTFL1 |
| MAATS1 |
| MACROD1 |
| MAFF |
| MAG |
| MAGEH1 |
| MAK |
| MAN1C1 |
| MAOA |
| MAOB |
| MAP1LC3A |
| MAP3K13 |
| MAPK4 |
| MARCO |

| |
|---|
| MARVELD3 |
| MASP1 |
| MASP2 |
| MAT1A |
| MATN2 |
| MBL2 |
| MBNL3 |
| MBOAT1 |
| MCHR1 |
| MDGA2 |
| MEFV |
| MEGF10 |
| MEI4 |
| MEIOB |
| MEP1B |
| MEST |
| METTL20 |
| METTL7A |
| MFAP3L |
| MFI2 |
| MFSD2A |
| MFSD4 |
| MGAM |
| MGMT |
| MGST2 |
| MLANA |
| MLF1 |
| MLIP |
| MLK4 |
| MMAB |
| MME |
| MMP19 |
| MMP25 |
| MMP7 |
| MMP8 |
| MMRN1 |
| MOGAT2 |
| MOGAT3 |
| MORC1 |
| MORC3 |
| MPDZ |
| MPPED1 |
| MPST |
| MRC1L1 |
| MREG |
| MRGPRF |
| MRO |
| MROH2A |
| MROH2B |
| MROH7 |
| MROH8 |
| MRPL23 |
| MRPS6 |
| MS4A2 |
| MS4A6A |
| MSMO1 |
| MST1 |
| MT1G |
| MT1H |
| MT1M |
| MTHFD1 |
| MTHFS |
| MTTP |
| MTUS2 |
| MUSK |
| MUT |
| MVK |
| MYCL |
| MYCT1 |
| MYH7B |

| |
|---|
| MYL3 |
| MYO15A |
| MYO16 |
| MYO1B |
| MYO1F |
| MYO3A |
| MYRIP |
| MYT1L |
| N4BP2L1 |
| NAAA |
| NAALAD2 |
| NAB2 |
| NAGS |
| NANOS1 |
| NAT1 |
| NAT2 |
| NAT8 |
| NCAM1 |
| NCF1 |
| NCKAP5 |
| NCMAP |
| NCR1 |
| NDNF |
| NDRG2 |
| NDST3 |
| NECAB2 |
| NEIL1 |
| NEU4 |
| NFAM1 |
| NFATC2 |
| NFE2 |
| NFIA |
| NFKBIA |
| NFKBIZ |
| NGF |
| NGFR |
| NGFRAP1 |
| NHSL1 |
| NIPAL1 |
| NIPSNAP1 |
| NLRC4 |
| NLRP11 |
| NLRP12 |
| NLRP14 |
| NLRP3 |
| NLRP6 |
| NMRK1 |
| NOL4 |
| NOS1 |
| NOS1AP |
| NOTUM |
| NOXO1 |
| NPAS3 |
| NPC1L1 |
| NPR1 |
| NPR2 |
| NR0B2 |
| NR1H3 |
| NR1I2 |
| NR1I3 |
| NR2F6 |
| NR5A2 |
| NRAP |
| NREP |
| NRG1 |
| NRG3 |
| NRG4 |
| NRN1 |
| NRXN1 |
| NSUN6 |

| |
|---|
| NSUN7 |
| NT5DC3 |
| NT5E |
| NTF3 |
| NTHL1 |
| NTN1 |
| NTN3 |
| NTN4 |
| NTRK1 |
| NUDT10 |
| NUDT4 |
| NUDT6 |
| NUGGC |
| NUP62CL |
| NXF3 |
| NYAP1 |
| OAF |
| OAS1 |
| OCLN |
| ODF3L1 |
| OGDHL |
| OIT3 |
| OLFM1 |
| OLFM4 |
| OLFML3 |
| OPRK1 |
| OR10J5 |
| OR13C3 |
| OR13C4 |
| OR13C5 |
| OR13C9 |
| OR2W3 |
| ORMDL3 |
| OSBPL6 |
| OSGIN1 |
| OTC |
| OTOA |
| OXER1 |
| OXT |
| P2RX3 |
| P2RX4 |
| P2RX6 |
| P2RY12 |
| P2RY13 |
| P2RY2 |
| P4HA1 |
| PACRG |
| PACSIN1 |
| PACSIN3 |
| PADI4 |
| PAGE5 |
| PAK7 |
| PALM3 |
| PAMR1 |
| PANK1 |
| PANX2 |
| PAQR7 |
| PAQR9 |
| PATZ1 |
| PBLD |
| PC |
| PCDH15 |
| PCDH20 |
| PCDH9 |
| PCDHAC1 |
| PCDHAC2 |
| PCK2 |
| PCLO |
| PCOLCE2 |
| PCP2 |

| |
|---|
| PCP4L1 |
| PCSK2 |
| PCSK9 |
| PCYT2 |
| PDCD1LG2 |
| PDE11A |
| PDE2A |
| PDE6G |
| PDE8B |
| PDIA5 |
| PDK2 |
| PDLIM2 |
| PDXP |
| PDZRN4 |
| PEBP1 |
| PEBP4 |
| PECR |
| PEG10 |
| PEG3 |
| PEMT |
| PER2 |
| PER3 |
| PEX11G |
| PFKFB1 |
| PGAM2 |
| PGAP3 |
| PGLYRP2 |
| PGM1 |
| PGM2 |
| PGM5 |
| PGRMC1 |
| PHACTR3 |
| PHGDH |
| PHLDA1 |
| PHLPP1 |
| PHOSPHO1 |
| PHYH |
| PHYHD1 |
| PHYHIPL |
| PI16 |
| PID1 |
| PIGR |
| PIK3AP1 |
| PIK3C2G |
| PIK3R1 |
| PIM1 |
| PIPOX |
| PITPNM3 |
| PKD2L1 |
| PKHD1L1 |
| PKLR |
| PKP2 |
| PLA1A |
| PLA2G12B |
| PLA2G16 |
| PLAC8 |
| PLCL2 |
| PLCXD2 |
| PLD5 |
| PLEK2 |
| PLEKHA4 |
| PLEKHA6 |
| PLEKHB1 |
| PLEKHF1 |
| PLEKHG6 |
| PLEKHG7 |
| PLG |
| PLGLB1 |
| PLGLB2 |
| PLIN1 |

| |
|---|
| PLIN2 |
| PLK3 |
| PLP1 |
| PLSCR4 |
| PM20D1 |
| PMEL |
| PMM1 |
| PMP2 |
| PNMA3 |
| PNMA6C |
| PNMAL2 |
| PNPLA3 |
| PNPLA7 |
| POFUT1 |
| POLR3GL |
| POMC |
| PON3 |
| POU6F2 |
| PPAP2B |
| PPARA |
| PPBP |
| PPFIBP2 |
| PPP1R15A |
| PPP1R1A |
| PPP1R3B |
| PPP1R3C |
| PPP4R4 |
| PPP6R2 |
| PQLC1 |
| PRAM1 |
| PRAP1 |
| PRELID2 |
| PRG4 |
| PRH2 |
| PRIMA1 |
| PRKAR2B |
| PRKCB |
| PRLR |
| PROC |
| PRODH2 |
| PROK2 |
| PROM1 |
| PROSER2 |
| PROZ |
| PRPSAP1 |
| PRR18 |
| PRR22 |
| PRR5 |
| PRRG4 |
| PRSS12 |
| PRSS22 |
| PRSS36 |
| PRSS42 |
| PRSS45 |
| PRSS50 |
| PRSS53 |
| PRSS8 |
| PSAT1 |
| PTCHD3 |
| PTCRA |
| PTGDR2 |
| PTGR1 |
| PTGS2 |
| PTH1R |
| PTK6 |
| PTMS |
| PTN |
| PTPRB |
| PTPRD |
| PTPRN2 |

| |
|---|
| PTPRS |
| PTPRT |
| PVALB |
| PVRL3 |
| PXMP2 |
| PYGL |
| PYROXD2 |
| PZP |
| QPRT |
| QRICH2 |
| QSOX1 |
| RAB17 |
| RAB25 |
| RAB26 |
| RAB27B |
| RAB39A |
| RAB3IL1 |
| RAD51AP2 |
| RAD54L2 |
| RAD9B |
| RAG1 |
| RALGPS2 |
| RALYL |
| RANBP3L |
| RARRES2 |
| RARRES3 |
| RASGEF1B |
| RASGRP2 |
| RASGRP4 |
| RASL10A |
| RASL10B |
| RASL11A |
| RASSF5 |
| RBP5 |
| RCAN1 |
| RCL1 |
| RD3L |
| RDH16 |
| RENBP |
| RET |
| RFNG |
| RFPL1 |
| RGL1 |
| RGN |
| RGPD2 |
| RGS18 |
| RGS3 |
| RGSL1 |
| RHBG |
| RHCE |
| RHOB |
| RIC3 |
| RIPK4 |
| RIPPLY1 |
| RIPPLY3 |
| RMDN2 |
| RMND5A |
| RND2 |
| RNF125 |
| RNF144B |
| RNF152 |
| RNF165 |
| ROBO2 |
| ROPN1L |
| RORC |
| ROS1 |
| RP11-10A14.4 |
| RP11-146D12.2 |
| RP11-181C3.1 |
| RP11-242G20.1 |

| |
|---|
| RP11-321F6.1 |
| RP11-422N16.3 |
| RP11-595B24.2 |
| RP11-650K20.3 |
| RP11-676J12.7 |
| RP11-766F14.2 |
| RP11-817J15.3 |
| RP11-867G23.8 |
| RP11-986E7.7 |
| RP11-998D10.1 |
| RPGRIP1 |
| RPS29 |
| RPS4Y1 |
| RPS6KA6 |
| RSAD2 |
| RSPH10B |
| RSPH4A |
| RSPO2 |
| RTN4RL1 |
| RTN4RL2 |
| RTP3 |
| RTP4 |
| RTTN |
| RUNDC3B |
| RXFP1 |
| RXRG |
| S100A1 |
| S100A12 |
| S100A14 |
| S100A8 |
| S100A9 |
| S1PR5 |
| SAA4 |
| SALL4 |
| SAMD4A |
| SAMD5 |
| SARDH |
| SAT2 |
| SATB1 |
| SC5D |
| SCG5 |
| SCGB3A1 |
| SCIMP |
| SCN11A |
| SCN2A |
| SCN3A |
| SCN7A |
| SCN9A |
| SCNN1B |
| SCNN1D |
| SCP2 |
| SCRN2 |
| SDC3 |
| SDK2 |
| SDPR |
| SEC14L2 |
| SEC14L3 |
| SEC14L4 |
| SELE |
| SELENBP1 |
| SELO |
| SEMA3D |
| SEMA3E |
| SEMA4A |
| SEMA4G |
| SEMA6A |
| SEMA6C |
| SEMA6D |
| SEPP1 |
| SERP2 |

| |
|---|
| SERPINA10 |
| SERPINA11 |
| SERPINA4 |
| SERPINA5 |
| SERPINA6 |
| SERPINA7 |
| SERPINC1 |
| SERPIND1 |
| SERPINE1 |
| SERPINF1 |
| SERPINF2 |
| SEZ6L |
| SFRP1 |
| SFRP5 |
| SFTPD |
| SFXN5 |
| SGCE |
| SGCZ |
| SH2D1B |
| SH2D4A |
| SH2D6 |
| SH3BP2 |
| SH3BP5 |
| SH3GL2 |
| SH3RF2 |
| SHB |
| SHBG |
| SHD |
| SHF |
| SHH |
| SHMT1 |
| SHROOM2 |
| SIGIRR |
| SIGLEC1 |
| SIGLEC11 |
| SIGLEC14 |
| SIGLEC15 |
| SIGLEC7 |
| SIGLEC9 |
| SIM1 |
| SIRPB1 |
| SIRT5 |
| SIVA1 |
| SKAP1 |
| SKIDA1 |
| SKOR1 |
| SLAIN1 |
| SLC10A1 |
| SLC13A5 |
| SLC15A1 |
| SLC16A1 |
| SLC16A10 |
| SLC16A12 |
| SLC16A2 |
| SLC16A4 |
| SLC16A9 |
| SLC17A2 |
| SLC17A8 |
| SLC18A2 |
| SLC19A1 |
| SLC19A2 |
| SLC19A3 |
| SLC22A1 |
| SLC22A10 |
| SLC22A24 |
| SLC22A25 |
| SLC22A7 |
| SLC22A9 |
| SLC23A1 |
| SLC24A2 |

| |
|---|
| SLC25A1 |
| SLC25A13 |
| SLC25A18 |
| SLC25A20 |
| SLC25A21 |
| SLC25A21-AS1 |
| SLC25A27 |
| SLC25A34 |
| SLC25A47 |
| SLC26A1 |
| SLC26A5 |
| SLC27A2 |
| SLC27A3 |
| SLC27A5 |
| SLC28A1 |
| SLC28A2 |
| SLC2A12 |
| SLC2A2 |
| SLC2A4RG |
| SLC2A9 |
| SLC30A1 |
| SLC30A10 |
| SLC31A2 |
| SLC34A1 |
| SLC35D1 |
| SLC37A4 |
| SLC38A11 |
| SLC38A4 |
| SLC39A4 |
| SLC39A5 |
| SLC3A1 |
| SLC43A3 |
| SLC45A3 |
| SLC47A1 |
| SLC4A1 |
| SLC51A |
| SLC52A1 |
| SLC5A1 |
| SLC5A7 |
| SLC5A9 |
| SLC6A1 |
| SLC6A12 |
| SLC6A13 |
| SLC6A19 |
| SLC6A20 |
| SLC6A4 |
| SLC7A8 |
| SLC7A9 |
| SLC8A1 |
| SLC9A3R2 |
| SLC9B2 |
| SLCO1A2 |
| SLCO1B3 |
| SLCO1B7 |
| SLCO2B1 |
| SLCO4C1 |
| SLITRK2 |
| SLITRK3 |
| SLITRK6 |
| SMAD6 |
| SMCO3 |
| SMIM1 |
| SMIM14 |
| SMIM19 |
| SMIM9 |
| SMLR1 |
| SMO |
| SMPD3 |
| SNCA |
| SNTB1 |

| |
|---|
| SNTG1 |
| SOAT2 |
| SOBP |
| SOCS2 |
| SOCS6 |
| SOD1 |
| SORCS1 |
| SORD |
| SORL1 |
| SOX10 |
| SOX5 |
| SPDYC |
| SPECC1L-ADORA2A |
| SPI1 |
| SPIB |
| SPIC |
| SPINT2 |
| SPOCK3 |
| SPP2 |
| SPSB3 |
| SPSB4 |
| SPTBN2 |
| SQLE |
| SRCIN1 |
| SRD5A1 |
| SRPX |
| SSTR1 |
| SSTR2 |
| ST14 |
| ST3GAL1 |
| ST3GAL6 |
| ST6GAL1 |
| ST6GALNAC2 |
| ST6GALNAC3 |
| ST8SIA3 |
| STAB1 |
| STAB2 |
| STAG3 |
| STARD10 |
| STARD4 |
| STEAP3 |
| STEAP4 |
| STMND1 |
| STPG2 |
| SUCNR1 |
| SULT1A1 |
| SULT1A2 |
| SULT1E1 |
| SULT2A1 |
| SUN2 |
| SUSD4 |
| SYBU |
| SYCE1 |
| SYDE2 |
| SYNE4 |
| SYNGR1 |
| SYT1 |
| SYT10 |
| SYT12 |
| SYT15 |
| SYT17 |
| SYT7 |
| SYTL4 |
| TACSTD2 |
| TADA1 |
| TAS1R3 |
| TBL1Y |
| TBX20 |
| TBXA2R |
| TBXAS1 |

| |
|---|
| TCEA3 |
| TCEAL2 |
| TCF21 |
| TCHH |
| TCL1A |
| TCP10L |
| TCP10L2 |
| TCTEX1D1 |
| TCTEX1D4 |
| TDRD10 |
| TDRD6 |
| TEF |
| TEK |
| TEKT2 |
| TEKT5 |
| TENM1 |
| TENM2 |
| TESK2 |
| TEX30 |
| TF |
| TFPI2 |
| TFR2 |
| TGFBR3 |
| THEMIS2 |
| THNSL1 |
| THOP1 |
| THRSP |
| TIAM1 |
| TIMD4 |
| TINAGL1 |
| TIPARP |
| TJP2 |
| TKTL1 |
| TLR4 |
| TM6SF2 |
| TMCO6 |
| TMEFF2 |
| TMEM105 |
| TMEM121 |
| TMEM125 |
| TMEM132C |
| TMEM132D |
| TMEM139 |
| TMEM150C |
| TMEM170B |
| TMEM176B |
| TMEM200B |
| TMEM200C |
| TMEM220 |
| TMEM232 |
| TMEM25 |
| TMEM252 |
| TMEM26 |
| TMEM27 |
| TMEM30B |
| TMEM37 |
| TMEM45B |
| TMEM47 |
| TMEM52 |
| TMEM56 |
| TMEM63C |
| TMEM71 |
| TMEM82 |
| TMEM86B |
| TMEM97 |
| TMIE |
| TMPO |
| TMPRSS2 |
| TMPRSS4 |
| TMPRSS6 |

| |
|---|
| TMPRSS9 |
| TMSB4Y |
| TNF |
| TNFAIP8L1 |
| TNFRSF11B |
| TNFSF10 |
| TNFSF11 |
| TNN |
| TNNC1 |
| TNR |
| TOM1L1 |
| TOX2 |
| TP53I13 |
| TP53INP1 |
| TPH2 |
| TPPP2 |
| TPRG1 |
| TPSAB1 |
| TPST2 |
| TRABD2B |
| TRAPPC3L |
| TRDN |
| TREH |
| TREM1 |
| TREML2 |
| TRHDE |
| TRIB1 |
| TRIM58 |
| TRIM63 |
| TRPC5 |
| TRPM6 |
| TRPM8 |
| TRPV4 |
| TRPV6 |
| TSHR |
| TSLP |
| TSPAN11 |
| TSPAN12 |
| TSPAN7 |
| TSPAN9 |
| TST |
| TSTD1 |
| TTBK1 |
| TTC36 |
| TTC38 |
| TTC40 |
| TTC7B |
| TTPA |
| TTR |
| TUBB1 |
| TUBE1 |
| TUSC1 |
| TXNDC16 |
| TXNIP |
| TXNRD2 |
| UAP1 |
| UGP2 |
| UGT1A1 |
| UGT2B10 |
| UGT2B15 |
| UGT2B17 |
| UGT2B7 |
| UGT3A1 |
| UNC13D |
| UNC79 |
| UNC93A |
| UPB1 |
| UROC1 |
| USH2A |
| USP18 |

| |
|---|
| USP44 |
| USP51 |
| UTY |
| VCAM1 |
| VEPH1 |
| VIL1 |
| VIPR1 |
| VIPR2 |
| VMO1 |
| VNN1 |
| VNN2 |
| VNN3 |
| VPS37B |
| VPS37D |
| VSIG4 |
| VSNL1 |
| VTCN1 |
| VWA3B |
| VWCE |
| VWDE |
| WBSCR27 |
| WDR17 |
| WDR72 |
| WNK2 |
| WNK3 |
| WNT11 |
| WNT5A |
| WNT5B |
| WNT7A |
| XAGE3 |
| XDH |
| XKR4 |
| XPNPEP2 |
| XRCC6BP1 |
| XYLB |
| YBX2 |
| YPEL1 |
| YPEL2 |
| ZBTB18 |
| ZC3H12C |
| ZCCHC6 |
| ZCWPW1 |
| ZDHHC19 |
| ZFP1 |
| ZFY |
| ZG16 |
| ZIC1 |
| ZMYND12 |
| ZNF175 |
| ZNF268 |
| ZNF311 |
| ZNF334 |
| ZNF354C |
| ZNF358 |
| ZNF367 |
| ZNF385B |
| ZNF385C |
| ZNF470 |
| ZNF471 |
| ZNF502 |
| ZNF511 |
| ZNF536 |
| ZNF572 |
| ZNF577 |
| ZNF648 |
| ZNF662 |
| ZNF676 |
| ZNF682 |
| ZNF812 |
| ZNF879 |

| |
|---|
| ZPBP |
| ZPLD1 |
| ZSCAN18 |
| ZYG11A |

**Supplementary Figure S1. (A) *DNAJB1-PRKACA* gene fusion. (B) *DNAJB1-PRKACA* fusion results in upregulation of *PRKACA* expression.**

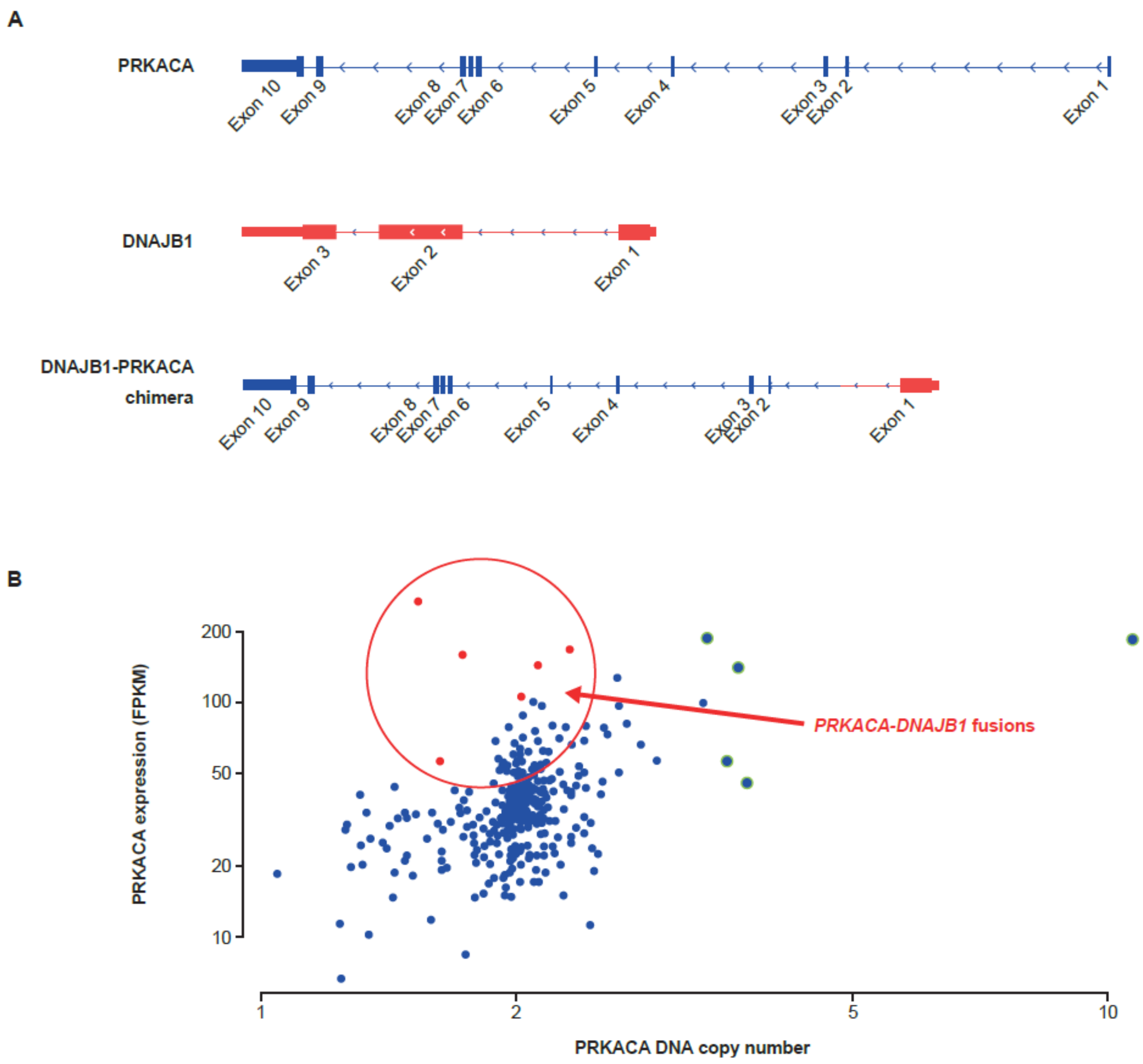

**Supplementary Figure S2. (A–B) BLU0588 dose-dependently decreases VASP phosphorylation in FCL PDX spheroid cultures (3 biological replicates).**

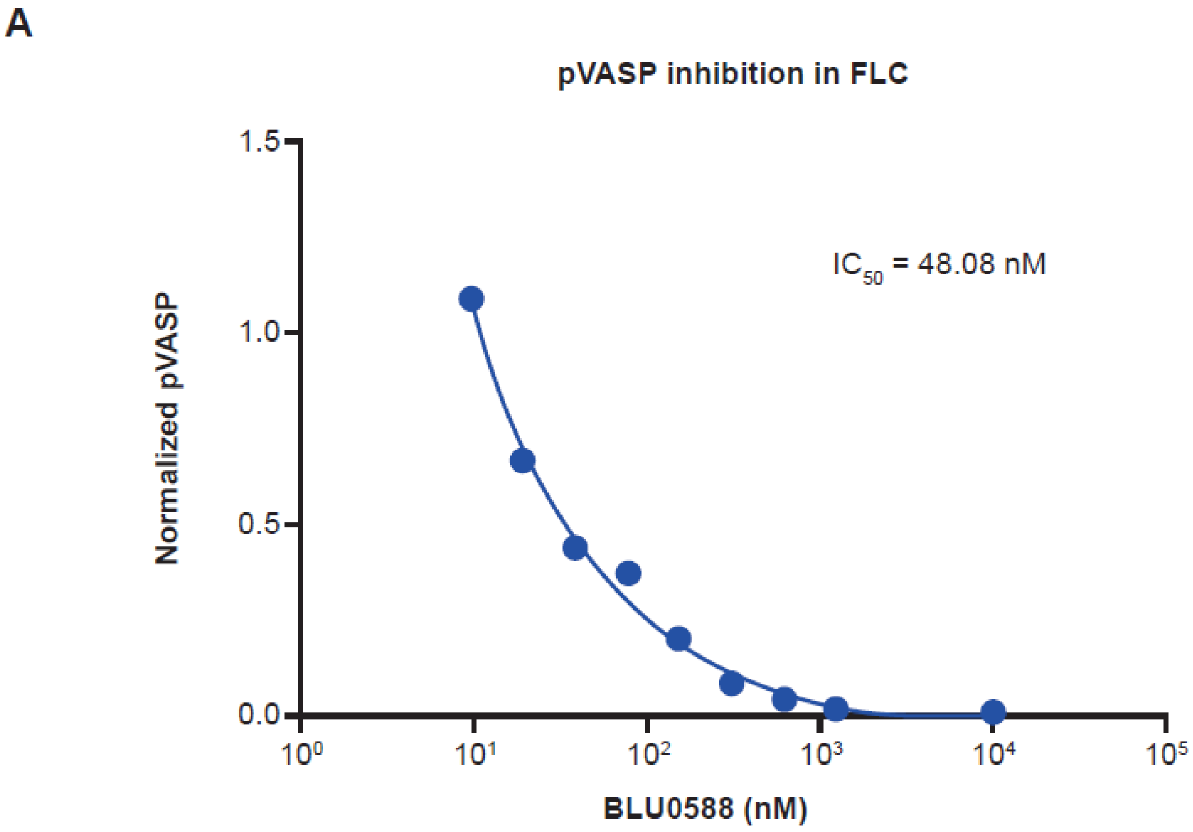

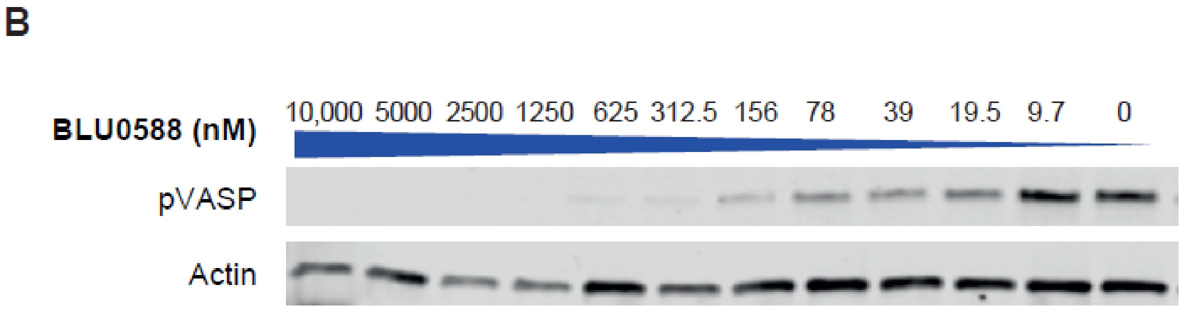

**Supplementary Figure S3. PRKACA transcript level reads by shRNA in FLC cell lines.**

Wilcoxon rank test *P* = 0.022 for all doxycycline vs no doxycycline.

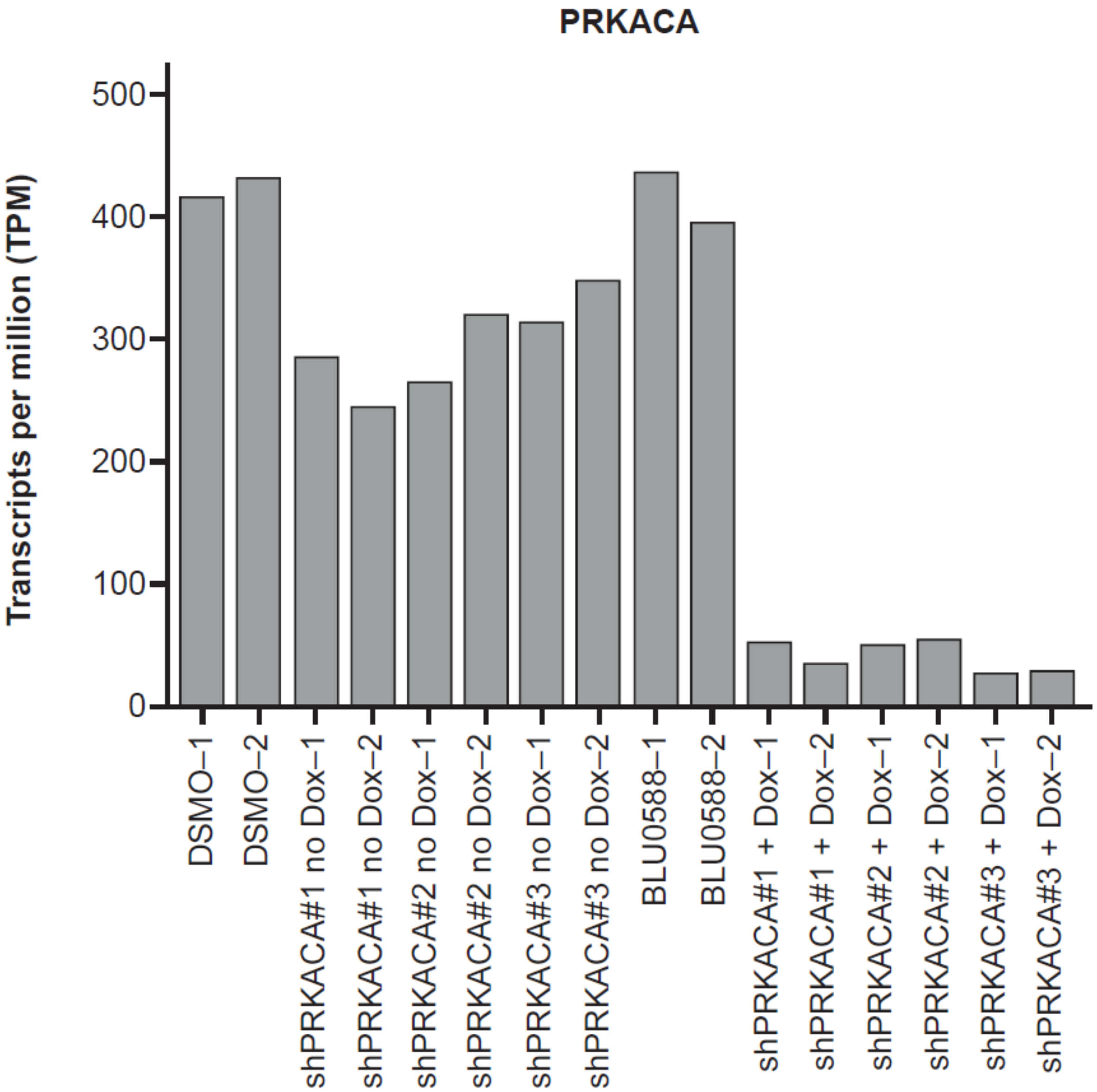

**Supplementary Figure S4. Overlap of consensus between PRKACA shRNA and published FLC gene signature(14).**

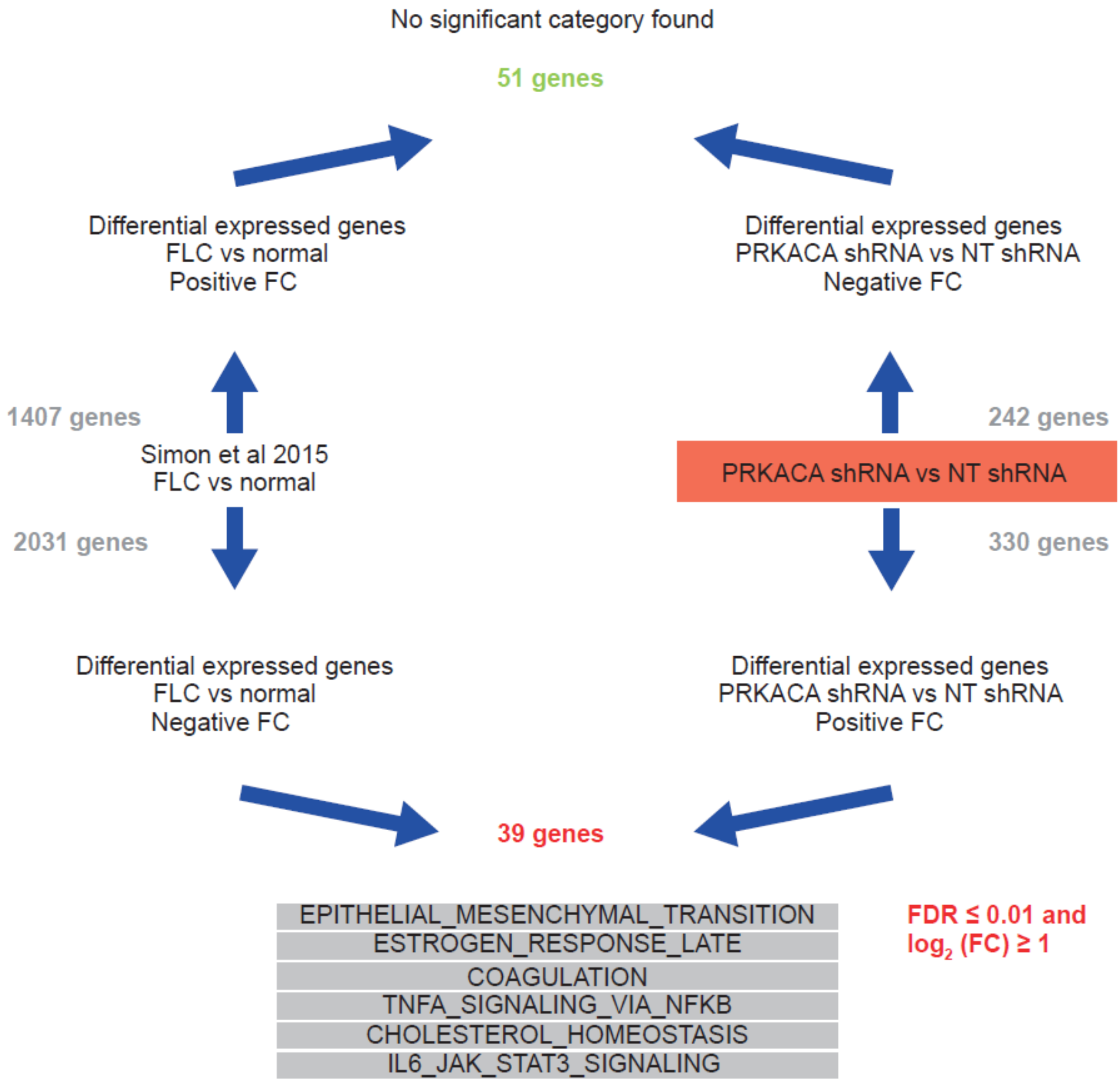

**Supplementary Figure S5. (A) Overlap of consensus shRNA with BLU0588 modulated gene expression. (B) Overlap of PRKACA inhibition signature with**

**FLC gene signature.**

A

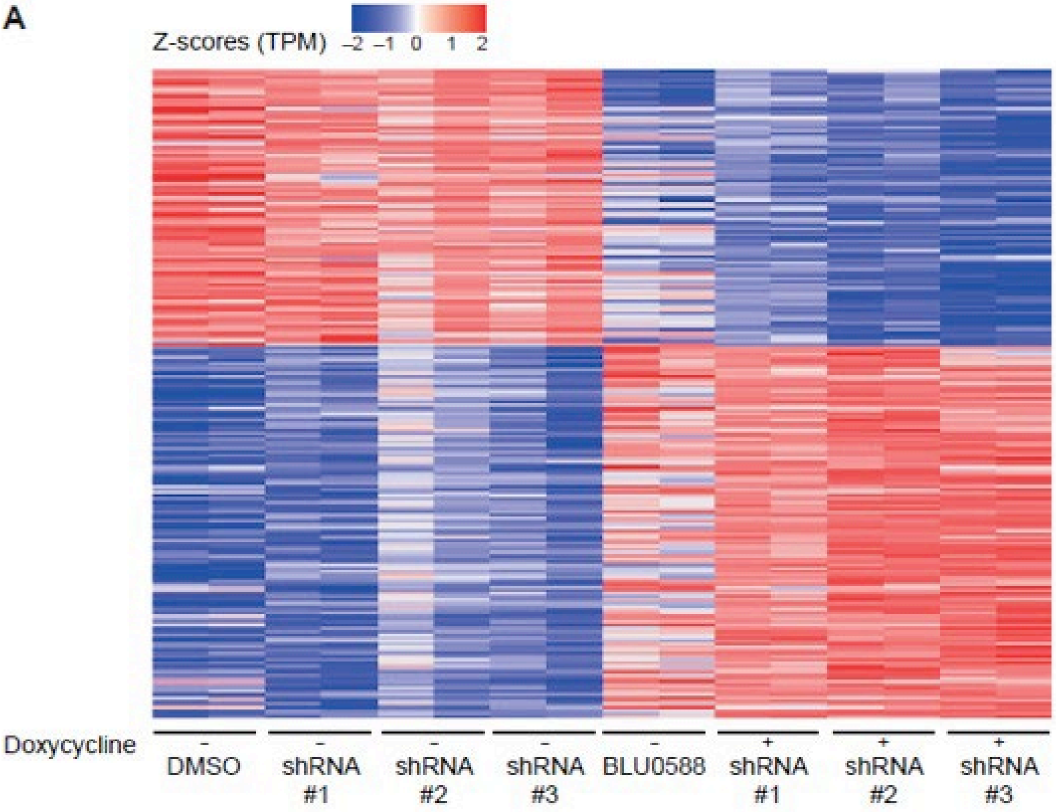

B

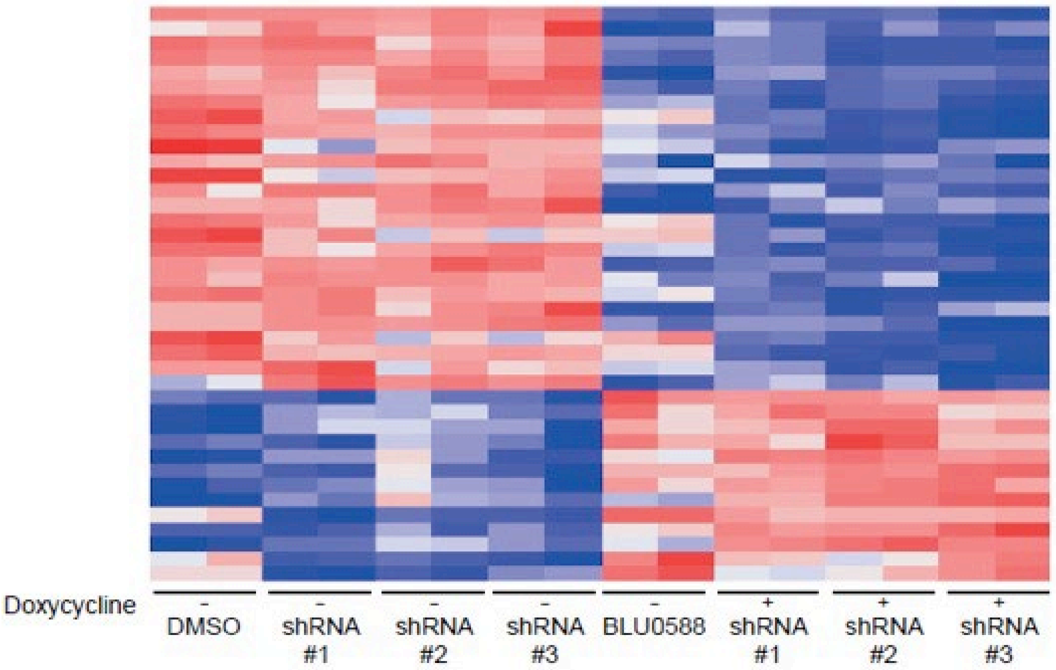

**Supplementary Figure S6. Transcription factor binding and network analysis.**

(A) Transcription factor motifs enriched in genes modulated by BLU0588, PRKACA shRNA knockdown, and the combined signature of PRKACA inhibition. (B) Clarivate Metacore analysis established networks connecting the differentially expressed genes and associated transcription factor motifs.

A

| Categories | Genes |
|---|---|
| 39 common diff genes (Simon, shRNA, BLU0588) | ADAMTSL2 CAPN5 CYP2S1 EBF4 ECM1 IGSF9 LIMD2 NGFR RGL1 SHH TP53INP1 VPS37D YPEL1 AKR1B15 ATP1B1 CABYR CPLX2 CPS1 DDIT4 EVA1C FSTL4 GK GPRIN3 HPD KCNU1 KRT86 KYNU MUC13 PBK PDE3A PDE4B PPARGC1A RHOBTB1 S100P SDCBP2 SLC16A11 SLC22A11 SLC5A6 TESC |
| HOMER TF motifs | HNF4A HNF1A HNF1B CUX2 HOXB13 JUNB ATF3 ESRRA FOXF1 NF1 FOS CEBPA CEBPB CEBPZ |
| Central target | PRKACA |

B

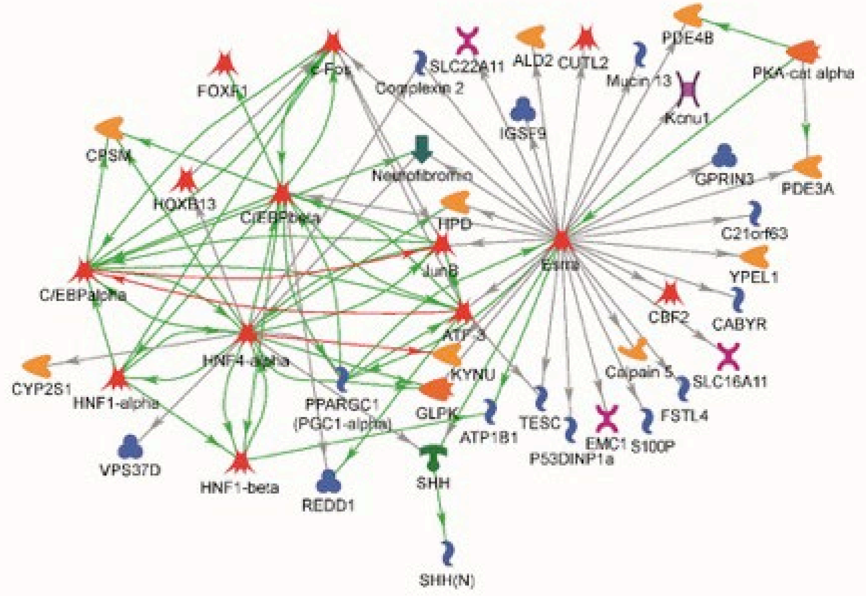

**Supplementary Figure S7.** BLU0588 dose-dependently reduces expression of genes normally upregulated in FLC (A) and dose-dependently increases expression of genes that are normally downregulated in FLC (B).

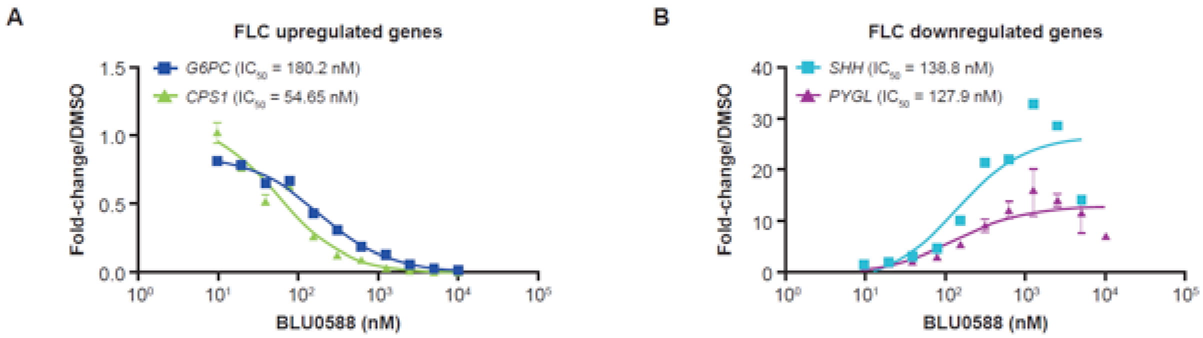
